# Functional determinants of a small protein controlling a broadly conserved bacterial sensor kinase

**DOI:** 10.1101/438515

**Authors:** Srujana S Yadavalli, Ted Goh, Jeffrey N Carey, Gabriele Malengo, Sangeevan Vellappan, Bryce E Nickels, Victor Sourjik, Mark Goulian, Jing Yuan

## Abstract

The PhoQ/PhoP two-component system plays a vital role in the regulation of Mg^2+^ homeostasis, resistance to acid and hyperosmotic stress, cationic antimicrobial peptides, and virulence in *Escherichia coli*, Salmonella and related bacteria. Previous studies have shown that MgrB, a 47 amino acid membrane protein that is part of the PhoQ/PhoP regulon, inhibits the histidine kinase PhoQ. MgrB is part of a negative feedback loop modulating this two-component system that prevents hyperactivation of PhoQ and may also provide an entry point for additional input signals for the PhoQ/PhoP pathway. To explore the mechanism of action of MgrB, we have analyzed the effects of point mutations, C-terminal truncations and transmembrane region swaps on MgrB activity. In contrast with two other known membrane protein regulators of histidine kinases in *E. coli*, we find that the MgrB TM region is necessary for PhoQ inhibition. Our results indicate that the TM region mediates interactions with PhoQ and that W20 is a key residue for PhoQ/MgrB complex formation. Additionally, mutations of the MgrB cytosolic region suggest that the two N-terminal lysines play an important role in regulating PhoQ activity. Alanine scanning mutagenesis of the periplasmic region of MgrB further indicates that, with the exception of a few highly conserved residues, most residues are not essential for MgrB’s function as a PhoQ inhibitor. Our results indicate that the regulatory function of the small protein MgrB depends on distinct contributions from multiple residues spread across the protein. Interestingly, the TM region also appears to interact with other non-cognate histidine kinases in a bacterial two-hybrid assay, suggesting a potential route for evolving new small protein modulators of histidine kinases.

## Introduction

Small proteins—proteins less than 50-amino acids long that are directly translated from small open reading frames—regulate a wide variety of cellular processes but, collectively, this class of proteins remains relatively understudied (reviewed in [1, 2]). For example, hundreds to thousands of small open reading frames have been reported in *Escherichia coli* [3–5], *Mycobacterium tuberculosis* [6] and human microbiomes [7]. A fraction of them have been confirmed with detections at the protein level [3–6] and the functions of most these proteins are unknown. In bacteria and bacteriophages, small proteins can be classified into two groups: secreted and non-secreted [1]. The former are mainly involved in communication or competition among organisms, whereas the latter are localized in the cytoplasm or cell membrane and play a role in diverse aspects of cell physiology. In *E. coli*, the list of small proteins that have been identified and verified continues to grow with improvements in computational and experimental techniques [3, 4, 8]. Interestingly, nearly one third have been shown or are predicted to be in the cytoplasmic membrane, suggesting many may be involved in sensing environmental cues and mediating stress responses.

Bacteria thrive in a multitude of niches, often under challenging physicochemical conditions, and have therefore evolved stress response systems to monitor the environment and modulate their physiology accordingly. Many of these stress responses are controlled by two-component signaling systems, one of the primary modes of bacterial signal transduction [9]. The PhoQ/PhoP system in *E. coli*, Salmonella, and related Gram-negative bacteria is a well-studied example of a two-component system that is critical for virulence and facilitates adaptation to conditions of low Mg^2+^ or Ca^2+^, low pH, osmotic upshifts and the presence of cationic antimicrobial peptides [10–14]. In response to input signal, the sensor histidine kinase PhoQ autophosphorylates and modulates the phosphorylation state of the response regulator PhoP, which functions as a transcription factor [10, 15]. At least three small proteins are regulated by PhoP: the 31-amino acid membrane protein MgtS is induced under magnesium limitation in *E. coli* and inhibits degradation of the magnesium transporter MgtA [16]; the 30-amino acid protein MgtR promotes the degradation of the virulence factor MgtC by the FtsH protease in *Salmonella enterica* serovar Typhimurium [17]; and the 47-amino acid protein MgrB directly inhibits PhoQ kinase activity, thereby mediating a negative feedback loop [18, 19]. All of these small proteins are membrane-localized, regulate proteins expressed in the PhoP regulon, and modify cell responses to the environment.

MgrB orthologs are found in numerous Enterobacteriaceae, and PhoQ inhibition has been demonstrated explicitly for a few distinct genera. The physiological importance of this inhibition and the associated negative feedback loop remain poorly understood. MgrB could serve as a point of control for additional input signals that modulate PhoQ. For example, MgrB activity is affected by the redox state of the periplasm via the protein’s two conserved periplasmic cysteines [20]. MgrB is not essential and appears to mainly act in a regulatory role, however it can have a significant impact on bacterial physiology. In *E. coli*, recent work has shown that strong stimulation of the PhoQ/PhoP system hinders cell division [21]. Cells grown in low levels of Mg^2+^ in the absence of MgrB form filaments due to strong activation of PhoQ in this condition. MgrB-mediated feedback inhibition thus appears to be important to appropriately limit PhoQ activity and prevent hyperactivation of the system. In at least some contexts, loss of *mgrB* can also confer a fitness advantage: mutation of *mgrB* has emerged as one of the primary mechanisms of acquired resistance to the last resort antibiotic colistin in clinical isolates of *Klebsiella pneumoniae* [22–25].

In addition to MgrB, the PhoQ histidine kinase is modulated by another membrane protein, SafA [26], which is present exclusively in *E. coli*, and acts as a connector between the EvgS/EvgA and PhoQ/PhoP two-component systems. Stimulation of EvgS by low pH upregulates SafA expression [27], which activates PhoQ kinase activity and leads to increased acid resistance. In at least some *E. coli* isolates, this connection is required for PhoQ activation in response to acid stress [28, 29]. Another example of a connector protein that modulates the activity of a histidine kinase is MzrA [30], which links the CpxA/CpxR and EnvZ/OmpR two-component systems. Although not as small as MgrB, both SafA and MzrA are relatively small for membrane proteins (65 and 127 amino acids, respectively) and, like MgrB, are bitopic with their N-termini in the cytoplasm.

MgrB acts by inhibiting PhoQ kinase activity [19], however, the mechanism by which this small protein inhibits a protein that is roughly ten times larger than itself, and the sequence determinants responsible for this action are largely unknown. In this work, we show that the transmembrane (TM) region of MgrB is required for the protein to form a complex with its target kinase PhoQ, in contrast with SafA and MzrA, which do not require their TM regions for their activity [31, 32]. Using site-directed mutagenesis combined with MgrB functional assays and *in vivo* Förster resonance energy transfer (FRET), we identified W20 in the MgrB TM region to be crucial for MgrB/PhoQ complex formation. Additionally, we identified 11 residues across different regions of MgrB that are important for its inhibitory function. Among these residues, two lysines in the cytoplasmic region likely act through their charged side chains to regulate PhoQ HAMP conformation, although they do not appear to affect MgrB/PhoQ complex formation. Surprisingly, a large number of amino acids in the small periplasmic region of MgrB can be replaced with alanine with only minor effects on protein function. Together our findings suggest that PhoQ inhibition by MgrB depends on coordinated interactions from amino acid residues spanning different regions of the protein. We also uncovered interactions of MgrB with other histidine kinases in bacterial two-hybrid assays, which may provide a starting point for evolving novel small membrane proteins that modulate other histidine kinases.

## Results

### Sequence analysis of MgrB

MgrB was shown to be a membrane-localized small protein with N-terminal-in and C-terminal-out topology [18]. To predict the transmembrane (TM) region in MgrB, we used eight algorithms (PHDHTM, DAS-TMFILTER, SCAMPI, TMHMM, TMpred, TMMTOP, PRED-TMR and MEMSAT) to analyze the entire amino acid sequence (Fig 1). The predictions were fairly convergent on the TM cytoplasm/membrane boundary, which was located at W6 or V7. In contrast, the predictions of the membrane/periplasm boundary were quite different among these programs, ranging from F24 to M27. In this manuscript, we use W6 to F24 as the predicted transmembrane region of MgrB.

**Fig 1.**
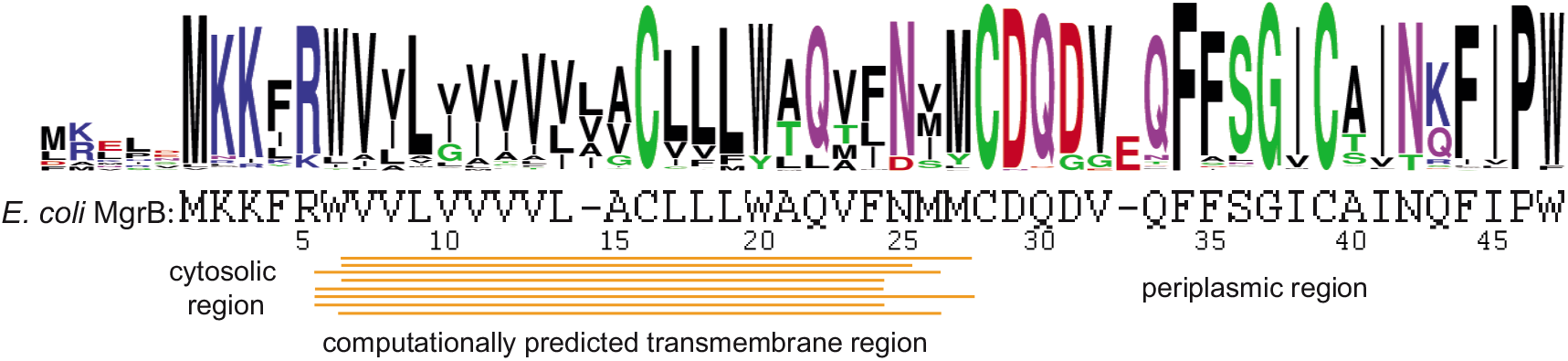
The conservation of MgrB sequence. The sequence logo of MgrB generated with bacterial MgrB sequences obtained from KEGG protein BLAST using *E. coli* MgrB as query highlights the degree of amino acid conservation. The MgrB sequences and multiple sequence alignment can be found in the supplementary file (MgrB_alignment).

To gain insight on potential functionally important residues in MgrB, we aligned over 100 bacterial MgrB sequences obtained from KEGG protein BLAST using *E. coli* MgrB as query (Fig 1 and supplementary file 1). As indicated in the sequence logo of the alignment, the amino acids were fairly conserved in the cytosolic and periplasmic regions. This high degree of conservation might originate from limitations in the diversity of MgrB sequences currently available in databases or suggest the functional importance of each conserved amino acid in these regions. In contrast, more variations and insertion of one amino acid were shown in the predicted TM region. Interestingly, two amino acids with polar side chains (C16 and Q22) and two tryptophans (W6 and W20) in the TM region are highly conserved, suggesting a functional importance.

### MgrB TM and periplasmic regions are necessary but not sufficient for MgrB function

As a first step towards understanding how MgrB inhibits PhoQ at a structural level, we looked at the role of the protein’s TM and periplasmic regions. To test the functional role of the MgrB TM region, we swapped the predicted MgrB TM region with either (i) a 20-amino acid TM from the *E. coli* maltose transporter MalF or (ii) a 21-amino acid variant of human glycophorin A (GpA) TM [33] (Fig 2A). The MgrB variants described in this section are derivatives of a functional N-terminally GFP tagged MgrB (GFP-MgrB) [18]. Both MgrB variants with swapped TM regions localized to the cell membrane (S1 Fig) and their expression levels were comparable to that of wild type (S2 Fig). TM swapping experiments of two previously studied histidine kinase regulators–SafA and MzrA—indicate that the transmembrane sequence is dispensable for their function [31, 32]. Indeed, our control experiments in which we replaced the TM sequences of SafA and MzrA with that of GpA confirmed these results (S3 Fig). Thus, for SafA and MzrA, the TM region appears to primarily act as an anchor to direct the regulatory domain of these proteins to the membrane. We monitored the effect of MgrB and its TM swap variants on PhoQ activity using a PhoQ/PhoP-regulated transcriptional reporter (P*_mgtA_-lacZ*) (Fig 2A). Control cells expressing no MgrB had a high level of *β*-galactosidase (*β*-gal) activity, indicating uninhibited PhoQ, whereas cells expressing WT MgrB had minimal *β*-gal activity, indicating PhoQ inhibition by WT MgrB as expected. In contrast to the inhibitory function of the WT MgrB, both MgrB TM swap variants (MgrB-MalF TM and MgrB-GpA TM) showed high *β*-gal activities that were about 80% of the activity of no-MgrB control, indicating that they are unable to inhibit PhoQ efficiently. These results suggest that the MgrB TM region not only functions as a membrane anchor but also has additional properties that are critical for MgrB to inhibit PhoQ.

**Fig 2.**
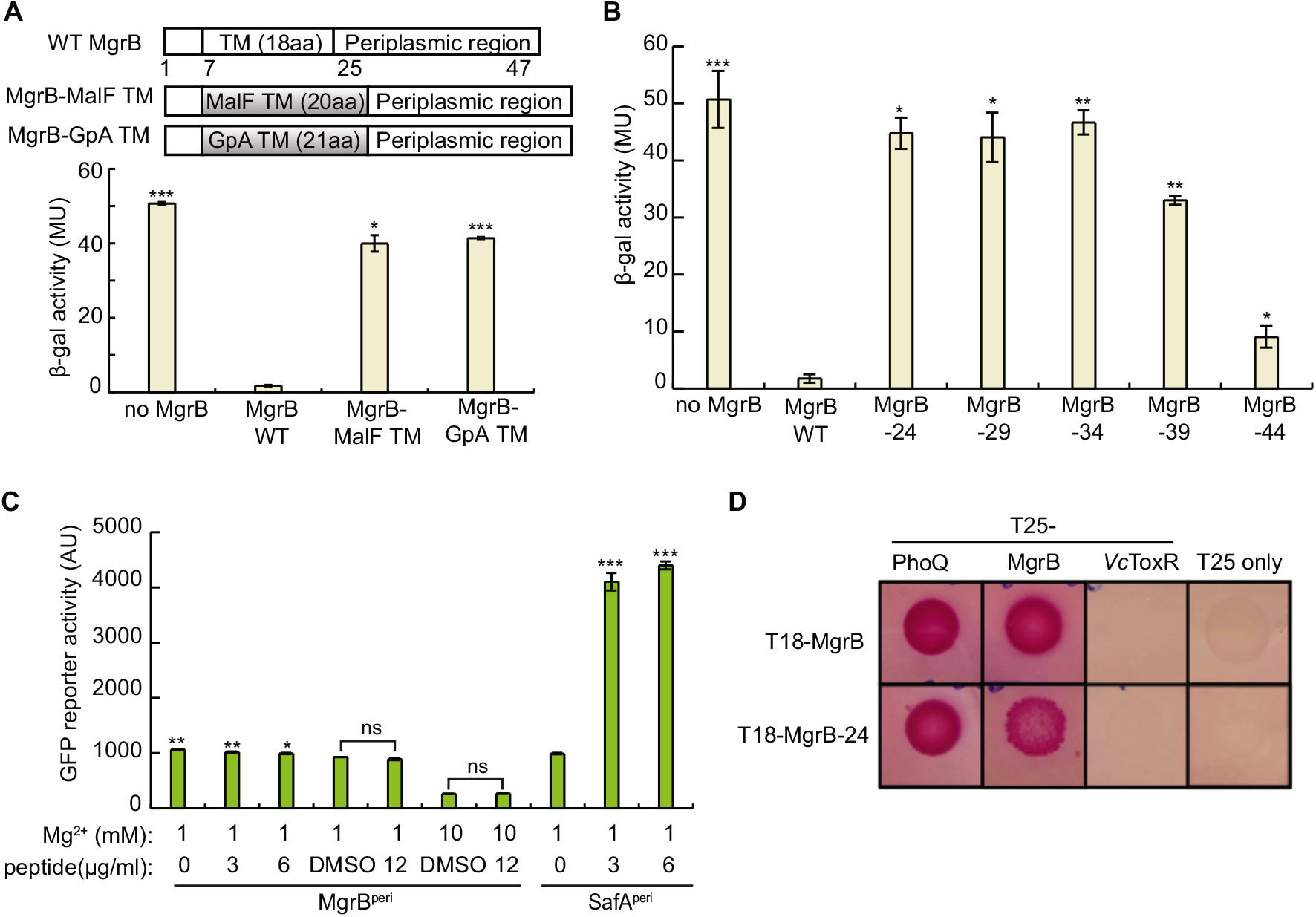
The MgrB transmembrane domain and periplasmic domain are essential but not sufficient for its function as a PhoQ inhibitor. **A**, Schematic representation of transmembrane (TM) domain swap constructs to replace the MgrB TM sequence with TM sequences derived from either the protein MalF or human glycophorin A, GpA. *β*-galactosidase activities of wild-type MgrB (pAL38), its variants MgrB-MalF TM (pSY3), MgrB-GpA TM (pSY4) and no MgrB control (pAL39) in a Δ*mgrB* strain containing a PhoQ/PhoP-regulated transcriptional reporter P*_mgtA_-lacZ* (AML67) is shown below. Inducer was not added as the leaky expression from the *trc* promoter resulted in sufficient levels for strong inhibition by WT MgrB. Data represent averages and ranges for two independent experiments. MU = Miller Units. **B**, effect of wild-type MgrB and the truncation mutants were measured in a Δ*mgrB* strain containing the PhoQ/PhoP-regulated transcriptional reporter P*mgtA-lacZ* (AML67) as in **A**. **C**, the activities of MgrB^Peri^ and SafA^Peri^ were monitored using PhoQ/PhoP-regulated transcriptional reporter P*_mgtLA_-gfp*. Peptides or DMSO were added to the *E. coli imp4213* culture with indicated amount. The GFP activities were measured after 2h incubation using flow cytometry. Data points represent averages of three independent experiments and error bars show SDs. **D**, bacterial two-hybrid (BACTH) colorimetric spot assays with cells expressing protein fusions to the T18 and T25 subunits of *B. pertussis* adenylyl cyclase. Representative images showing interactions between MgrB-24 (T18-MgrB-24) and partner proteins PhoQ (T25-PhoQ) and MgrB (T25-MgrB). Interactions between MgrB-PhoQ, MgrB-MgrB, MgrB-T25 only and (MgrB-24)-T25 only were included as positive and negative controls. As an additional control, interactions were tested between MgrB-24 and *Vibrio cholerae* membrane protein ToxR (*Vc*ToxR), and also between MgrB and *Vc*ToxR. Each pair of interactions was assayed in at least three independent trials in a *cyaA^-^* BACTH host strain containing *lacl*^q^ (SAM85), on maltose MacConkey indicator plates (results from one representative experiment are shown). The plates were incubated at 30°C for 48 h before images of plates were acquired. The pink color indicates reconstitution of adenylyl cyclase activity. The statistical significances of changes in reporter activity were calculated with *t* tests by comparing to wild type (in A, B) or to the DMSO negative control (in C) and indicated with asterisks (*** *P* ≤ 0.001, ** *P* ≤ 0.01, * *P* ≤ 0.05, ns = not significant) in A, B and C.

As swapping the TM sequence in MgrB severely affected its activity, we tested whether an MgrB mutant lacking the periplasmic region inhibits PhoQ. This variant—MgrB-24, in which N25 is replaced with a stop codon—showed proper membrane localization (S1 Fig), but could not repress reporter gene expression (Fig 2B), indicating that this truncated variant composed of the MgrB TM and short N-terminal cytoplasmic regions is not functional. To assess the minimum length of MgrB needed for efficient repression of PhoQ/PhoP-regulated reporter gene expression, we tested other C-terminal truncations of MgrB: MgrB-29, MgrB-34, MgrB-39, MgrB-44, corresponding to replacing the codons for Q30, F35, A40 and I45 with a stop codon, respectively. These mutants displayed similar membrane localization to that of WT MgrB (S1 Fig). Of these four truncations, MgrB-29 and −34 did not show any detectable repression, MgrB-39 showed slight repression, and MgrB-44, which lacks only the last two residues, showed the strongest repression, although last variant was still compromised in activity compared with WT (Fig 2B). Evidently, amino acid interactions in both the TM and the periplasmic region of MgrB influence the protein’s ability to inhibit PhoQ.

Previous work suggested that the periplasmic region of MgrB is required for PhoQ inhibition [18]. Since the TM swaps described above change the TM helix length, their loss of activity could be due to a change in the helical phase at the C-terminus of the TM helix creating different orientations of the periplasmic region. It is therefore possible that the periplasmic region of MgrB alone could inhibit PhoQ, while the TM helix only ensures its correct orientation. Indeed, the chemically synthesized periplasmic region of SafA (SafA^peri^) was previously shown to be sufficient for PhoQ activation when added exogenously [34]. Therefore, we tested whether the MgrB periplasmic region (MgrB^peri^) could function similarly to full-length MgrB (Fig 2C) by adding MgrB^peri^ to an *E. coli* culture. To enable the MgrB^peri^ peptide to access the periplasm, we used an *imp-4213* strain, which has increased outer membrane permeability [35]. We monitored PhoQ activity with a GFP reporter fused to the promoter region of a PhoQ/PhoP regulated gene (pUA66 P*_mgtLA_-gfp*) in the presence or absence of chemically synthesized MgrB^peri^. The positive control, chemically synthesized SafA^peri^, enhanced PhoQ activity significantly, consistent with the previous report [34]. In contrast, MgrB^peri^ did not show any inhibitory effect on PhoQ activity in the concentration range that we tested, though DMSO the solvent for MgrB^peri^ peptide showed a slight inhibition of *gfp* reporter expression. As MgrB^peri^ is short and likely capable of sampling different conformations, our results suggest that, unlike SafA, the MgrB periplasmic region alone is not sufficient to inhibit PhoQ. Taken together, the above results indicate that both the MgrB TM and periplasmic regions are required but not sufficient to inhibit PhoQ.

### MgrB lacking its periplasmic region is capable of physical interaction with PhoQ

Although MgrB-24 (N-terminal region+TM region) does not inhibit PhoQ, it could still play a role in establishing physical contact with PhoQ. To explore this possibility, we used a bacterial two-hybrid (BACTH) system based on split adenylyl cyclase [36, 37]. In the assay, reconstitution of the T18 and T25 fragments restores the activity of adenylate cyclase (CyaA), which leads to an increase in cAMP levels. This increased cAMP in turn enables efficient induction of the genes required for maltose catabolism, resulting in pink colonies on maltose MacConkey plates. To assay interactions of MgrB lacking its periplasmic region with PhoQ, we prepared N-terminal fusions of MgrB-24 and PhoQ to T18 and T25 fragments, respectively. As controls, we tested interactions between MgrB–PhoQ, MgrB–*Vc*ToxR (*Vibrio cholerae* inner membrane protein ToxR) as well as MgrB–T25 fragment alone. Spot assays of cells expressing T18-MgrB-24 and T25-PhoQ produced a dark pink color after 48 h at 30 °C, whereas cells expressing T18-MgrB-24 and the T25 fragment alone were colorless (Fig 2D). Spots of cells expressing T18-MgrB-24 and either T25-MgrB or T25-MgrB-24 were also pink. Control cells expressing T18-MgrB and either T25-PhoQ or T25-MgrB formed dark pink spots, showing an interaction of MgrB with PhoQ and possibly with itself, consistent with previous work [18], whereas control cells expressing either T18-MgrB or T18-MgrB-24 with either T25-*Vc*ToxR or T25 fragment alone remained colorless. Taken together, these results suggest that the MgrB N-terminus+TM region interacts with PhoQ and may also interact with itself to facilitate formation of a multimeric complex.

### A small number of functionally important residues are spread across different regions of MgrB

To explore the relative importance of different residues in MgrB, we made alanine substitutions and analyzed their effects on PhoQ inhibition. Interestingly, a majority of the substitutions in MgrB did not affect PhoQ/PhoP reporter gene expression, but 11 substitutions spread across different regions showed significant activation of the reporter gene (Figs 5, S4, and S5), indicating impaired MgrB activity. We confirmed that each of these 11 MgrB variants was localized to the membrane with expression levels that were comparable to that of wild-type MgrB (S1 and S2 Fig). Below we discuss these 11 MgrB variants in more detail.

#### a. Two lysines in the MgrB cytosolic region regulate PhoQ activity

In the short N-terminal cytosolic region of MgrB (M1 to R5), we made alanine substitutions at each position except the first methionine. Here, we used the PhoQ/PhoP regulated GFP reporter (pUA66 P*_mgtLA_-gfp*) to monitor the activities of MgrB variants. Control cells having no MgrB expression showed a high intensity of GFP fluorescence, while cells expressing wild-type MgrB had much lower reporter fluorescence (Fig 3A). The K2A and K3A substitutions displayed the most effect on *gfp* reporter expression, with fluorescence comparable to the no-MgrB control (Fig 3A). F4A and R5A had about 50% and 80% activity, respectively. K2A and K3A, however, only retained about 20% activity (Fig 3A, see Materials and Methods for the definition of normalized activity).

**Fig 3.**
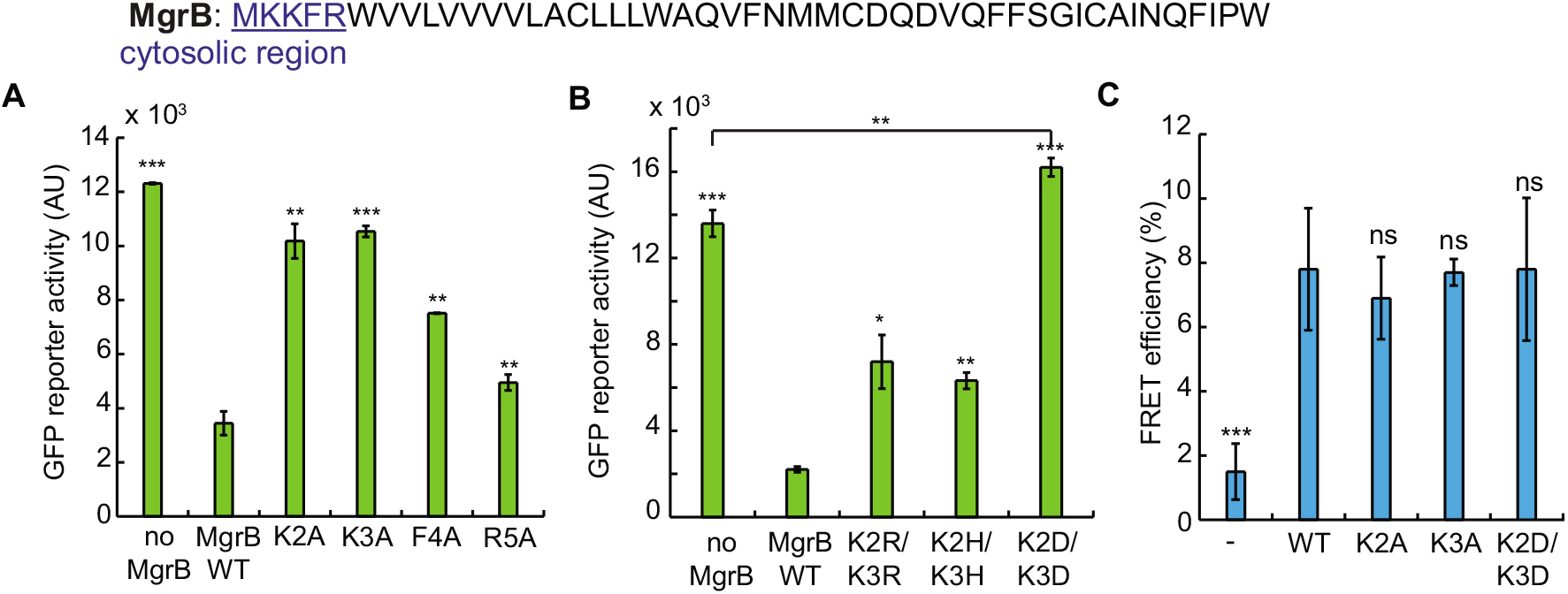
Two lysines in MgrB cytosolic region regulate PhoQ activity through their charged side chain. **A**, the activities of MgrB mutants with single alanine substitution in the cytosolic region were tested using the PhoQ/PhoP-regulated transcriptional reporter P*_mgtLA_-gfp* in a Δ*mgrB* strain. MgrB mutants were constructed in a pBAD33 vector and their expression was induced with 0.008% arabinose. **B**, the activities of MgrB K2K3 double mutants were tested as in **A**. The reporter activity change was significant (*P* = 0.0045) when comparing MgrB K2DK3D to the no_MgrB control. **C**, FRET efficiencies were measured between PhoQ-mNeonGreen and mCherry (-), wild-type MgrB (WT) or MgrB mutants (K2A, K3A, K2DK3D and K2EK3E). MG1655 Δ*phoQ* Δ*mgrB* strain was transformed with pBAD33 *phoQ-mneongreen* and pTrc99a *mcherry-mgrB* variants. Protein expression was induced with 0.008% arabinose and 10 μM IPTG. FRET was measured for each protein pair and the efficiencies were calculated as described in Materials and Methods. All data points represent averages of at least three independent experiments and error bars show SDs. The statistical significances of changes in reporter activity or FRET efficiency were calculated with *t* tests by comparing to wild type or as specified and indicated with asterisks (*** *P* ≤ 0.001, ** *P* ≤ 0.01, * *P* ≤ 0.05, ns = not significant).

To further investigate whether this loss of activity might be caused by deficient protein interactions, we used *in vivo* acceptor photobleaching FRET [38] to test complex formation between these two MgrB variants and PhoQ. In this assay, MgrB variants and PhoQ were tagged with a red fluorescent protein (mCherry) and a green fluorescent protein (mNeonGreen) respectively and co-expressed in an *E. coli* Δ*phoQ* Δ*mgrB* strain. Complex formation between tagged proteins typically results in close proximity and consequently energy transfer between the donor mNeonGreen and the acceptor mCherry, leading to quenching of the mNeonGreen fluorescence. The latter can be detected as an increase in the mNeonGreen fluorescence upon photobleaching of mCherry that abolishes FRET (see Materials and Methods). The apparent FRET efficiency could be defined as the resulting increase in the donor fluorescence divided by its fluorescence after bleaching [39]. While apparent FRET efficiencies may vary depending on the optical set-up, for intra-molecular mNeonGreen-mCherry tandem constructs the reported values are in the range of 15-20% [40]. We observed about 7.0% FRET efficiency for PhoQ-mNeonGreen/mCherry-MgrB WT, whereas the negative control, PhoQ-mNeonGreen/mCherry, showed only about 1.5% FRET efficiency. Interestingly, K2A and K3A mutants showed similar FRET efficiency as MgrB WT (Fig 3C), suggesting that substitutions at these two positions do not significantly affect the MgrB/PhoQ complex formation. Therefore, K2 and K3 may be instead directly involved in regulating PhoQ conformation and promoting a kinase-inactive state of the enzyme.

We speculated that MgrB might regulate PhoQ conformation through the positively charged side chains of K2 and K3. To test this hypothesis, we mutated K2 or K3 to a negatively charged amino acid. Mutating K2 or K3 individually to aspartate only slightly reduced MgrB function (S4 Fig A and B). Furthermore, when we substituted these two lysines individually with 19 other amino acids, many of the mutants retained more than 50% activity (S4 Fig A and B). Thus, neither K2 nor K3 alone is essential for MgrB function. We then tested the effect of replacing both K2 and K3 with aspartate. MgrB K2DK3D localized to the cell membrane similarly to wild-type MgrB (S1 Fig) but with some reduced expression (S2 Fig). At this expression level, the K2DK3D mutant appeared to be a mild PhoQ activator rather than an inhibitor and showed a 20% higher reporter activity compared to the no MgrB control (Fig 3B). Increasing MgrB K2DK3D expression, either by doubling the amount of arabinose inducer or by using a stronger promoter (*trc* promter), produced a small level of inhibition that was much less than the level of inhibition of wild-type MgrB (S4 Fig C and D). Double mutants, in which the lysines were replaced with either of the positively charged amino acids histidine or arginine (K2HK3H and K2RK3R), on the other hand, retained about 60% inhibitory activity (Fig 3B). FRET analyses also showed that MgrB K2DK3D had a FRET efficiency comparable to that of wild type (Fig 3C), indicating that this MgrB variant forms complexes with PhoQ. Taken together, our results suggest that the two lysines in the MgrB cytosolic region regulate PhoQ function through their positively charged side chains without contributing to PhoQ/MgrB complex formation in *E. coli*. We interpret the fact that single amino acid substitutions of K2 or K3 to uncharged residues has little effect on activity as indicating that the remaining lysine is sufficient to mediate the required interaction. We also note that the K3R substitution was inactive (S4 Fig B), which suggests that it is not only the charge of the lysine sidechains that are important for MgrB activity.

#### b. The TM helix is involved in PhoQ/MgrB complex formation through W20

Given that the MgrB TM sequence is necessary for its function, we wanted to further identify specific residues that might contribute to PhoQ inhibition. Polar amino acids such as N and Q can facilitate TM domain interactions by forming stable hydrogen bonds [41]. Aromatic residues are known to preferentially occur in the lipid-water interface regions [42, 43] and could be important for membrane anchoring as well as surface interactions. An MgrB sequence alignment identified four polar or aromatic residues in the transmembrane region that are highly conserved: W6, C16, W20 and Q22. Based on these considerations, we mutated the four highly conserved residues and F24 individually to alanine. The W6A and C16A substitutions did not affect MgrB function (S5 Fig and Fig 4A); the result for C16A is consistent with previous observations [20]. The F24A substitutions displayed modest effect on reporter gene expression (~8-fold higher *β*-gal activity compared to WT MgrB) and the change in activity for the Q22A mutant was not statistically significant (Fig 4A). The W20A substitution, on the other hand, greatly affected MgrB function, with *β*-gal reporter activity that was ~30-fold higher than that of WT MgrB (Fig 4A). These results were also reproduced with the GFP reporter system and untagged MgrB (S5 Fig, except that the Q22A showed significant activity loss (~70%) in this case. Collectively, the results from both approaches indicate that W20, F24 and possibly Q22 in the TM region are functionally important.

**Fig 4.**
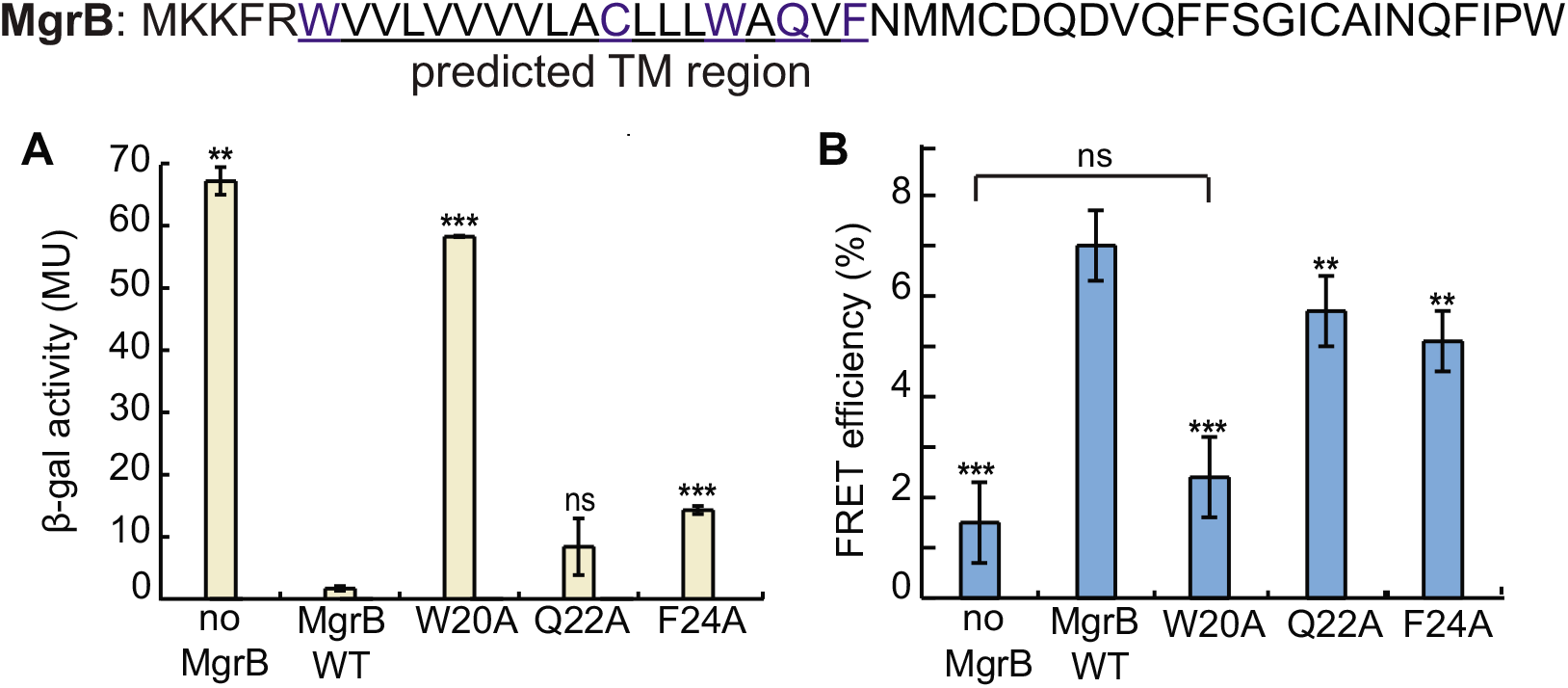
W20 in MgrB transmembrane domain is essential for PhoQ/MgrB complex formation. **A**, *β*-galactosidase levels were measured in a Δ*mgrB* strain containing the PhoQ/PhoP-regulated transcriptional reporter P*_mgtA_-lacZ* (AML67) and also containing either an empty vector (pAL39), wild-type MgrB (pAL38), or the indicated mutants MgrB W20A (pSY72), Q22A (pSY73) and F24A (pSY74). Inducer (IPTG) was not added as the leaky expression from the *trc* promoter resulted in sufficient levels of protein. Mutants with *β*-galactosidase activity ≥ 5-fold above wild type were selected for further analysis. Data represent averages and ranges for two independent experiments. MU = Miller Units. **B**, FRET efficiencies were measured between PhoQ-mNeonGreen and indicated mCherry-MgrB variants as described in Figure 3C. When compared to wild type, the reduction of FRET efficiencies is significant according to *t* test (*** *P* < 0.001, ** *P* < 0.01). Data points represent averages of at least three independent experiments and error bars show SDs. The statistical significances of changes in reporter activity or FRET efficiency were calculated with *t* tests by comparing to wild type and indicated with asterisks (*** *P* ≤ 0.001, ** *P* ≤ 0.01, * *P* ≤ 0.05, ns = not significant).

Our bacterial two-hybrid data suggest that the MgrB TM region plays a role in establishing physical contact with PhoQ. To pinpoint the essential residues in MgrB TM region that are responsible for MgrB/PhoQ complex formation, we performed FRET analyses on MgrB TM point mutants (Fig 4B). Compared to wild-type MgrB, which had 7.0% FRET efficiency, Q22A and F24A showed slightly reduced FRET efficiency, 5.7% and 5.2%, respectively. W20A, on the other hand, produced a much greater change, with FRET efficiency reduced to 2.4%, which was comparable to the negative control. These results indicate that W20 in the TM region is a key residue for MgrB/PhoQ complex formation.

#### c. The periplasmic region has six functionally important residues

In the periplasmic region, mutants C28A and C39A showed the highest effects (80-90 fold higher *β*-gal activity than WT MgrB), consistent with previous data indicating that these cysteines are important for MgrB activity [20]. In addition, mutants G37A, D31A, F34A and W47A displayed a significant increase in reporter gene expression – approximately 30-, 8-, 10-, and 10-fold relative to WT MgrB, respectively (Fig 5). These results were also reproduced with GFP reporter system and untagged MgrB constructs (S5 Fig). The fact that mutating residues C28, G37 and C39 showed the strongest effect on the reporter activity suggests that these residues are likely important for structural stability of the MgrB periplasmic region.

**Fig 5.**
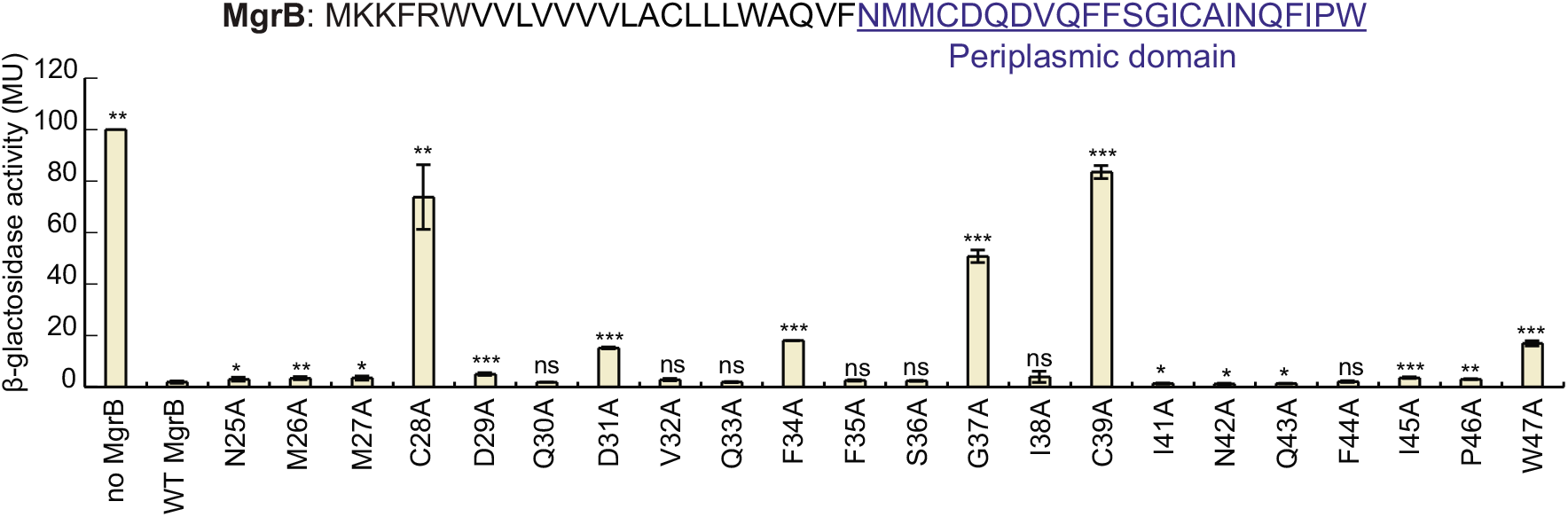
Effects of site-directed alanine substitutions of MgrB periplasmic residues on PhoQ activity. *β*-galactosidase levels were measured in a Δ*mgrB* strain containing the PhoQ/PhoP-regulated transcriptional reporter P*_mgtA_-lacZ* (AML67) and also containing either an empty vector (pAL39), wild-type MgrB (pAL38), or the indicated mutants (pJC1 through pJC7 and pTG1 through pTG15; see Table S2 for details). Inducer (IPTG) was not added as the leaky expression from the *trc* promoter resulted in sufficient levels of protein. Data for wild-type MgrB and each mutant was normalized to the no-MgrB control. Mutants with *β*-galactosidase activity ≥ 5-fold above wild type were selected for further analysis. Averages and standard deviations for three independent experiments are shown. MU = Miller Units. The statistical significances of changes in reporter activity were calculated with *t* tests by comparing to wild type and indicated with asterisks (*** *P* ≤ 0.001, ** *P* ≤ 0.01, * *P* ≤ 0.05, ns = not significant).

### A charged residue in the PhoQ TM2 – HAMP junction is involved in MgrB inhibition

Our results suggest that the two N-terminal lysines in MgrB regulate PhoQ activity through their charged side chains. To further explore which PhoQ residue(s) in PhoQ interact with these lysines, we screened for PhoQ mutants for which MgrB K2DK3D showed increased inhibitory activity. Based on the MgrB and PhoQ topologies, we hypothesized that K2 and K3 of MgrB interact with residues in the PhoQ TM2-HAMP region (Fig 6A). We therefore mutated this region of PhoQ by error-prone PCR and screened for PhoQ mutants that have reduced activity in the presence of MgrB K2DK3D. About 100 colonies with decreased green fluorescent reporter protein were selected and the *phoQ* genes were sequenced. Mutants with missense mutations are highlighted in the supplementary file (PhoQ_RM_screening). Most of the missense mutations were clustered in/near TM2-HAMP junction and the second helix of HAMP domain. We focused on the former, as residues in this region are more likely to be close enough to interact with the lysines in MgrB. Among them, R219 (labeled in Fig 6A), which was the most frequent among identified mutations, appears to be the most promising residue.

**Figure 6.**
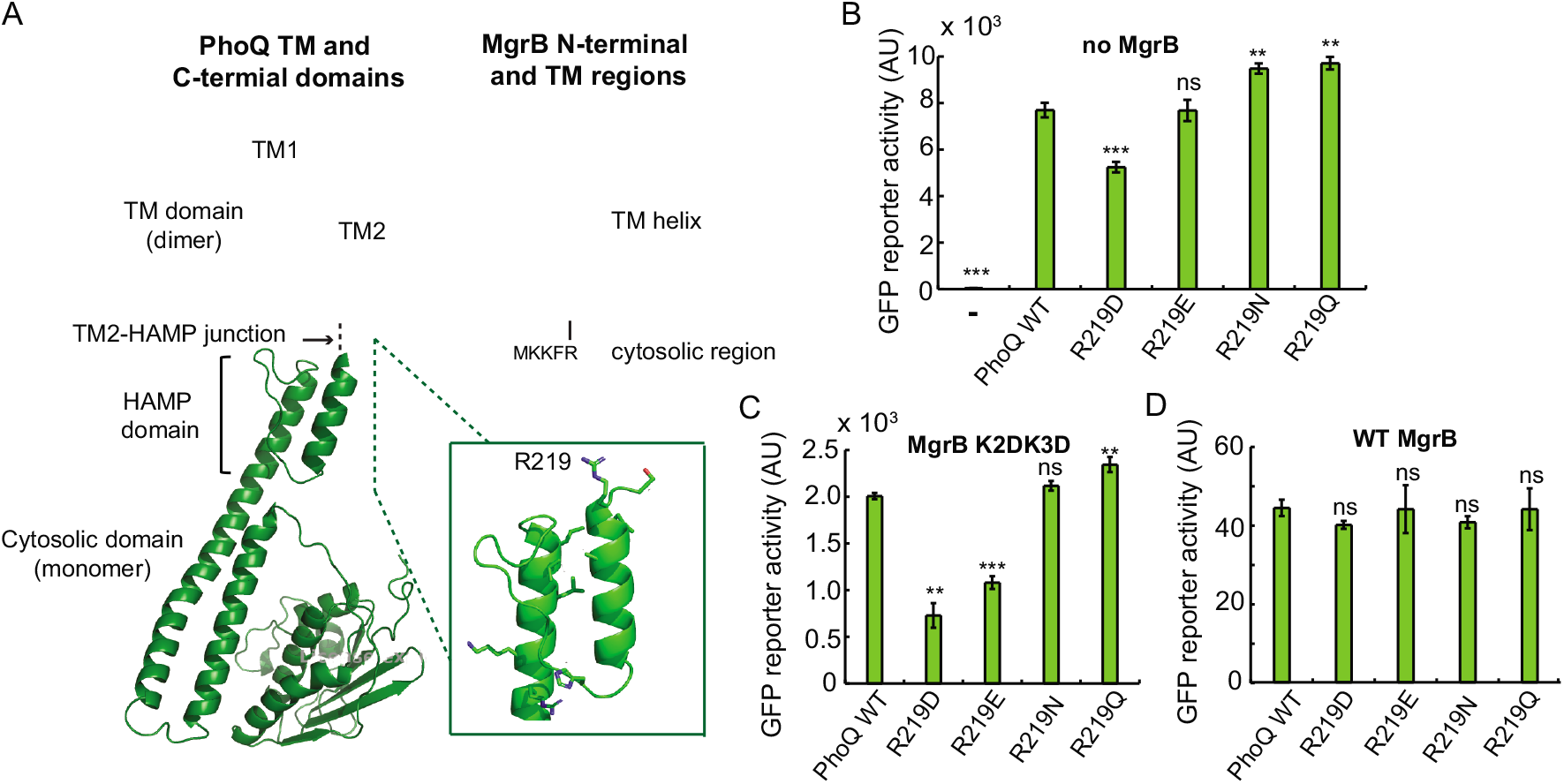
PhoQ R219 is involved in interactions with MgrB K2 and K3. Models of TM [14] and cytosolic domains [44] of PhoQ and MgrB are shown in (**A**). The MgrB TM model was constructed *ab initio* via Rosetta (https://www.rosettacommons.org). PhoQ R219 residue is labeled and shown with side chain in a zoomed-in view. **B**, the activity of PhoQ R219 mutants was tested using the PhoQ/PhoP-regulated transcriptional reporter P*_mgtLA_-gfp* in a Δ*phoQ*Δ*mgrB* strain. PhoQ mutants were expressed from the plasmid pBAD33 with 0.008% arabinose. The activities of PhoQ R219 mutants were tested in the presence of MgrB K2DK3D (**C**) or wild type (**D**) using the same GFP reporter as in **B**. MgrB variants were expressed from a pTrc99a vector with no inducer (IPTG) added, as the leaky expression from the *trc* promoter resulted in sufficient levels of protein. Averages and standard deviations for three independent experiments are shown in **B**-**D**. The statistical significances of changes in reporter activity were calculated with *t* tests by comparing to wild type and indicated with asterisks (*** *P* ≤ 0.001, ** *P* ≤ 0.01, * *P* ≤ 0.05, ns = not significant).

We therefore mutated PhoQ R219 to negatively charged amino acids (D and E) as well as the amino acids with amide side chains (N and Q). Compared to wild-type PhoQ, R219D has reduced activity (about 70% of WT), while the other three mutants (R219E, R219N and R219Q) have activities that are either similar to or slightly greater than that of wild type (Fig 6B). Next, we tested the activity of these PhoQ mutants in the presence of MgrB K2DK3D. Compared to wild-type PhoQ, R219D and R219E are indeed repressed further by MgrB K2DK3D (Fig 6C), suggesting that reversing the side chain charge of PhoQ R219 compensates the effect of the charge inversion of MgrB N-terminal lysines. The inhibition of PhoQ R219N and R219Q by MgrB K2DK3D, on the other hand, was comparable to the inhibition of wild-type PhoQ, indicating that charge but not hydrogen bonding is important for interactions of R219D and R219E with MgrB K2DK3D. We also found that wild-type MgrB inhibited all four PhoQ R219 mutants to a similar extent as the inhibition of wild-type PhoQ (Fig 6D), suggesting the existence of other factors involved in PhoQ inhibition by wild-type MgrB.

### Inhibition of *E. coli* PhoQ by MgrB orthologs

MgrB orthologs have previously been identified in several enterobacterial species [18]. To examine the activities of these MgrB natural variants on *E. coli* PhoQ, we expressed MgrB from *Klebsiella pneumoniae, Yesinia pestis, Photorhabdus laumondii, Proteus penneri, Providencia stuartii, Serratia sp.MYb239, Enterobacter ludwigii* and *Salmonella enterica serovar typhimurium* in *E. coli* K-12 MG1655. All orthologs except one (MgrB-*E. ludwigii*) localized to the membrane (S1 Fig F). MgrB-*E. ludwigii*, which is missing the positively charged residues (K2, K3) in its N-terminus showed defective localization, suggesting the lack of a signal peptide. We found that most of the orthologs expressed at levels comparable to that of *E. coli* MgrB, but the ortholog from *E. ludwigii* expressed quite poorly in *E. coli* (~5-fold lower than *E. coli* MgrB see Fig 7). We then measured reporter gene expression for PhoP-regulated transcription as a function of activity of each MgrB ortholog (Fig 7). Unsurprisingly, *E. ludwigii* showed high reporter gene expression, indicating low PhoQ inhibition. MgrB from *K. pneumoniae, S. sp.MYb239*, and *S. typhimurium* showed small but significant increases in reporter activity (~4-fold, 2.5-fold, and 2-fold, respectively). Interestingly, *Y. pestis, P. penneri* and *P.stuartii* MgrB produced reporter activities that were significantly lower than that of *E. coli* MgrB, suggesting that they repress *E. coli* PhoQ more strongly than does *E. coli* MgrB.

**Figure 7.**
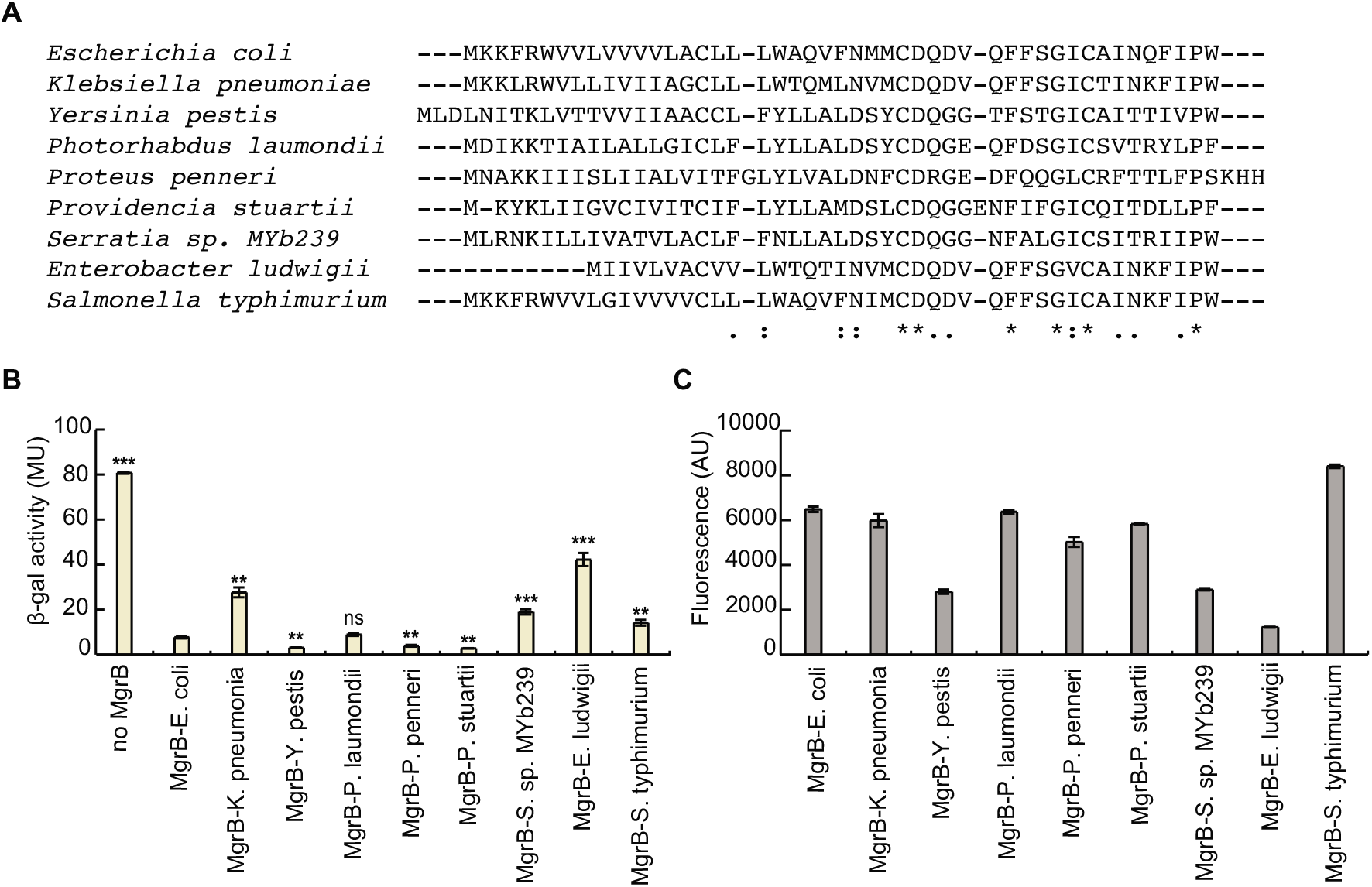
Effect of MgrB orthologs on *E. coli* PhoQ activity. **A**, Sequence alignment of MgrB orthologs. **B**, *β*-galactosidase levels of a Δ*mgrB* strain containing the PhoQ/PhoP-regulated transcriptional reporter P*_mgtA_-lacZ* (AML67) and also containing either an empty vector (pAL39), wild-type MgrB (pAL38), or a plasmid expressing the indicated MgrB orthologs. Inducer (IPTG) was not added as the leaky expression from the *trc* promoter resulted in sufficient levels of protein. Averages and standard deviations for three independent experiments are shown. MU = Miller Units. The statistical significances of changes in reporter activity were calculated with *t* tests by comparing to wild type and indicated with asterisks (*** *P* ≤ 0.001, ** *P* ≤ 0.01, * *P* ≤ 0.05, ns = not significant). **C**, GFP-MgrB expression levels for cells expressing MgrB orthologs were imaged by fluorescence microscopy, and GFP fluorescence levels were quantified (see Methods for details). Data represent averages and ranges for two independent experiments comprised of 200 cells each, AU = Arbitrary Units.

### MgrB interaction with non-cognate histidine kinases

To test if MgrB is capable of physically interacting with histidine kinases other than PhoQ, we made constructs of *E. coli* EnvZ, AtoS, CpxA, and PhoR fused to the T25 fragment of adenylyl cyclase and performed BACTH analysis by spot assays as well as by measuring *β*-gal activities of liquid cultures. Intriguingly, cells expressing T18-MgrB and either T25-AtoS, - EnvZ, or -PhoR showed a high level of activity, although less than the *β*-gal activity of cells expressing T18-MgrB and T25-PhoQ (Fig 8). Cells expressing T18-MgrB and T25-CpxA, on the other hand, displayed a level of *β*-gal activity only marginally higher than that of the negative controls. These results suggest that MgrB is capable of physically interacting with EnvZ, AtoS, and PhoR, at least in the context of the bacterial two-hybrid assay. To test whether MgrB affects the activity of these histidine kinases, we measured gene expression from promoters regulated by these histidine kinases, in the presence or absence of plasmid driven MgrB expression. However, we did not see any effect of MgrB on gene expression controlled by any of these histidine kinases (data not shown). We also tested several other histidine kinases: ArcB, BaeS, BasS, CpxA, CusS, EvgS, NtrB, RstB and TorS (S6 Fig). None of them showed interactions with MgrB that were comparable to that of PhoQ.

**Figure 8.**
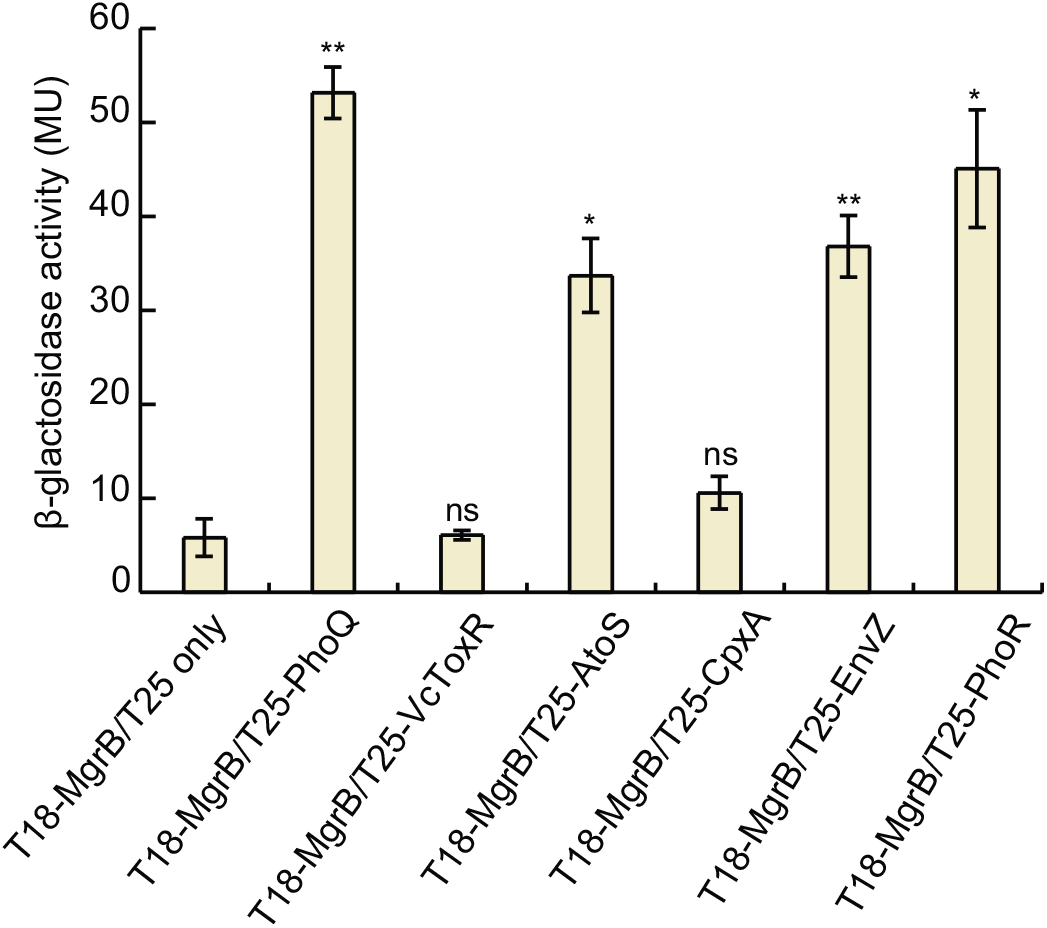
MgrB interacts with other histidine kinases in a bacterial two-hybrid assay. Bacterial two-hybrid (BACTH) assays with cells expressing protein fusions to the T18 and T25 subunits of *B. pertussis* adenylyl cyclase. *β*-gal expression from the *lac* promoter was monitored as a measure of interactions between T18-MgrB and T25 fusion with either PhoQ, AtoS, CpxA, EnvZ or PhoR. The T25 fragment alone and T25-*Vc*ToxR (*Vibrio cholerae* ToxR) serve as negative controls. *β*-gal assays were carried out in a *cyaA^-^* BACTH host strain containing *lacl*^q^ (SAM85). Data represent averages and ranges for two independent experiments. MU = Miller Units. The statistical significances of changes in reporter activity were calculated with *t* tests by comparing to the negative control (T18-MgrB/T25 only) and indicated with asterisks (*** *P* ≤ 0.001, ** *P* ≤ 0.01, * *P* ≤ 0.05, ns = not significant).

## Discussion

In this study, we have explored the regions of the small protein MgrB that are important for its activity as an inhibitor of the PhoQ sensor kinase. Using a combination of biochemical and biophysical approaches, we identified 11 amino acids in MgrB that, at least in the context of alanine substitution, are important for the protein’s inhibition of PhoQ. These functionally important residues are spread across different regions of the molecule, consistent with our observation that TM replacement or periplasmic region truncation leads to a loss of MgrB function. The fact that the MgrB TM sequence is required for the protein’s activity distinguishes MgrB from at least two other bitopic membrane proteins that regulate histidine kinases (SafA and MzrA); the TM domains of SafA and MzrA apparently serve primarily as membrane anchors [31, 32]. In addition, our results indicate that the MgrB TM region together with the small cytoplasmic region interacts with PhoQ, and that residue W20 in the TM region is essential for MgrB/PhoQ complex formation. Tryptophan is often found in transmembrane proteins, especially at the ends of TM helixes as a membrane anchor due to its unique amphipathic side chain [45, 46]. W20 in MgrB does not appear to position near the computationally predicted membrane/periplasm interface. It is possible that the unique side chain of W20 interacts with polar and/or aromatic residues in the PhoQ transmembrane domain, such as the functionally important N202 [47]. The mechanism of how W20 mediates PhoQ/MgrB complex formation awaits further investigations.

We identified two lysines in the MgrB cytosolic region that are important for the protein’s function. Although these residues are not individually essential, replacing both with negatively charged residues significantly affected MgrB activity. A screen for PhoQ mutants that increase the inhibitory activity of MgrB K2DK3D identified R219 in PhoQ. When this residue was mutated to a negatively charged amino acid, the inhibitory activity of MgrB K2DK3D was partially restored, suggesting the possibility that MgrB inhibitory activity of PhoQ is in part mediated by charge-charge repulsion between K2K3 in MgrB and R219 in the PhoQ TM2-HAMP junction. This hypothesis is consistent with the observation that K2 and K3 did not contribute to MgrB/PhoQ complex formation, since the HAMP domain is conformationally dynamic and transduces signals via conformational changes [44, 48]. It is also noteworthy that the TM2-HAMP junction has been reported to be essential for signaling by other membrane proteins such as the chemoreceptor Tar [49]. In fact, some modifications of the TM2-HAMP junction result in signal inversion in *E. coli* [49]. The partial suppression by PhoQ R219D indicates the existence of other factors involved in PhoQ/MgrB interaction at this residue. Mutating arginine to aspartate not only changes the charge but also the length of the side chain (as well as other properties), which may have an impact on PhoQ/MgrB interaction. Interestingly, we observed activity inversion (from activation to inhibition) of MgrB K2DK3D when increasing its expression. Considering its reduced membrane localization and possible heterogeneous cell population under arabinose induction, it is possible that the mild activating activity we observed for lower expression level was due to factors other than the direct effects of the mutant MgrB on PhoQ. Alternatively, different amounts of MgrB K2DK3D might lead to different energy states or conformations of the PhoQ TM2-HAMP junction, and thus changes in PhoQ activity. Lastly, we note that some MgrB orthologs have a longer N-terminal cytosolic region and some do not have positive charged residues at position 2 and 3. Due to the dynamic nature of the HAMP domain and the TM2-HAMP junction, it is possible that these MgrB orthologs have different interactions with PhoQ.

Previous work indicates that the periplasmic region of MgrB and, in particular, two conserved cysteines in this region, are critical for PhoQ inhibition [18, 20]. To identify other periplasmic residues in MgrB that are required for PhoQ inhibition, we analyzed individual alanine substitutions across the entire periplasmic region. Remarkably, despite the small size of this region and the level of conservation, substitutions at most positions had little effect on MgrB activity. Of the 22 residues in the periplasmic region, only six were individually important for full activity: the two conserved cysteines (C28 and C39), as previously noted [20], as well as G37, D31, F34 and W47. Furthermore, substitutions at these latter four residues lowered but did not eliminate MgrB activity. Interestingly, mutations for each of these positions are also associated with colistin-resistant *Klebsiella* isolates [23, 24, 50–55]. Taken together, our findings show that the sequence determinants for MgrB activity are not restricted to either one of the regions – cytosolic, periplasmic or TM, and that functionality is mediated by collective interactions of key amino acid residues scattered across the polypeptide.

We also tested a number of MgrB orthologs derived from other enterobacteria with varying sequence similarity to *E. coli* MgrB. Most orthologs that we tested efficiently inhibited *E. coli* PhoQ. The slight decrease in activities of *K. pneumoniae* and *S. typhimurium* MgrB could be due to the presence of a glycine residue within the TM regions, which may affect the conformational flexibility of the TM helix and cause a defect in binding and inhibiting PhoQ. On the other hand, *Y. pestis, P. penneri* and *P.stuartii* MgrB appear to be more efficient than *E. coli* MgrB in inhibiting *E. coli* PhoQ. It is notable that the tryptophan residue (W20) found in the TM region of *E. coli* MgrB is replaced by a tyrosine (Y20) in these orthologs. It is possible that a hydrogen bond involving Y20 or the smaller size of tyrosine relative to tryptophan affects the interaction of MgrB with PhoQ.

Unexpectedly, we also found that MgrB lacking its periplasmic region, as well as full-length MgrB, interact with several other histidine kinases in addition to PhoQ in a bacterial two-hybrid assay. However, we did not observe a significant effect of MgrB on gene expression regulated by these histidine kinases. The TM domains of many dimeric histidine kinases such as PhoQ and EnvZ are believed to form a four-helix bundle [48]. Our results may reflect an interaction between the MgrB TM and a conserved conformation of the transmembrane domain shared by multiple histidine kinases. The cytosolic and periplasmic regions of MgrB apparently provide additional interactions required for specific regulation of PhoQ. It will be interesting to determine if there are other small membrane proteins that have evolved to interact with other histidine kinases using structural features similar to those of the MgrB TM region. In addition, the results from our study of MgrB, as well as studies of other small membrane proteins, may provide a platform for reengineering these proteins to regulate new targets through rational design and directed evolution.

## Materials and Methods

### Strains, plasmids and cloning

See Supporting information for tables of strains (S1 Table), plasmids (S2 Table), and primers (S3 Table) used in this study. All strains were derived from *E. coli* K-12 MG1655. Gene deletions and reporter constructs were transferred between strains using transduction with P1_vir_. Deletions in the Keio collection [56] that were used for strain construction were confirmed by PCR using primers flanking the targeted gene. Kanamycin resistance markers were excised, when required, using the FLP-recombinase-expressing plasmid pCP20 [57]. Strain SAM72 was constructed by first excising the kanamycin resistance cassette from the *ompC-mCherry* transcriptional reporter strain AFS256 to give SAM71. Kanamycin resistance from the *mzrA* deletion strain JW3067 was then transduced into SAM71 by P1 transduction. A deletion of *safA* (SAM73) was constructed as described in [28, 57]. SAM73 was then used to construct SAM76 by transducing the *mgtA-lacZ* transcriptional reporter from TIM199. Strain SAM85, which is a *lacl*^q^ containing version of BTH101 was made by conjugation of the bacterial two-hybrid host strain BTH101 (F-) with XL1-Blue (F+). This strain was used instead of BTH101 as it gave reduced background in our MacConkey plate assays. Strain JingY90 was constructed by introducing *phoQ* deletion to JingY34 using P1 transduction followed by kanamycin resistance cassette removal. Plasmids expressing *gfpA206K* fusions to *mgrB* mutants (pSY3-9, pSY72-75, pJC1-7, pTG1-15) were created by either QuikChange site-directed mutagenesis (Stratagene) or Inverse PCR [58] using pAL38 as template. Plasmids expressing *gfpA206K* fusions of *mzrA* (pSY29)), *safA* (pSY31) and *mgrB* orthologs (pSV1-8) were constructed by Gibson assembly [59]. The gene fragments for *mzrA* and *safA* were PCR amplified from the MG1655 genome, and gene fragments corresponding to the different *mgrB* orthologs were synthesized commercially (Integrated DNA Technologies, Inc.). Plasmids pSY32 (MzrA TM swap) and pSY33 (SafA TM swap) were made by Inverse PCR using pSY29 and pSY31 as templates, respectively. Plasmid pSY34 expressing T18-MgrB-24 only was created using pAL33 as template by Inverse PCR. For all the plasmids mentioned above, see S3 Table for specific primer sequences used for each construction. Plasmids pSY61, pSY64 and pSY68 were constructed as follows: *atoS*, *cpxA*, and *phoR* were PCR-amplified using MG1655 genome as the template and the following primer pairs – atoS-xbaI-U1/atoS-kpnI-L1, cpxA-xbaI-U1/cpxA-kpnI-L1 and phoR-xbaI-U2/phoR-kpnI-L1, respectively, digested with XbaI/KpnI restriction enzymes and then cloned into pKT25 at XbaI/KpnI sites. Plasmid pJY151 was constructed by inserting PCR amplified *mgrB* to SacI/XbaI sites in pBAD33RBS vector. Plasmid pJY569 expressing *mcherry-mgrB* was generated by inserting *mgrB* downstream of *mcherry* at BamHI/PstI sites in pBO2 [60]. Plasmid pJY573 was generated by inserting *mNeonGreen* downstream of phoQ at SalI/HindIII sites in pBAD33RBS phoQ [14]. Plasmids expressing *mgrB* variants were constructed using Q5 site-directed mutagenesis kit (NEB). All constructs were confirmed by DNA sequencing.

### Media and growth conditions

Liquid cultures were grown at 37 °C with aeration, unless otherwise indicated, in either LB Miller medium (Fisher Scientific) or in minimal A medium [61] containing 0.2% glucose, 0.1% casamino acids and 1 mM MgSO_4_. For routine growth on solid medium, LB or minimal medium containing bacteriological grade agar (Fisher Scientific) was used. The antibiotics ampicillin, kanamycin and chloramphenicol were used at concentrations 50–100, 25-50 and 20–34 μg mL^−1^, respectively. The *lac* and *trc* promoters were induced with isopropyl *β-D-1*-thiogalactopyranoside (IPTG) at a final concentration of 10 μM in FRET analysis or 1 mM when indicated. When IPTG was not mentioned in the description of the culture conditions, the basal transcription from the *trc* promoter was used to drive expression. The araBAD promoter was induced with 0.008% arabinose. For bacterial two-hybrid screening, MacConkey/maltose indicator plates were prepared as described [36] using MacConkey agar base powder (Difco™), 100 μg mL^−1^ ampicillin, 25 μg mL^−1^ kanamycin, 0.5 mM IPTG and 1% maltose.

### *β*-galactosidase reporter gene assay

Cultures of strain AML67 carrying *gfpA206K* fusions of either wild-type *mgrB* (pAL38), *mgrB* mutants (pSY3-9, pSY72-75, pJC1-7, pTG1-15), *mgrB* orthologs (pSV1-8) or *gfpA206K-only* (pAL39) plasmids were grown overnight in LB with 50 μg mL^−1^ ampicillin. The next day, they were diluted 1: 1000 into the same medium and grown to OD_600_ ~0.3-0.4 at 37 °C. Cells were permeabilized with chloroform/SDS and assayed as in [61]. For high-throughput measurements in a 96-well format, the protocol was modified as described in Thibodeau *et al*.[62].

### GFP reporter assay

*E. coli imp-4213* strain carrying GFP reporter plasmid (pUA66 P*_mgtLA_-gfp*) was grown overnight at 37 °C in Luria–Bertani (LB) medium supplemented with 1 or 10 mM MgSO_4_, then diluted 1:100 to fresh MgSO_4_ supplemented LB medium. DMSO, MgrB or SafA C-terminal peptides (synthesized by Genscript) were added to the culture with indicated amount. The cultures were grown at 37 °C with vigorous shaking for two hours. The fluorescence of cells was then monitored with a BD LSRFortessa SORP flow cytometer (BD Biosciences) and the acquired data were analyzed as described before [14]. To test the function of MgrB variants, *E. coli* MG1655 *ΔmgrB* strain was transformed with GFP reporter plasmid (pUA66 P*_mgtLA_-gfp*) [14] and a pBAD33 vector carrying wild-type or mutant *mgrB* genes. The resulting transformants were grown overnight at 37 °C in LB medium supplemented with 10 mM MgSO_4_, then diluted 1:100 to fresh LB medium supplemented with 1 mM MgSO_4_ and 0.008% arabinose. The cultures were grown at 37 °C with vigorous shaking for two hours to reach early log phase (OD=0.4-0.5). The fluorescence of cells was monitored and the acquired data were analyzed as described before [14]. The normalized MgrB mutant activity was calculated as following:

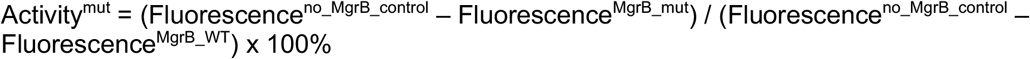

Where appropriate, 50 μg mL^−1^ kanamycin and 34 μg mL^−1^ chloramphenicol was added to the growth media.

### Acceptor photobleaching FRET analysis

The expression of PhoQ-mNeonGreen and mCherry-MgrB was induced in *E. coli* ΔphoQΔmgrB strain with 0.008% arabinose and 10 μM IPTG, respectively. The cultures were grown to mid-log phase (OD=0.6) in LB media supplemented with 10 mM MgSO_4_ and appropriate antibiotics. Cells were harvested and washed two times with pre-chilled tethering buffer (20 mM potassium phosphate pH7.0, 1 μM methionine, 10 mM lactic acid and 10 mM magnesium sulfate). The surface of a glass-bottom 24-well plate (Greiner) was treated with 0.1% poly-L-lysine (Sigma) for 10 minutes at room temperature followed by rinsing with tethering buffer. Cells were then added to the wells and incubated at room temperature for 10 minutes to allow attachment. Unattached cells were removed by washing two times and attached cells were overlaid with tethering buffer afterwards.

Acceptor photobleaching FRET was performed using a dual layer Nikon Ti-E inverted fluorescence microscope equipped with a fluorescence lamp (X-cite Exacte, Lumen Dynamics), Perfect Focus System (PFS) and NIS-Elements AR software (version 4.40 Nikon). Images were acquired through a 40x Plan Apo NA 0.95 objective in the mNeonGreen (donor) and mCherry (acceptor) channels, excitation power was adjusted both by controlling the fluorescence lamp output and with ND filters. Specifically, the donor was excited at 482/18 nm and its emission was detected at 525/50 nm and the acceptor was excited at 585/29 while its emission was detected at 647/57. Excitation and emission filters were mounted respectively on excitation and emission filter wheels, while beam splitters (495LP for mNeonGreen excitation and 605LP for mCherry excitation) were on the lower dichroic wheel of the microscope. Images were recorded using an iXon 897-X3 EM-CCD camera (Andor), the acquisition time was set to 1 second for both channels and EM gain was kept in the range of 100-280. Acceptor (mCherry) photobleaching was induced by a 150 mW DPSS 593 nm laser (ACAL Bfi) that was reflected on the sample by a zt594 DCRB laser beam splitter (AHF) mounted on the upper dichroic wheel.

For each well we acquired two to three sequences of images on isolated sample areas with the following protocol: (a) first, two images were taken in the acceptor channel; (b) then sixty images were taken in the donor channel followed by (c) ten seconds acceptor photobleaching (no image acquisition); afterwards, (d) forty images were taken in the donor channel; and then (d) two images were taken in the acceptor channel. In order to avoid donor fluorescence recovery from interrupted illumination, the sample was continuously illuminated with a 482/18 nm excitation light, even during step (c). FRET efficiency was calculated as the donor signal increase divided by the total donor signal after acceptor photobleaching. In order to correct for the donor photobleaching present during steps (b) to (d), we performed linear fitting (RStudio) of the donor fluorescence signal versus time for both pre- and post-bleaching curves.

### Bacterial two-hybrid assays

For the colorimetric spot assay, multiple clones from the transformation plate were inoculated in 3 mL of LB containing 100 μg mL^−1^ ampicillin and 25 μg mL^−1^ kanamycin. (Several clones were picked in order to reduce heterogeneity [36]. Cultures were grown overnight at 30 °C with shaking. The next day, 3 μL of each culture was spotted on MacConkey/maltose indicator plates and plates were then incubated at 30 °C. For each pair of plasmids, triplicate experiments were performed. Color change on the indicator plates was recorded after 48 h of incubation. Alternatively, *β*-gal assays were performed on the overnight liquid cultures prepared as described for the colorimetric assay.

### Random mutagenesis of the PhoQ TM2-HAMP region

The GeneMorph II random mutagenesis kit (Stratagene) was used to mutagenize the TM2-HAMP-encoding region of *phoQ* (in pBAD33 *phoQ*) using primers that specifically target this region. The library was transformed into an *E. coli ΔphoQΔmgrB* strain carrying a plasmid GFP reporter (pUA66 *PmgtL-gfp*) and an MgrB K2DK3D expression plasmid (pTrc99a *mgrB K2DK3D*). The transformants were grown on LB plates containing arabinose to induce PhoQ expression. *E. coli* colonies with repressed PhoQ have weak green fluorescence. Dim colonies were selected and their plasmids were sequenced to identify mutations in PhoQ. Site-directed mutagenesis was performed using Q5 site-directed mutagenesis kit (NEB).

## Supporting information

PhoQ RM screening

MgrB alignment

## Acknowledgements

We thank J. Zhu for the BACTH plasmid pKT25-toxR; S. Hoch and S. G. Sierra for technical assistance; members of the Goulian, Sourjik, Zhu and Binns labs for helpful discussions; P. Shah for a critical reading of the manuscript. This work was supported by NIH award GM118059 (to B.N.), NIH award GM080279 (to M.G.) and NSF award DMR11-20901 (to M.G.).

## Author contributions

S.S.Y., J.N.C. V.S., J.Y. and M.G. designed the experiments. S.S.Y., J.Y., G.M., T.G., S.V. and J.N.C. performed the experiments. B.N. contributed new reagents/analytical tools. S.S.Y., T.G., S.V., J.N.C., G.M., V.S., J.Y. and M.G. analyzed the data. S.S.Y., J.Y., V.S., M.G. wrote the manuscript and all authors edited the manuscript.

## Supporting information

### S1 Text. Materials and methods

#### Fluorescence microscopy

Strains were grown in minimal medium containing with antibiotics as needed and single-cell measurements were performed essentially as described previously [21]. For MgrB localization experiments, saturated overnight cultures of AML67 strains carrying *gfpA206K* fusions of either wild-type *mgrB* (pAL38), *mgrB* mutants (pSY3-9, pSY72-75, pJC1-7, pTG1-15), *mgrB* orthologs (pSV1-8) or *gfpA206K-only* (pAL39) were diluted 1:1,000 into fresh medium containing 100 μg mL^−1^ ampicillin and grown at 37 °C with aeration to an OD_600_ ~0.2–0.3 (~4-5 h) before visualization. For single-cell fluorescence measurements from transcriptional reporter strains, cultures were grown as outlined above, then rapidly cooled in an ice-water slurry and streptomycin was added to a final concentration of 250 μg mL^−1^ to stop protein synthesis. Microscope slides were prepared with 1% agarose pads and cell fluorescence was measured by imaging and quantified with in-house software as described previously [21, 63].

#### Membrane protein isolation and Western blot analysis

Cells expressing Flag tagged MgrB variants were harvested and resuspended in lysis buffer (50mM Tris-Cl pH8.0, 100 mM NaCl and protease inhibitors). The membrane fraction was collected as described before with modifications [34]. Specifically, cells were disrupted by homogenization at 4 °C and cell debris were remove by centrifugation at 2,300 x g for 10 minutes. The lysate was centrifuged again at 21,000 x g for 60 minutes at 4 °C to pellet the membrane fraction. The membrane pellet was washed with 1 ml lysis buffer and dissolved in SDS sample buffer overnight at 4 °C. The samples were heated at 60 °C for 10 minutes and the proteins were separated on a 16% tricine gel. The separated proteins were then transferred to Immobilon-P^SQ^ PVDF membrane (Merckmillipore) in transfer buffer containing 20% methanol and Flag-flexilinker-MgrB was detected with anti-Flag primary antibody (Sigma) and IRDye 800CW-conjugated secondary antibody (LI-COR). The protein bands were visualized with the Odyssey CLx imaging system (LI-COR) and quantified with ImageJ.

**S1 Table.**
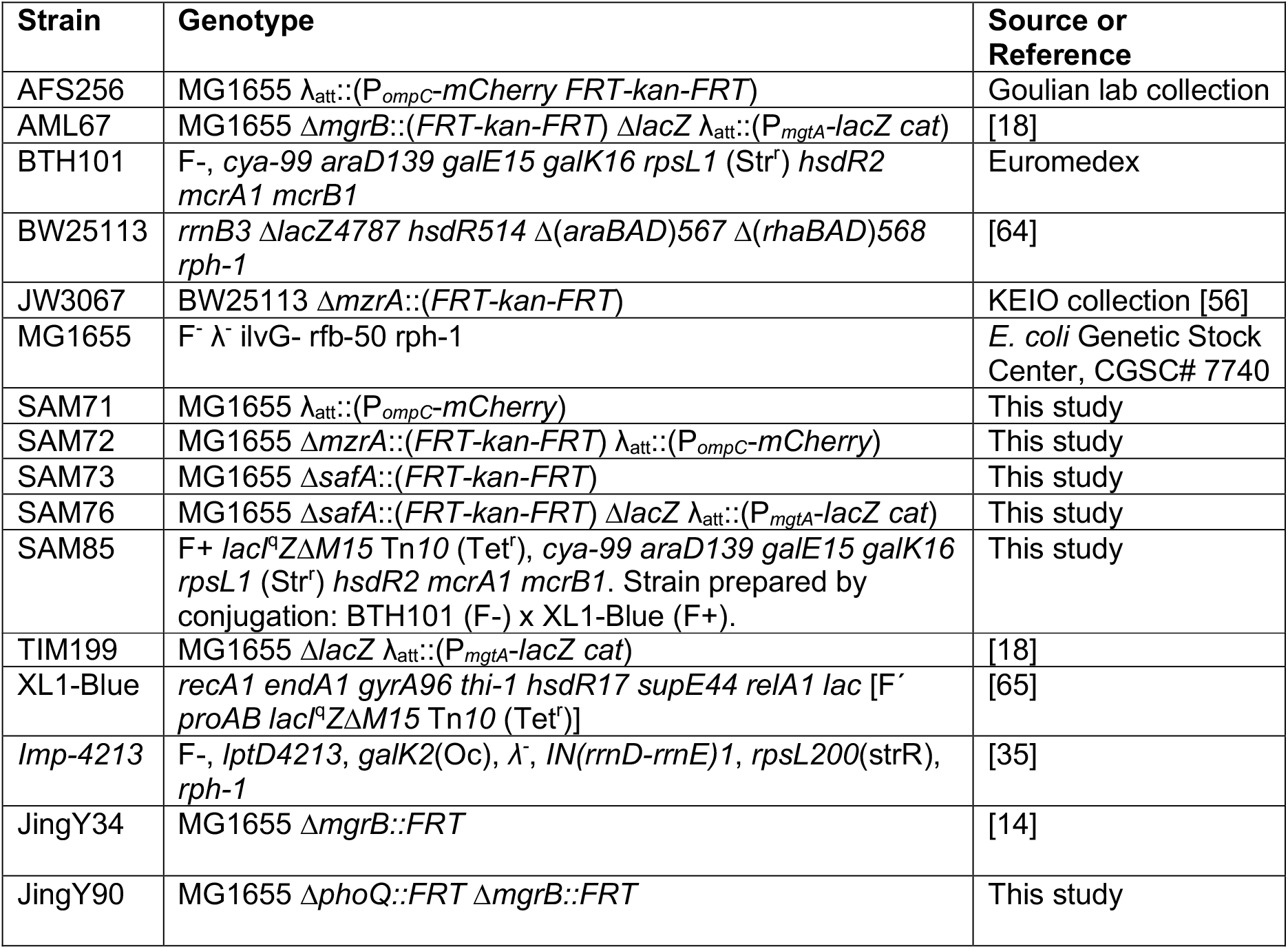
Strains

**S2 Table.**
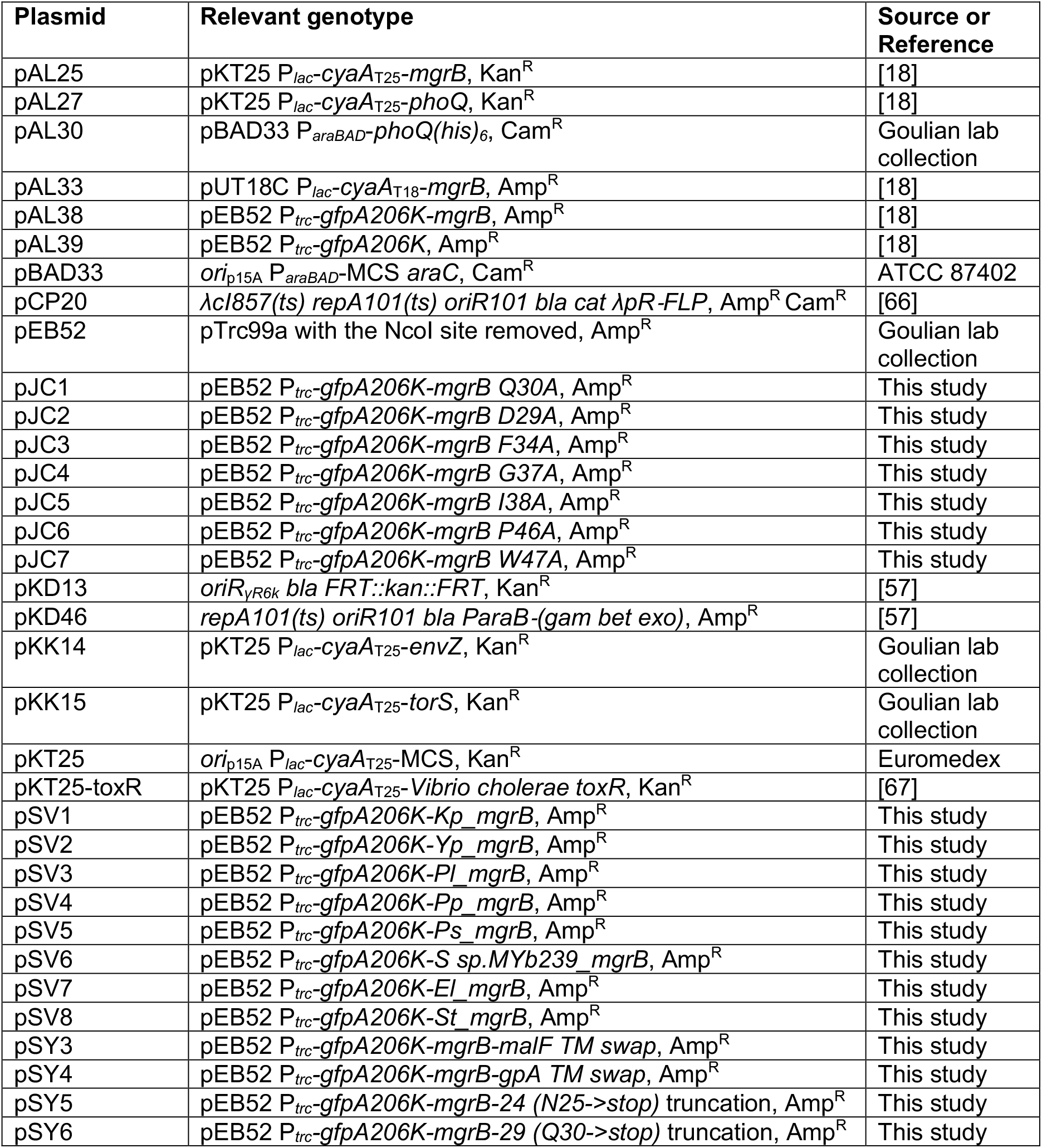
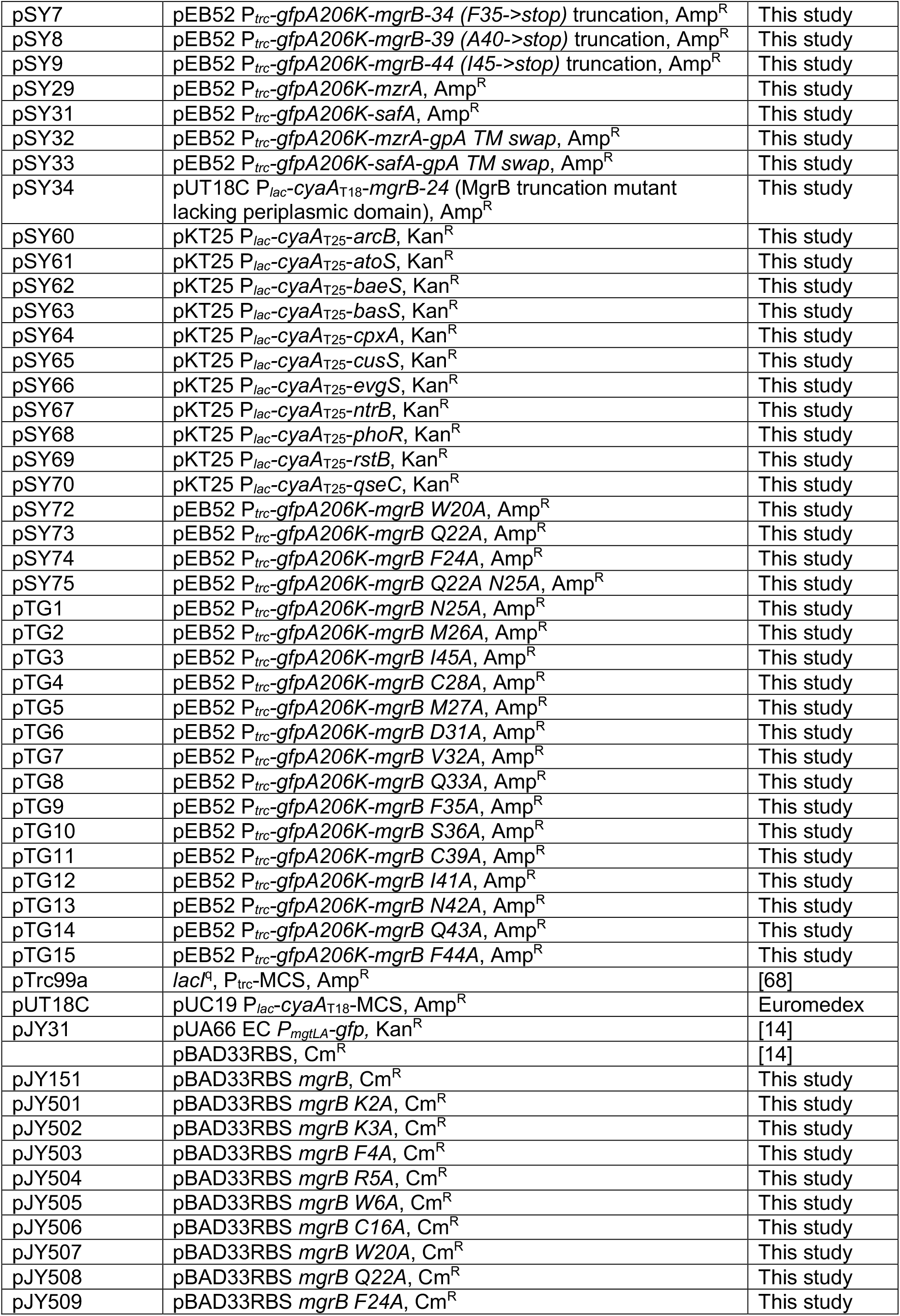
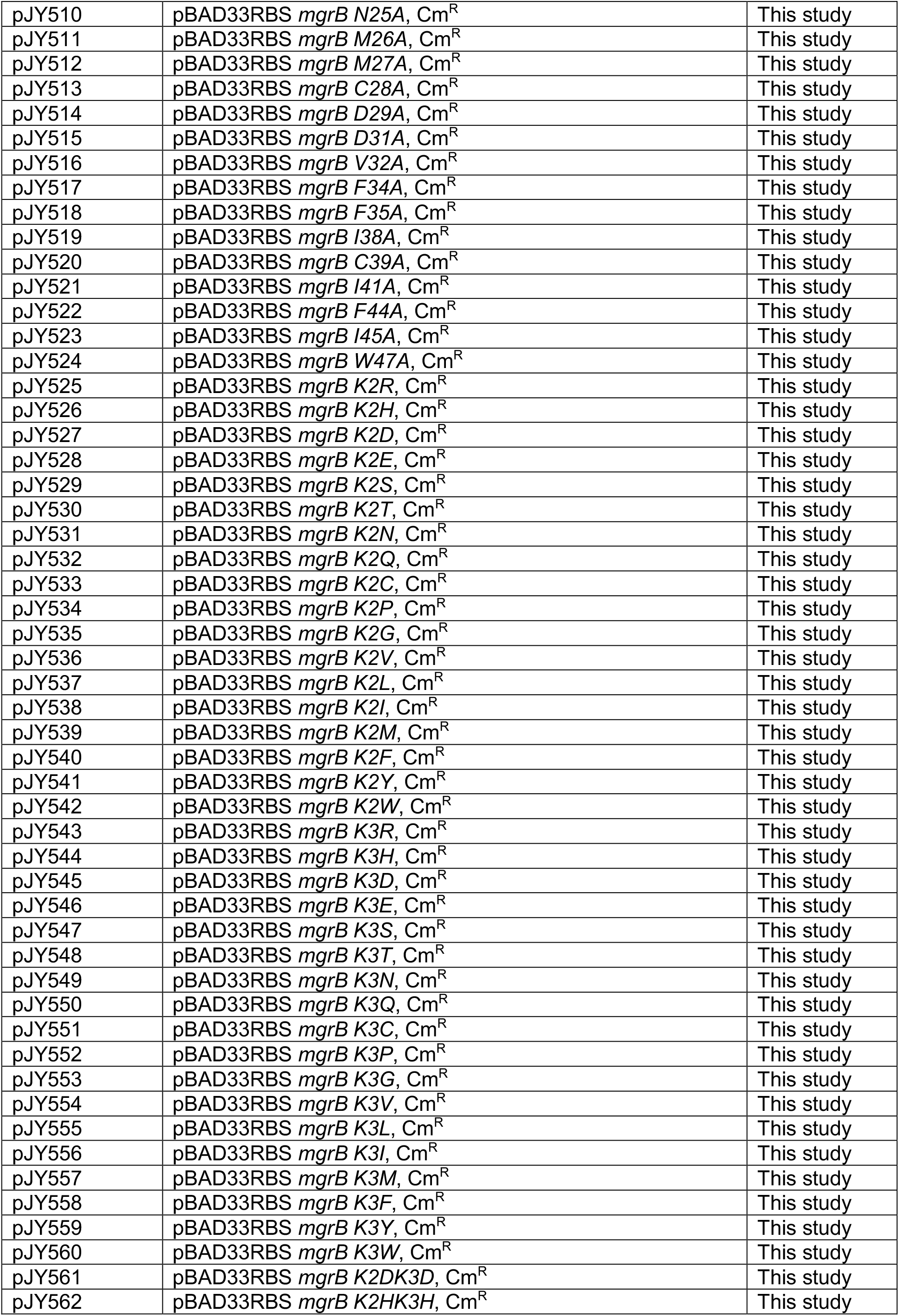
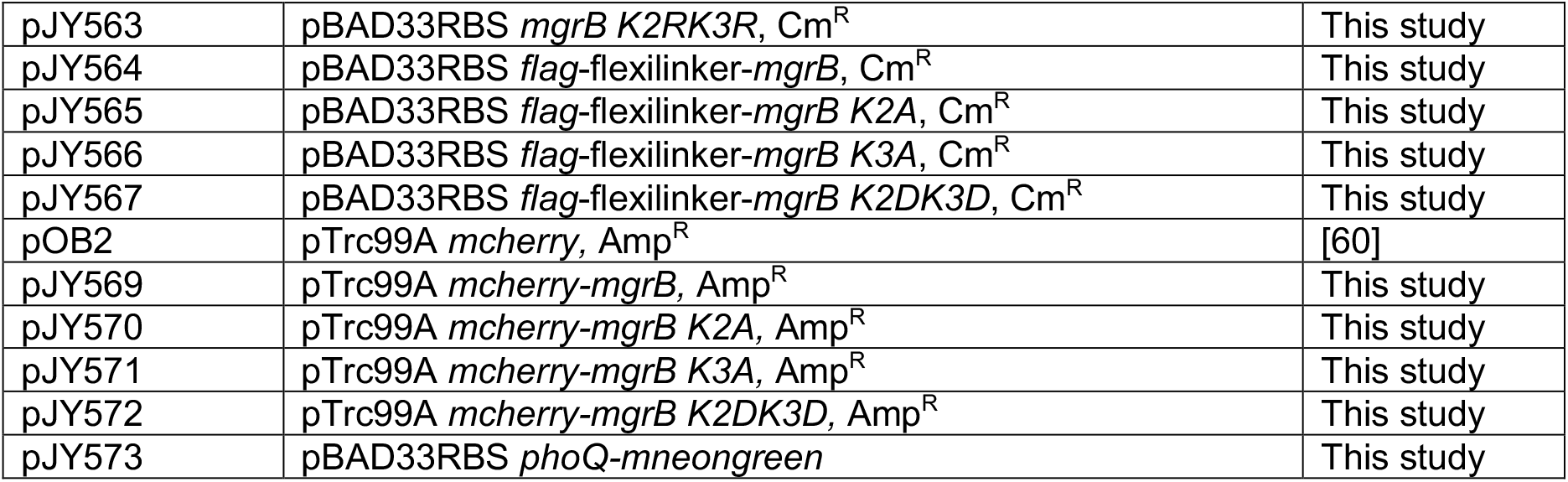
Plasmids

**S3 Table.**
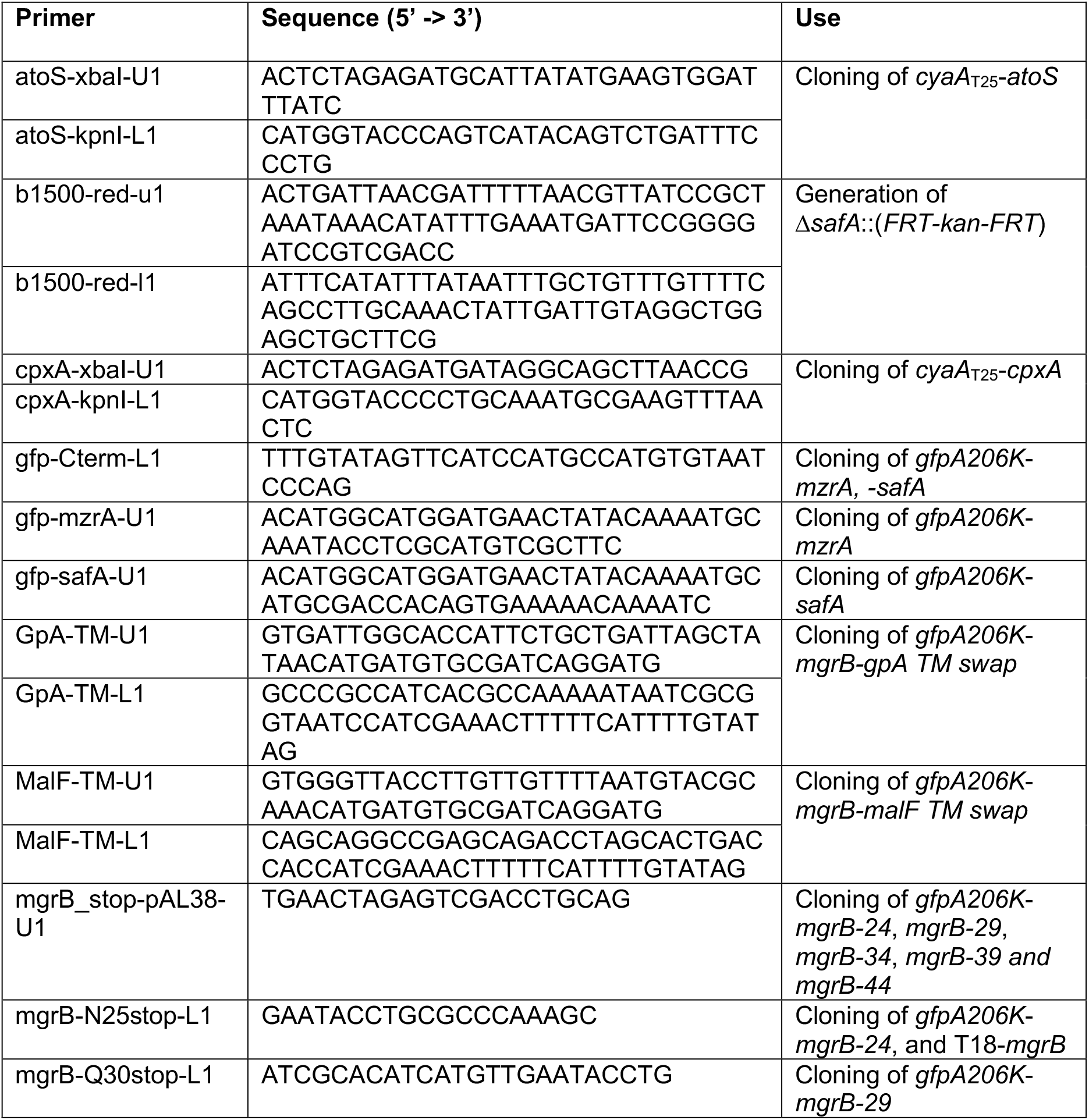
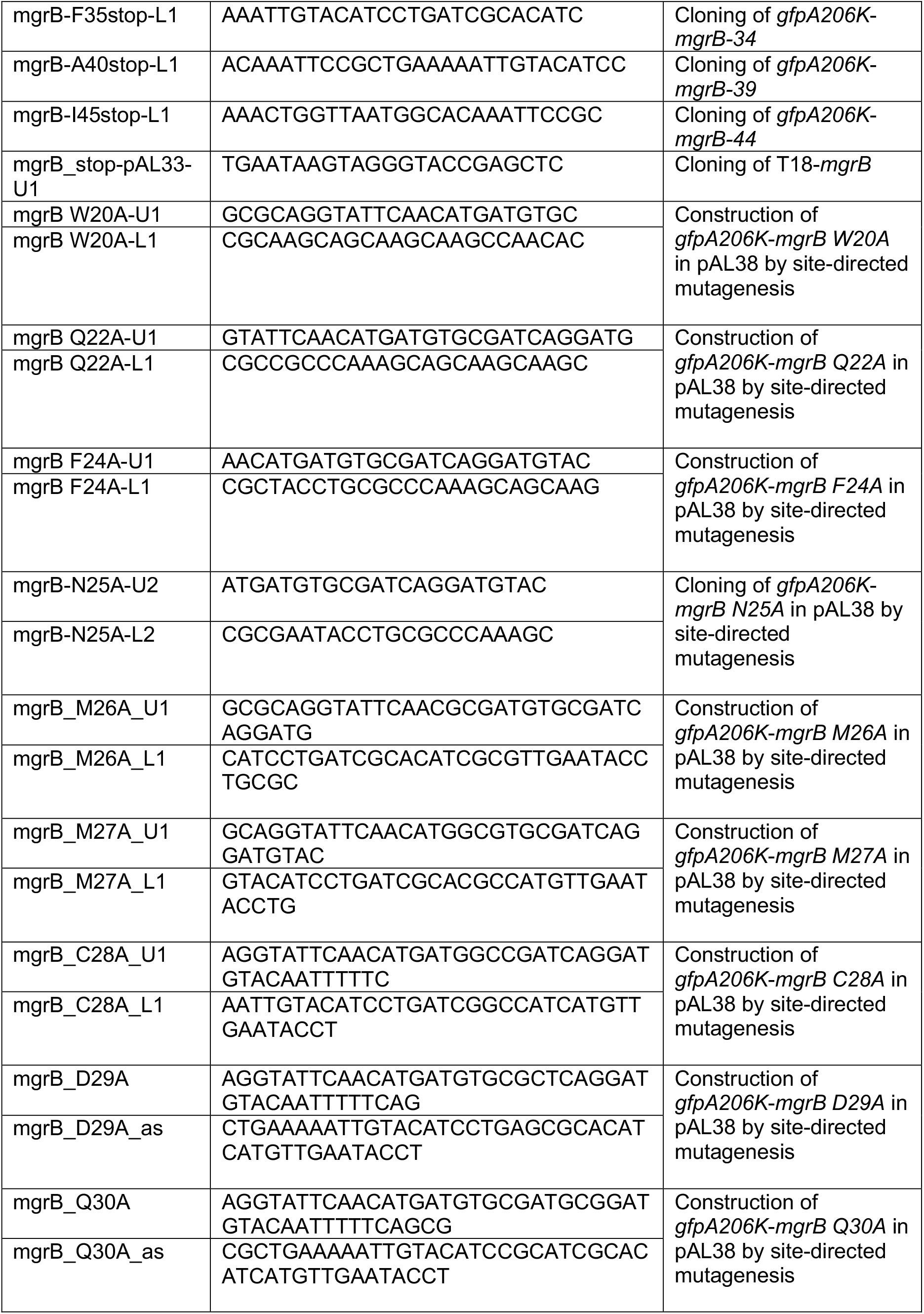
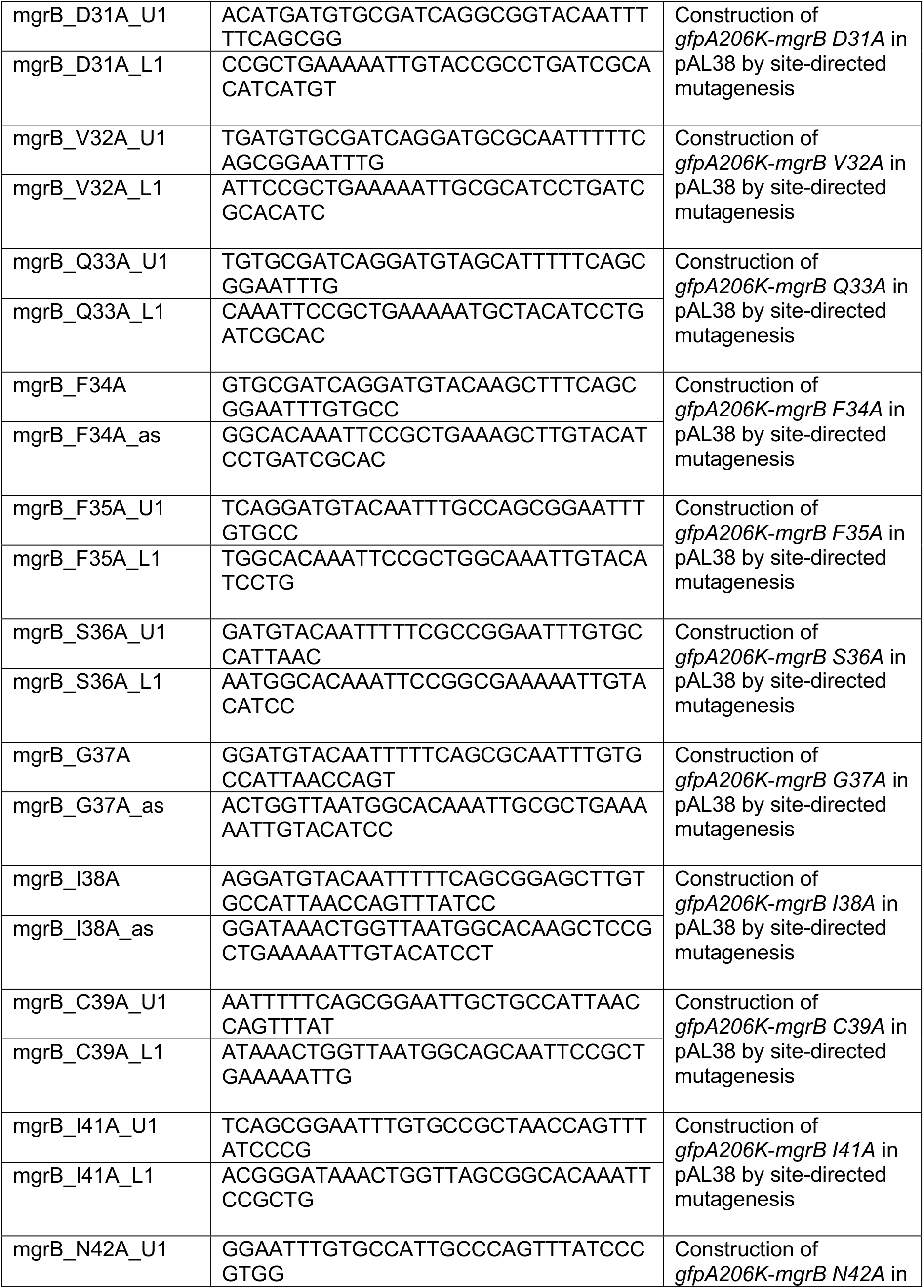
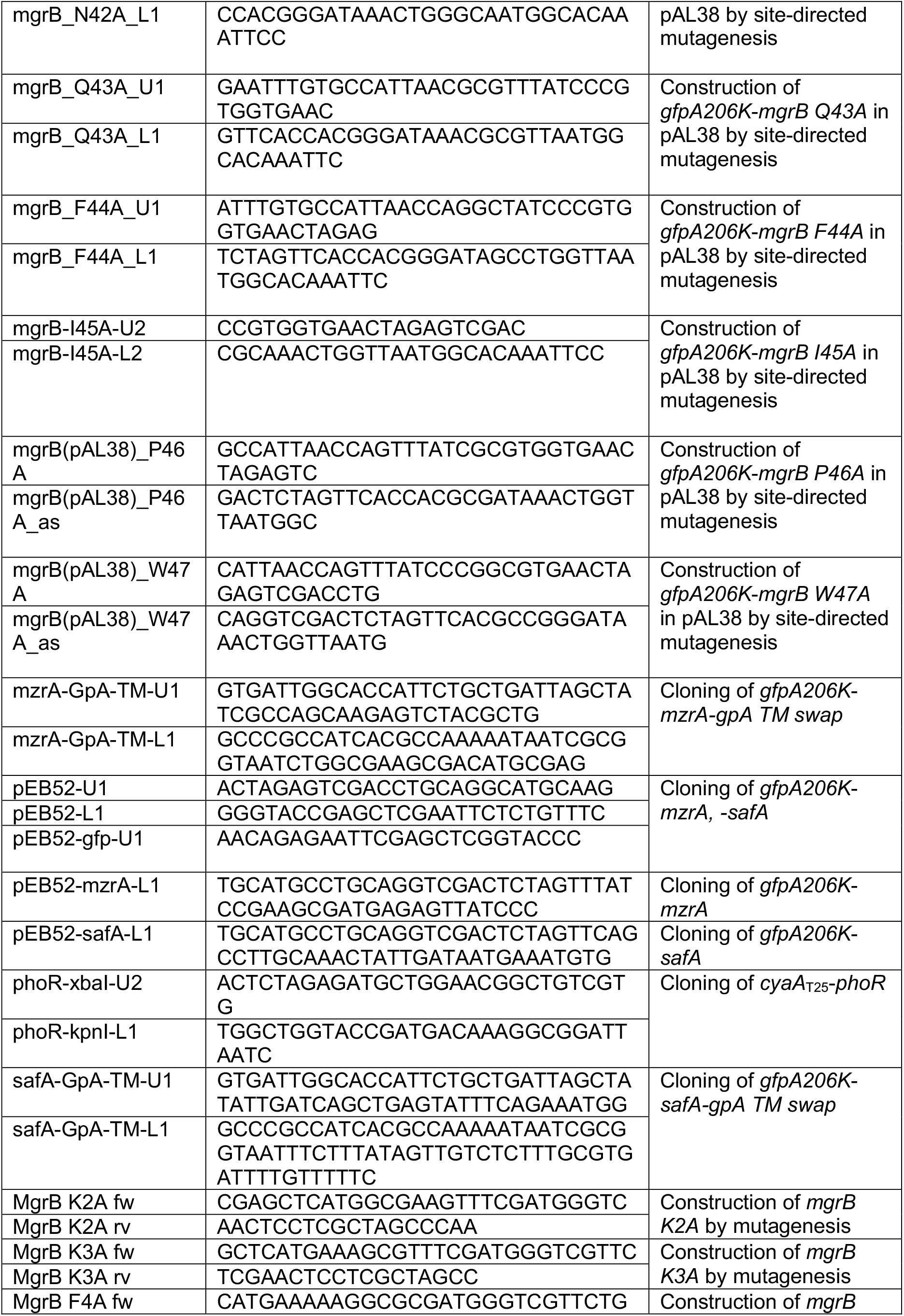
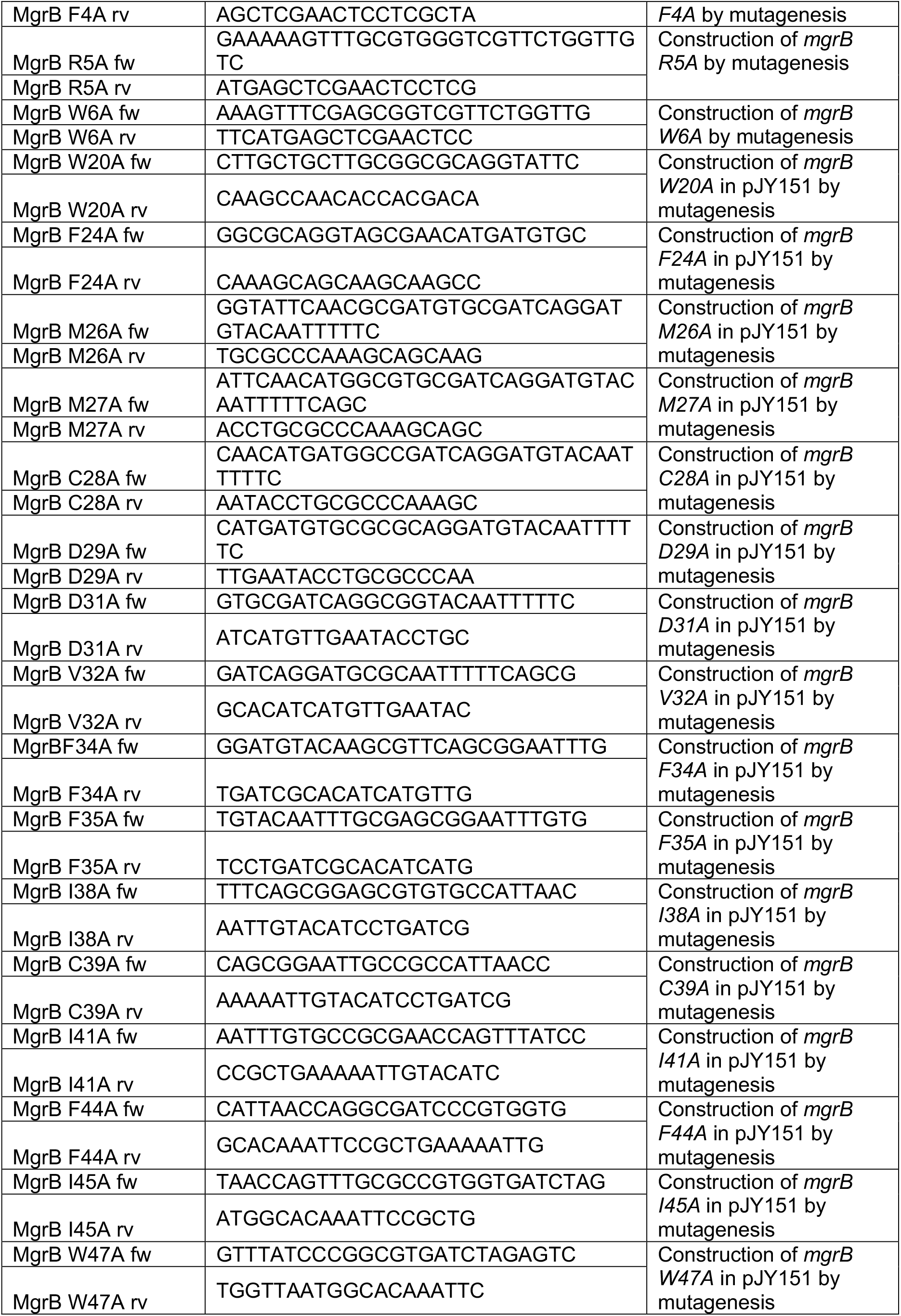
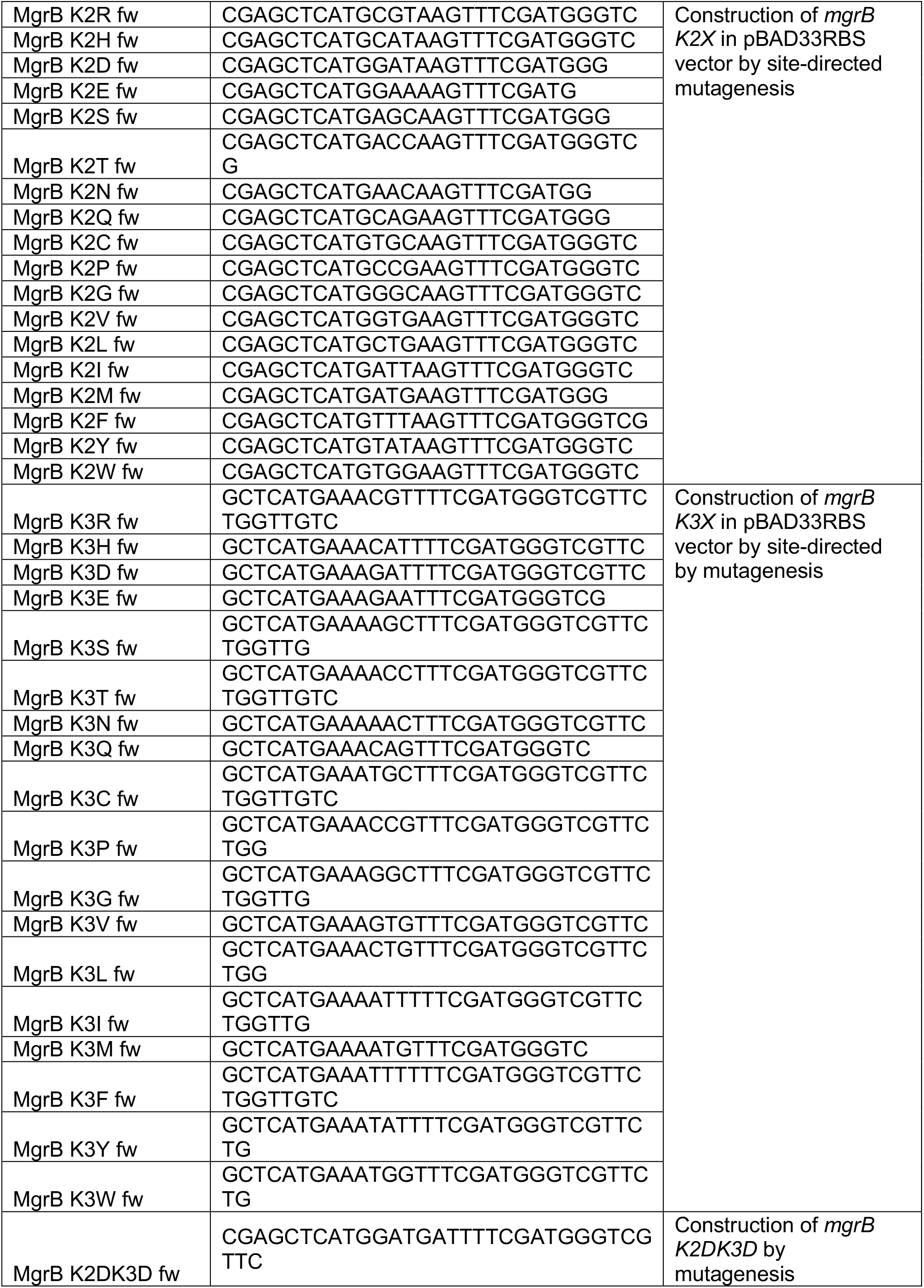
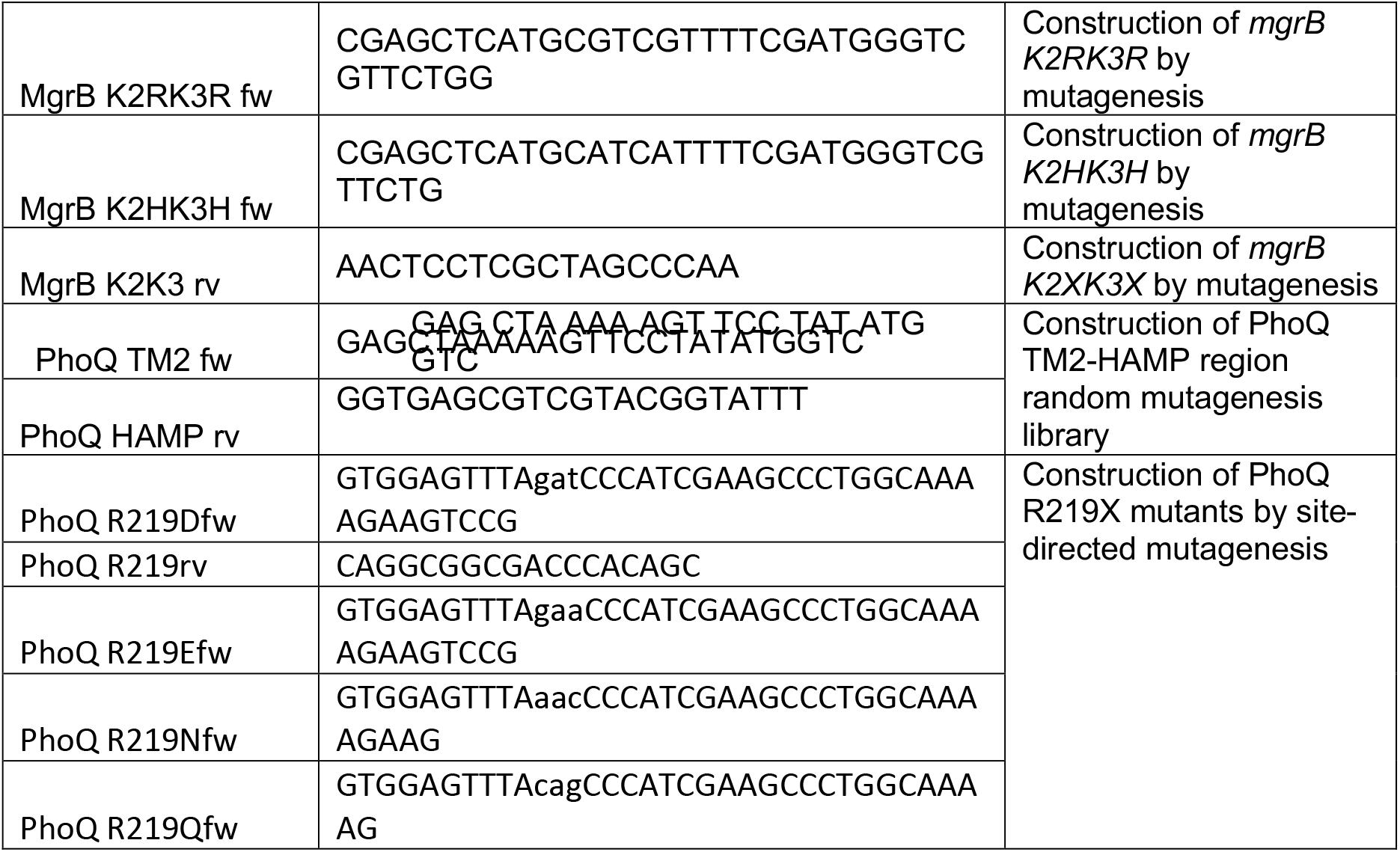
Primers

### Supplementary Figures

**S1 Fig.**
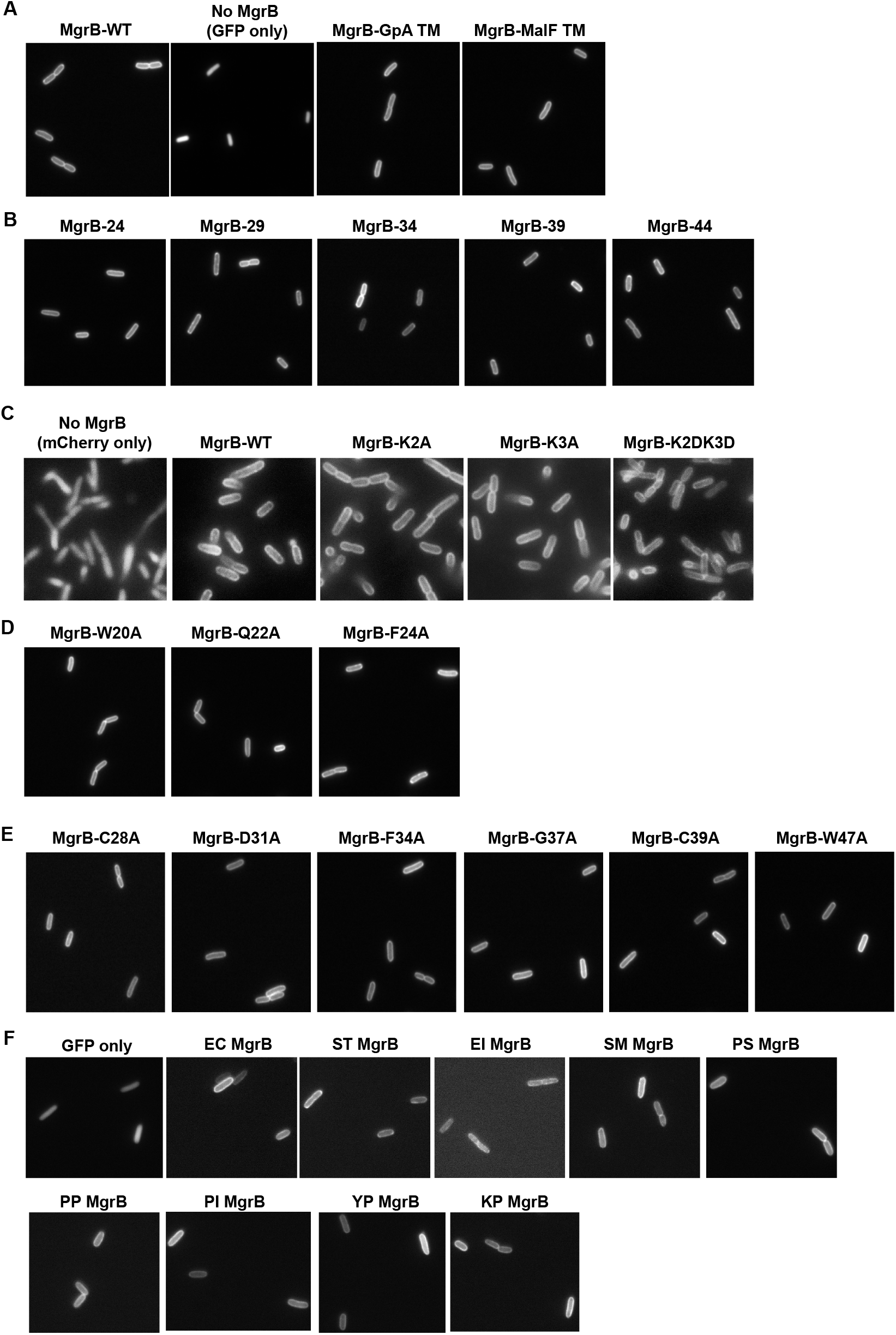
Visualization of cellular localization of MgrB and its variants by microscopy. **A**, Representative fluorescence microscopy images of cells expressing wild-type MgrB, GFP-only, and MgrB variants where the transmembrane region is swapped with transmembrane domain from either human GpA or MalF, respectively. **B**, Representative green fluorescence images of cells expressing C-terminal truncations of MgrB – MgrB-24 (TM only construct, no periplasmic domain), MgrB-29, MgrB-34, MgrB-39 and MgrB-44. **C**, Representative fluorescence microscopy images of cells expressing mCherry-only, mCherry tagged wild-type MgrB and MgrB variants with indicated mutations. **D**, Representative green fluorescence images of cells expressing TM region single Ala mutants of MgrB – W20A, Q22A and F24A. **E**, Representative green fluorescence images of cells expressing periplasmic single Ala mutants of MgrB – C28A, D31A, F34A, G37A, C39A and W47A. *E. coli* MG1655 with *mgrB* deletion is the strain background in C and AML67 is the strain background in A, B, D and E. **F**, Representative green fluorescence images of cells expressing MgrB orthologs (EC- *E. coli*, ST- *Salmonella enterica serovar typhimurium*, EL- *Enterobacter ludwigii*, SM- *Serratia sp.MYb239*, PS- *Providencia stuartii*, PP- *Proteus penneri*, PL- *Photorhabdus laumondii*, YP- *Yesinia pestis* and KP- *Klebsiella pneumoniae*). Inducer was not added as the leaky expression from the *trc* promoter resulted in sufficient levels of protein except for constructs in panel C, where 10 μM IPTG was added.

**S2 Fig.**
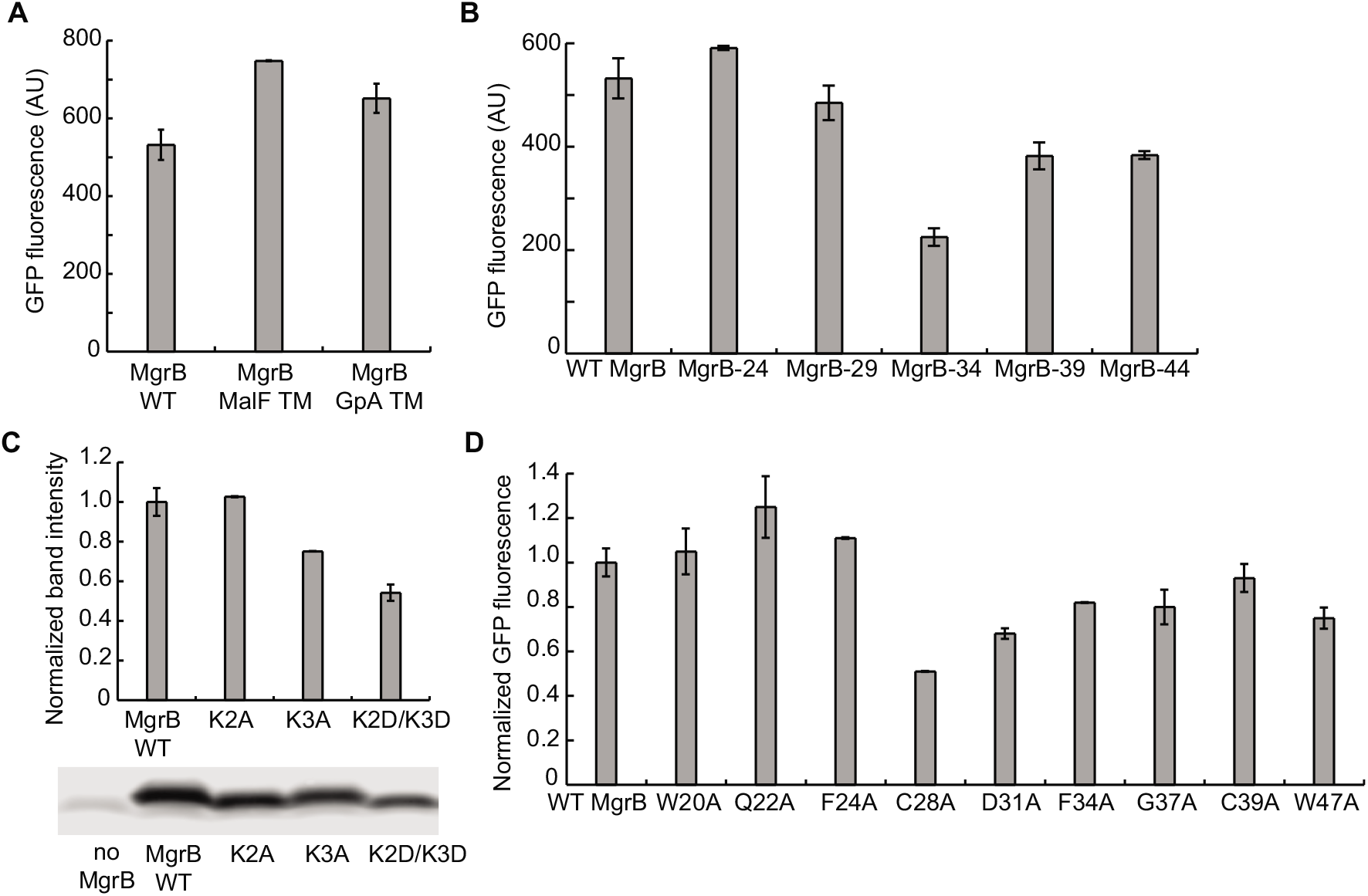
GFP-MgrB expression levels for MgrB variants relative to wild type. GFP-MgrB expression levels for cells expressing (**A**) MgrB TM hybrids, (**B**) MgrB truncation mutants and (**D**) the site-directed single alanine mutants were imaged by fluorescence microscopy and GFP fluorescence levels were quantified (see Methods for details). Data represent averages and ranges for two independent experiments comprised of at least 50 cells each, AU = Arbitrary Units. **C**, The expression level of the FLAG tagged wild-type and MgrB mutants were monitored by western blot. A representative western blot is below. The band intensities were quantified using ImageJ and normalized to wild-type MgrB after taking account of total protein amount in each sample. Data represent averages of three independent experiments and error bars show SDs.

**S3 Fig.**
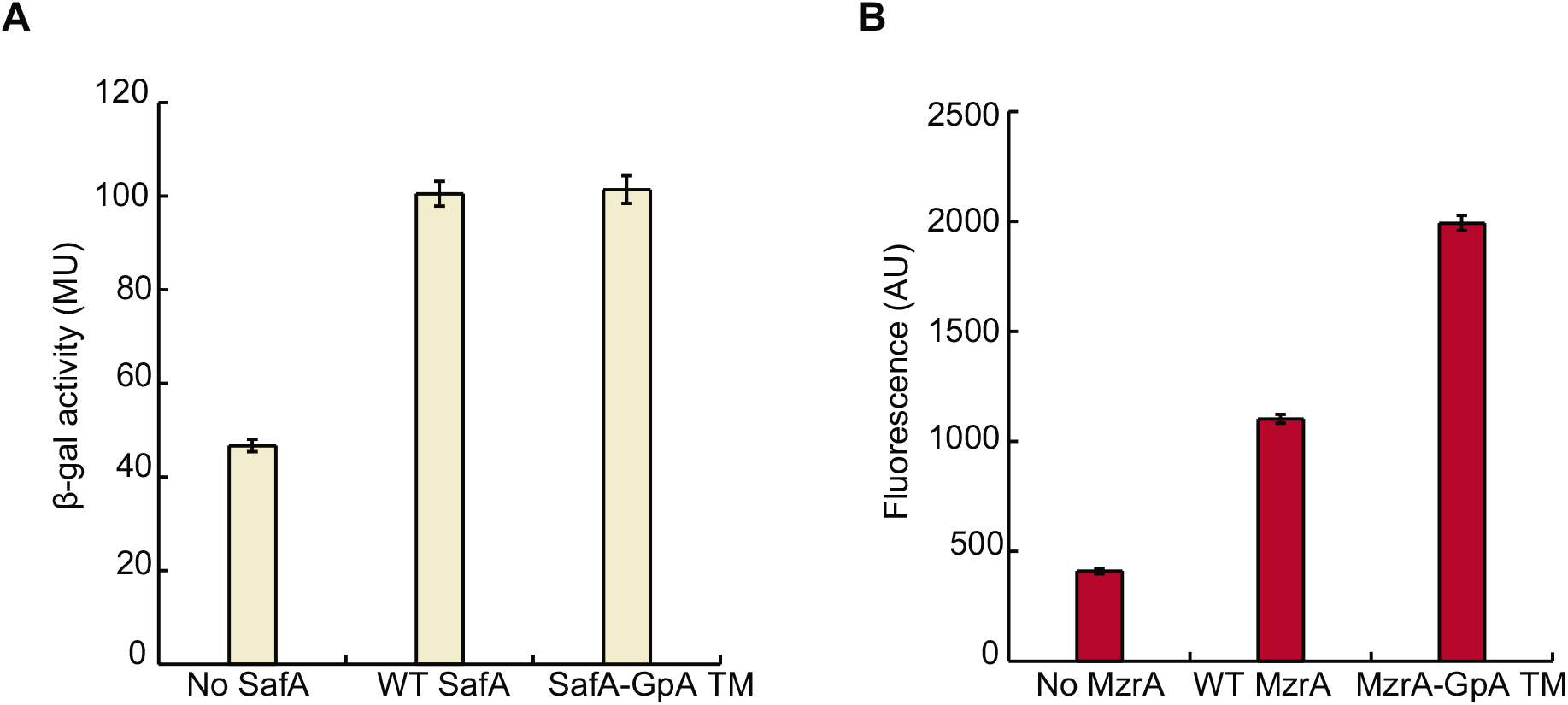
Transmembrane sequences of histidine kinase regulators, SafA and MzrA, are dispensable for their function. **A**, Activities of wild-type SafA and its variant where the transmembrane region is swapped with transmembrane domain from human glycophorin A (GpA) measured as a function of *β*-galactosidase levels in a Δ*safA* strain containing PhoQ/PhoP-regulated transcriptional reporter P*_mgtA_-lacZ* (SAM76), MU = Miller Units. **B**, Activities of wild-type MzrA and its variant where the transmembrane region is swapped with transmembrane domain from human GpA measured as a function of red fluorescence levels in a Δ*mzrA* strain containing EnvZ/OmpR-regulated transcriptional reporter P*_ompc_-mCherry* (SAM72), AU = Arbitrary Units. Inducer (IPTG) was not added as the leaky expression from the *trc* promoter resulted in sufficient levels of protein. Data represent averages and ranges for two independent experiments.

**S4 Fig.**
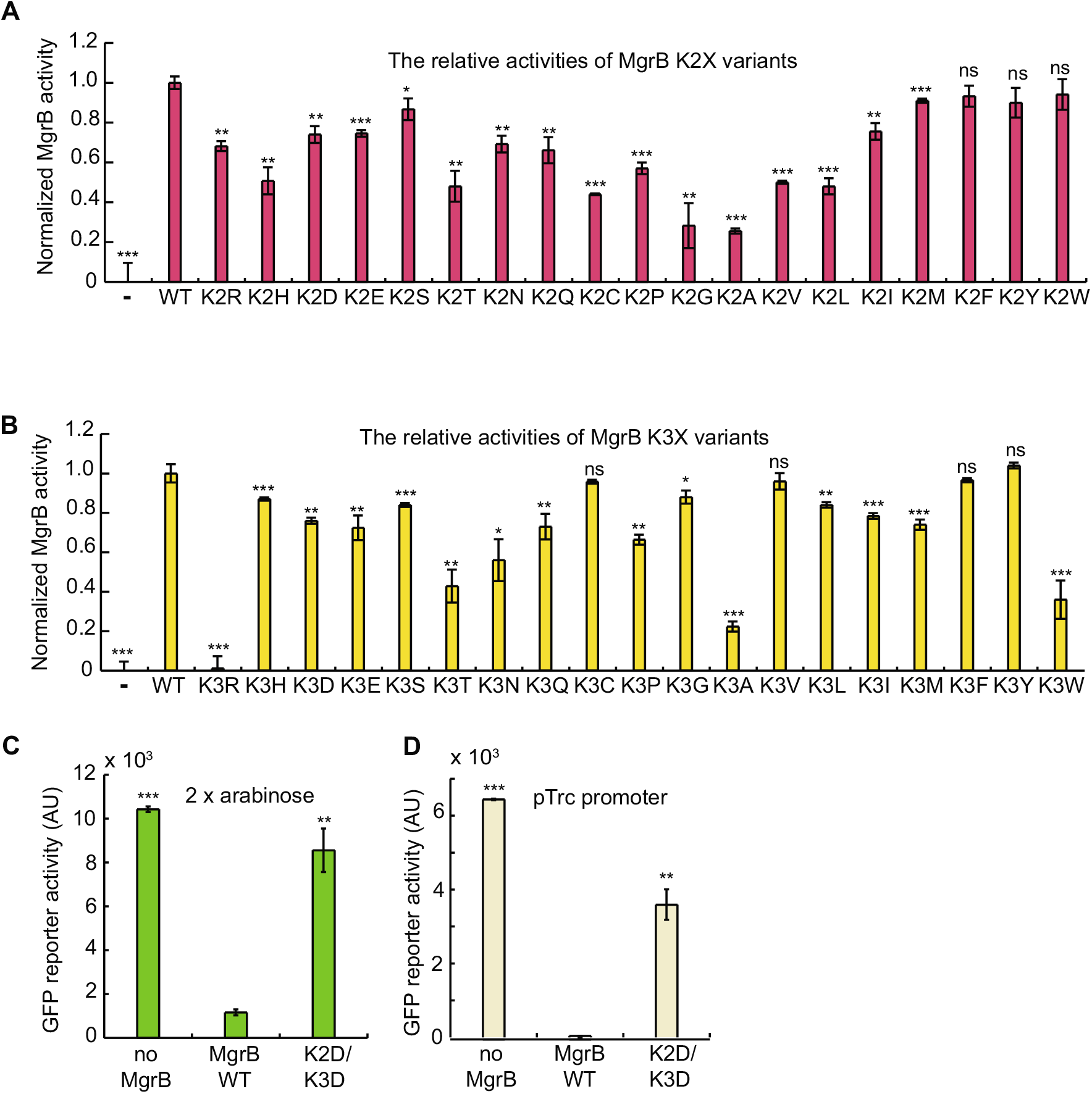
The activities of MgrB variants with mutations in the cytosolic region. The normalized activities of MgrB mutants with single alanine substitutions at K2 (**A**) and K3 (**B**) were tested using the PhoQ/PhoP-regulated transcriptional reporter P*_mgtLA_-gfp* in a Δ*mgrB* strain (see Materials and methods for the definition of normalized activity). MgrB mutants were constructed in a pBAD33 vector and their expression was induced with 0.008% arabinose. The activities of MgrB K2DK3D were tested as in **A** but with double the amount of arabinose inducer (0.016%) (**C**) or with a different plasmid expression construct (**D**). Specifically, in **D**, MgrB variants were expressed from the basal activity of the *trc* promoter (i.e. without inducer) in the plasmid pTrc99a. Mutants with normalized MgrB activity less than 0.4 were selected for further analysis. Data points represent averages of at least three independent experiments and error bars show SDs. The statistical significances of changes in reporter activity or normalized MgrB activity were calculated with *t* tests by comparing to wild type and indicated with asterisks (*** *P* ≤ 0.001, ** *P* ≤ 0.01, * *P* ≤ 0.05, ns = not significant).

**S5 Fig.**
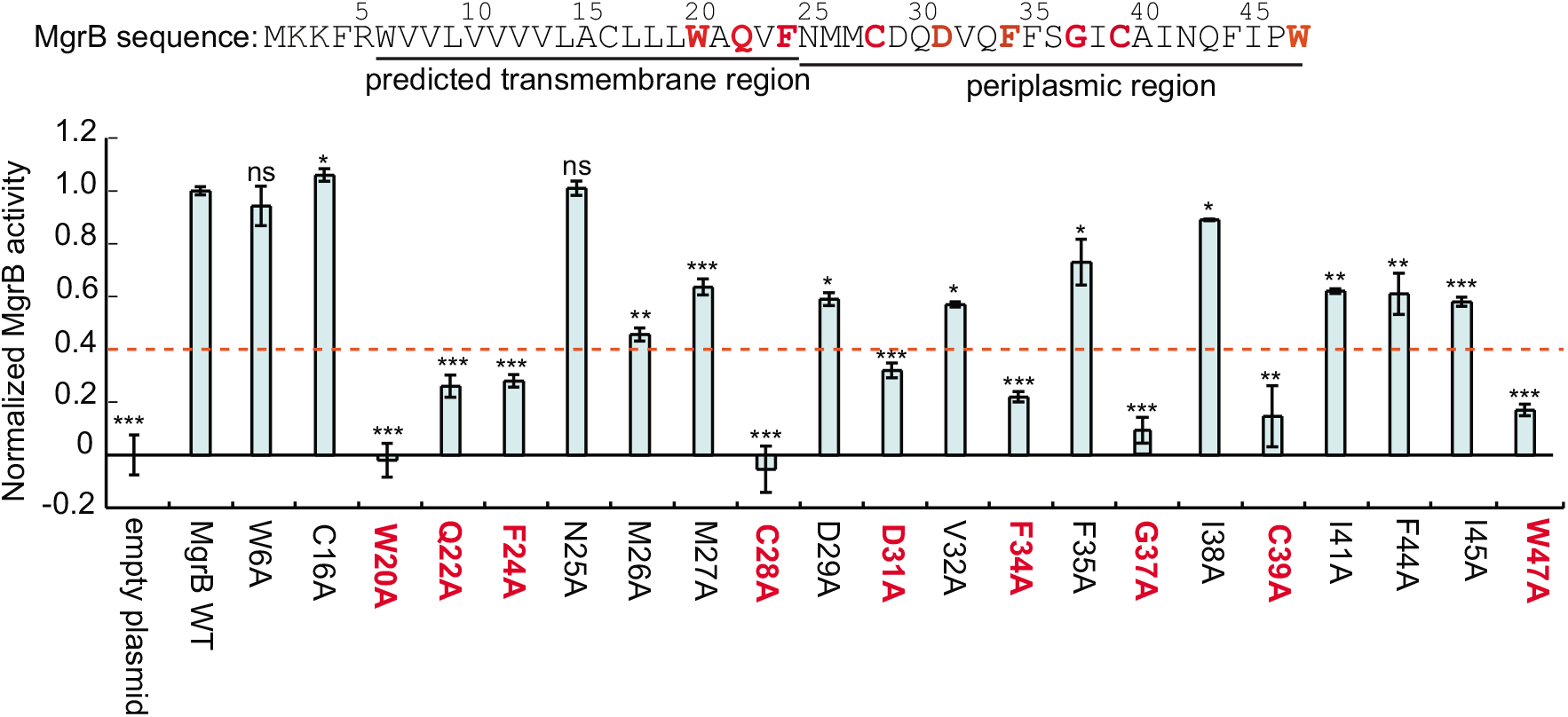
The normalized activities of MgrB variants with alanine substitutions at indicated positions. The activities of MgrB mutants with single alanine substitution were tested using PhoQ/PhoP-regulated transcriptional reporter P*_mgtLA_-gfp* in a Δ*mgrB* strain. MgrB mutants were constructed in a pBAD33 vector and their expression was induced with 0.008% arabinose. The normalized activity was calculated and plotted (see Materials and methods for the definition of normalized activity). Mutants with normalized MgrB activity less than 0.4 (as indicated by the red dashed line) were selected for further analysis. All data points represent averages of at least three independent experiments and error bars show SDs. The statistical significances of changes in normalized MgrB activity were calculated with *t* tests by comparing to wild type and indicated with asterisks (*** *P* ≤ 0.001, ** *P* ≤ 0.01, * *P* ≤ 0.05, ns = not significant).

**S6 Fig.**
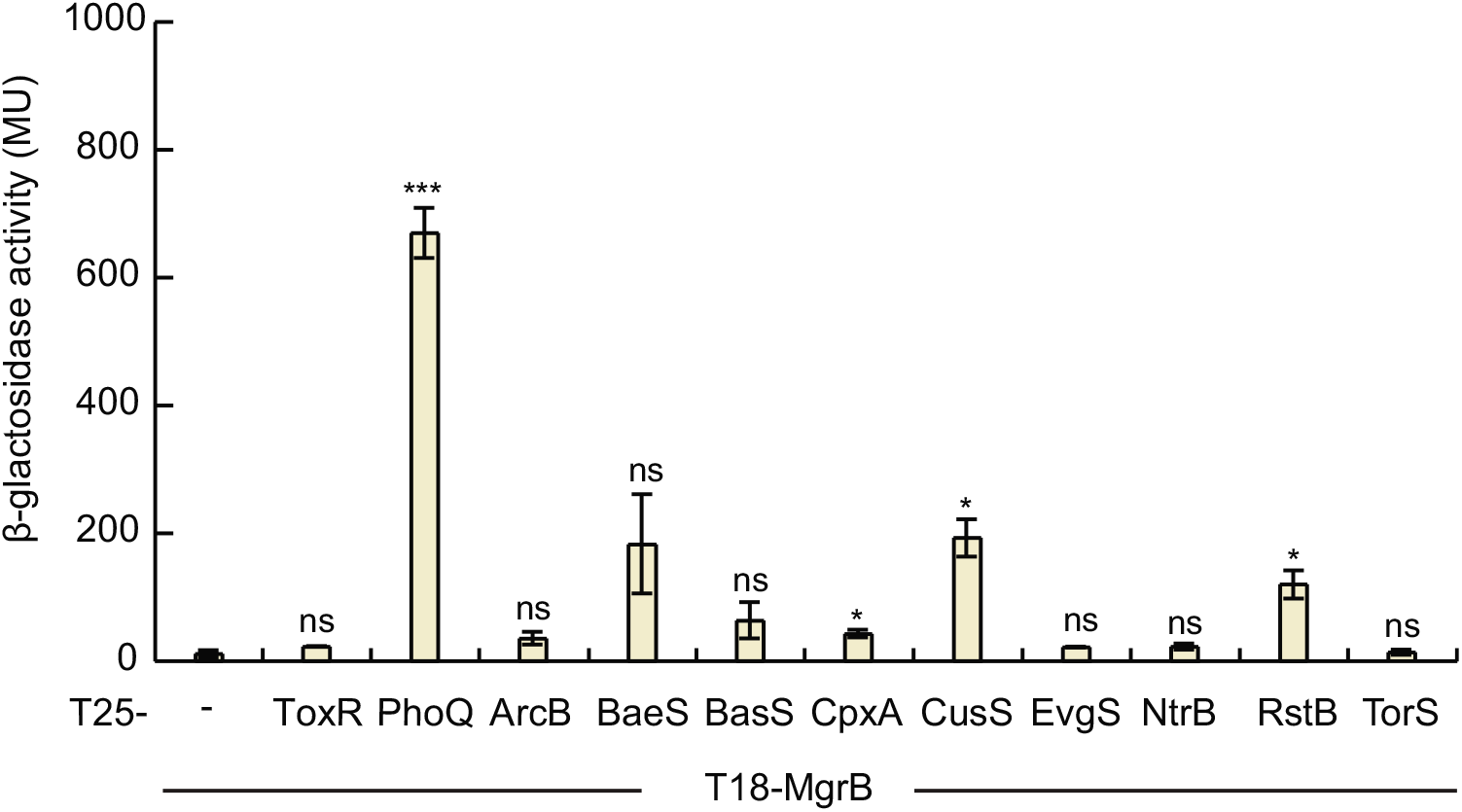
MgrB interacts with other histidine kinases in a bacterial two-hybrid assay. Bacterial two-hybrid (BACTH) assays with cells expressing protein fusions to the T18 and T25 subunits of *B. pertussis* adenylyl cyclase. *β*-gal expression from the *lac* promoter was monitored as a measure of interactions between T18-MgrB and T25 fusion with either ToxR, ArcB, BaeS, CpxA, CusS, EvgS, NtrB, RstB and TorS. The statistical significances of changes in reporter activity were calculated with *t* tests by comparing to the negative control (T18-MgrB/T25 only) and indicated with asterisks (*** *P* ≤ 0.001, ** *P* ≤ 0.01, * *P* ≤ 0.05, ns = not significant).

---

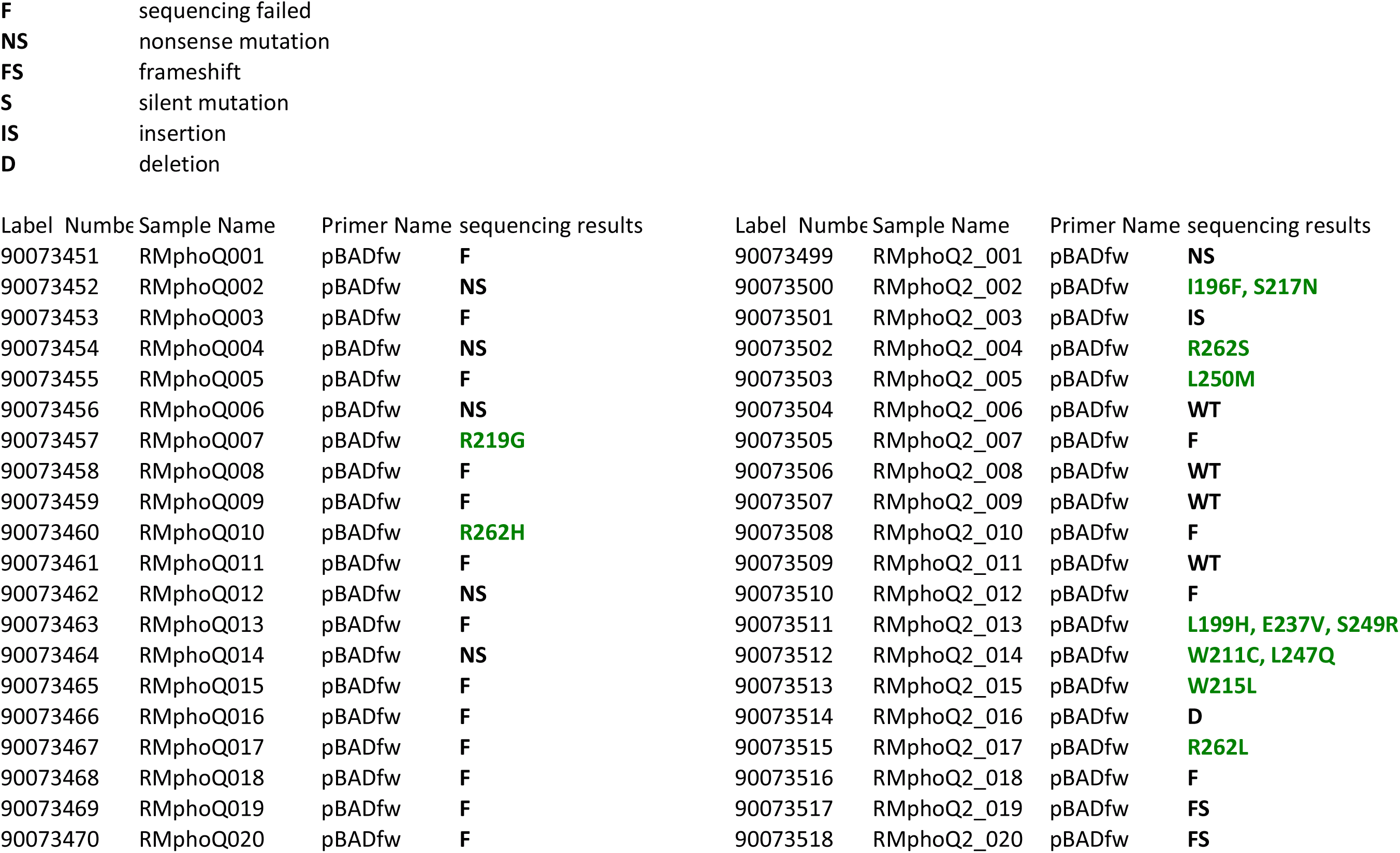
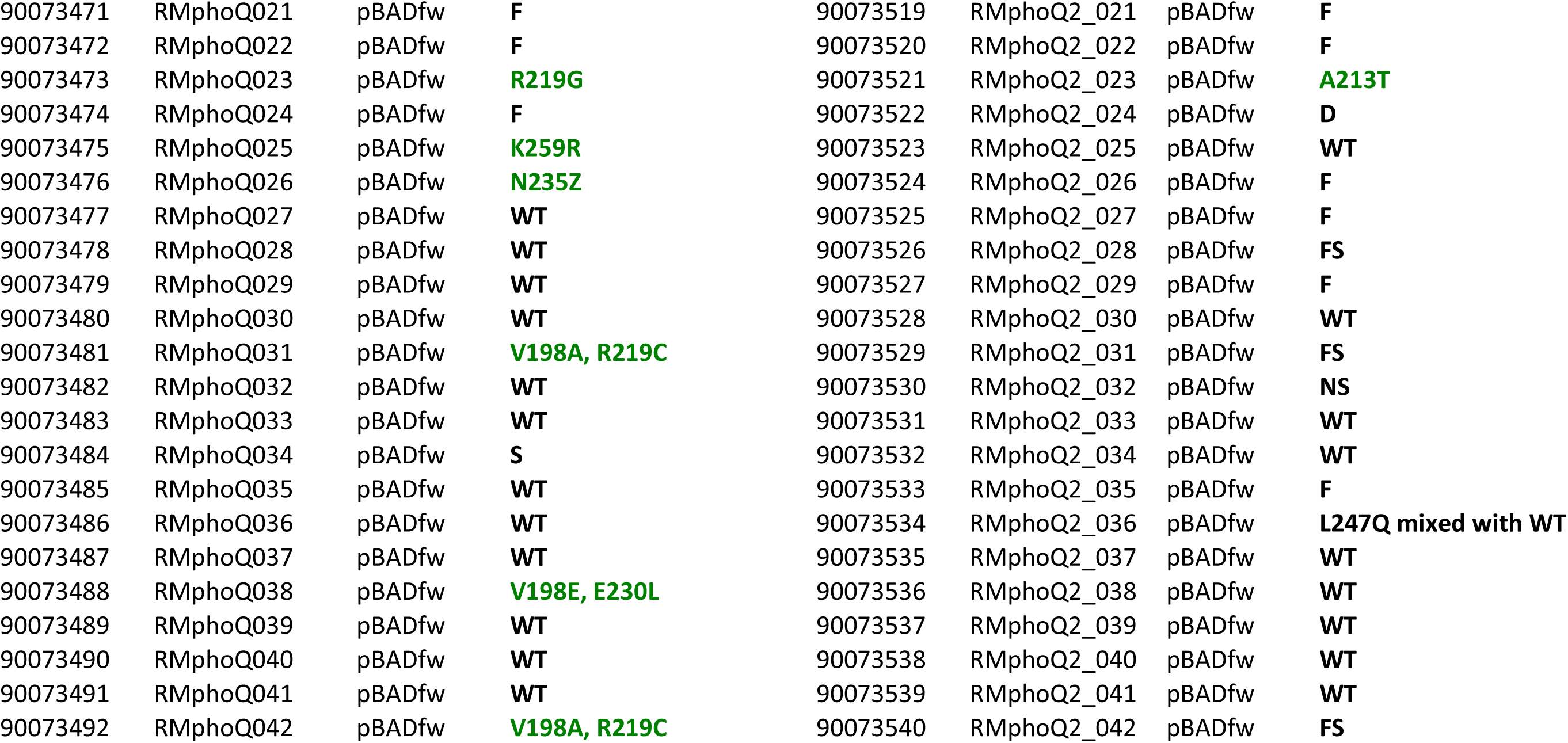
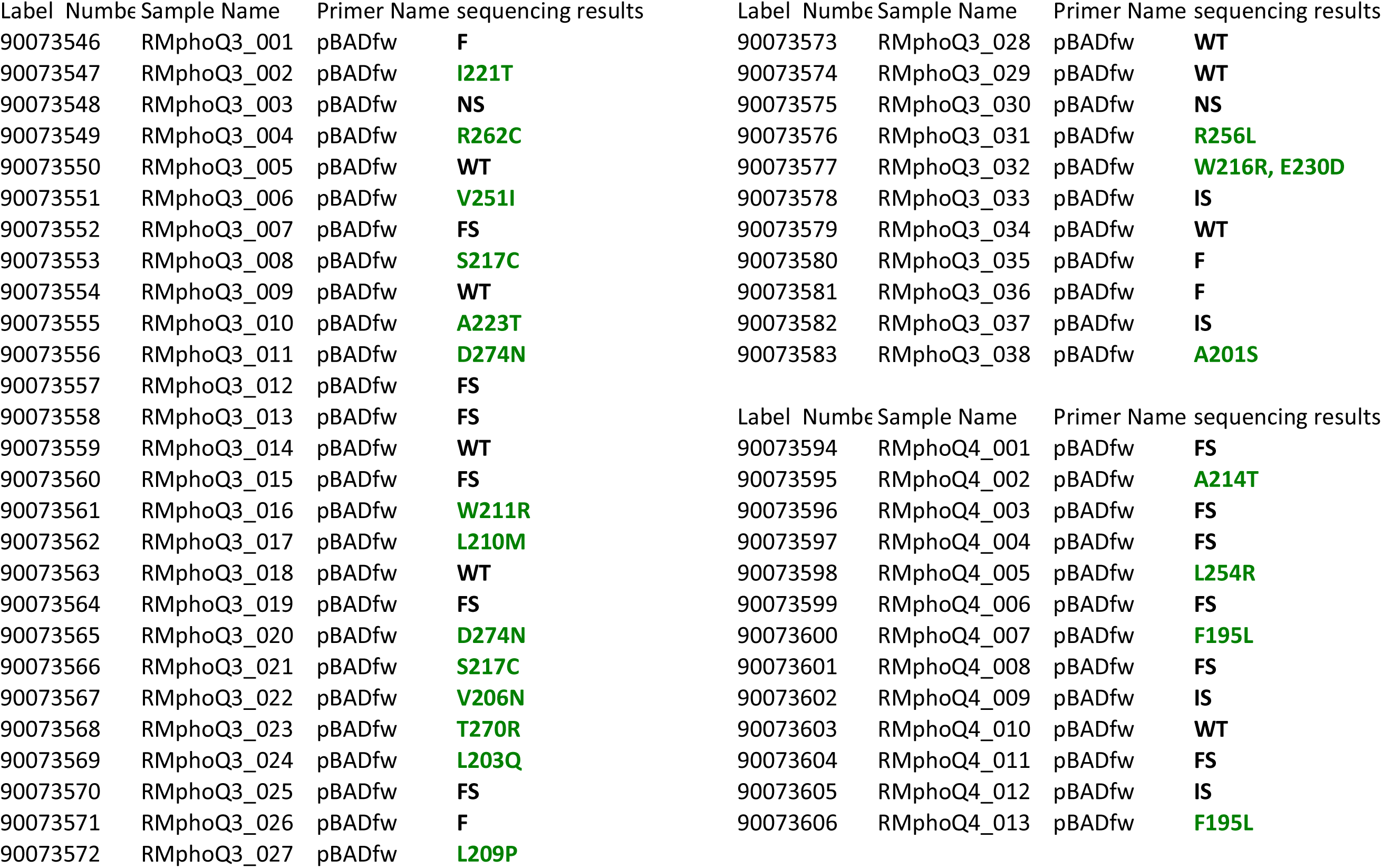

## Notes

### Competing Interest Statement

The authors have declared no competing interest.

## References

1. Duval M, Cossart P. Small bacterial and phagic proteins: an updated view on a rapidly moving field. Curr Opin Microbiol. 2017;39:81–8. doi: 10.1016/j.mib.2017.09.010. PubMed PMID: 29111488.

2. Storz G, Wolf YI, Ramamurthi KS. Small proteins can no longer be ignored. Annu Rev Biochem. 2014;83:753–77. doi: 10.1146/annurev-biochem-070611-102400. PubMed PMID: 24606146; PubMed Central PMCID: PMCPMC4166647.

3. VanOrsdel CE, Kelly JP, Burke BN, Lein CD, Oufiero CE, Sanchez JF, et al. Identifying New Small Proteins in Escherichia coli. Proteomics. 2018;18(10):e1700064. doi: 10.1002/pmic.201700064. PubMed PMID: 29645342; PubMed Central PMCID: PMCPMC6001520.

4. Weaver J, Mohammad F, Buskirk AR, Storz G. Identifying Small Proteins by Ribosome Profiling with Stalled Initiation Complexes. MBio. 2019;10(2). doi: 10.1128/mBio.02819-18. PubMed PMID: 30837344; PubMed Central PMCID: PMCPMC6401488.

5. Nakahigashi K, Takai Y, Kimura M, Abe N, Nakayashiki T, Shiwa Y, et al. Comprehensive identification of translation start sites by tetracycline-inhibited ribosome profiling. DNA Res. 2016;23(3):193–201. doi: 10.1093/dnares/dsw008. PubMed PMID: 27013550; PubMed Central PMCID: PMCPMC4909307.

6. Carol Smith JGC, Jing Wang, Keith M. Derbyshire, Todd A. Gray, Joseph T. Wade. Pervasive translation in Mycobacterium tuberculosis. bioRxiv. 2019. doi: https://doi.org/10.1101/665208.

7. Sberro H, Fremin BJ, Zlitni S, Edfors F, Greenfield N, Snyder MP, et al. Large-Scale Analyses of Human Microbiomes Reveal Thousands of Small, Novel Genes. Cell. 2019;178(5):1245–59 e14. doi: 10.1016/j.cell.2019.07.016. PubMed PMID: 31402174.

8. Hemm MR, Paul BJ, Schneider TD, Storz G, Rudd KE. Small membrane proteins found by comparative genomics and ribosome binding site models. Mol Microbiol. 2008;70(6):1487–501. doi: 10.1111/j.1365-2958.2008.06495.x. PubMed PMID: 19121005; PubMed Central PMCID: PMCPMC2614699.

9. Stock AM, Robinson VL, Goudreau PN. Two-Component Signal Transduction. Annu Rev Biochem. 2000;69(1):183–215. doi: papers2://publication/doi/10.1146/annurev.biochem.69.1.183.

10. Groisman EA. The Pleiotropic Two-Component Regulatory System PhoP-PhoQ. J Bacteriol. 2001;183(6):1835–42. doi: papers2://publication/doi/10.1128/JB.183.6.1835-1842.2001.

11. Garcia Vescovi E, Soncini FC, Groisman EA. Mg^2+^ as an extracellular signal: environmental regulation of *Salmonella* virulence. Cell. 1996;84(1):165–74. PubMed PMID: 8548821.

12. Prost LR, Daley ME, Le Sage V, Bader MW, Le Moual H, Klevit RE, et al. Activation of the bacterial sensor kinase PhoQ by acidic pH. Mol Cell. 2007;26(2):165–74. doi: 10.1016/j.molcel.2007.03.008. PubMed PMID: 17466620.

13. Bader MW, Sanowar S, Daley ME, Schneider AR, Cho U, Xu W, et al. Recognition of antimicrobial peptides by a bacterial sensor kinase. Cell. 2005;122(3):461–72. doi: 10.1016/j.cell.2005.05.030. PubMed PMID: 16096064.

14. Yuan J, Jin F, Glatter T, Sourjik V. Osmosensing by the bacterial PhoQ/PhoP two-component system. Proc Natl Acad Sci U S A. 2017;114(50):E10792–E8. doi: 10.1073/pnas.1717272114. PubMed PMID: 29183977; PubMed Central PMCID: PMCPMC5740661.

15. Goulian M. Two-component signaling circuit structure and properties. Curr Opin Microbiol. 2010;13(2):184–9. doi: papers2://publication/doi/10.1016/j.mib.2010.01.009.

16. Wang H, Yin X, Wu Orr M, Dambach M, Curtis R, Storz G. Increasing intracellular magnesium levels with the 31-amino acid MgtS protein. Proc Natl Acad Sci U S A. 2017;114(22):5689–94. doi: 10.1073/pnas.1703415114. PubMed PMID: 28512220; PubMed Central PMCID: PMCPMC5465934.

17. Alix E, Blanc-Potard AB. Peptide-assisted degradation of the Salmonella MgtC virulence factor. EMBO J. 2008;27(3):546–57. doi: 10.1038/sj.emboj.7601983. PubMed PMID: 18200043; PubMed Central PMCID: PMCPMC2241655.

18. Lippa AM, Goulian M. Feedback inhibition in the PhoQ/PhoP signaling system by a membrane peptide. PLoS Genet. 2009;5(12):e1000788. doi: 10.1371/journal.pgen.1000788. PubMed PMID: 20041203; PubMed Central PMCID: PMCPMC2789325.

19. Salazar ME, Podgornaia AI, Laub MT. The small membrane protein MgrB regulates PhoQ bifunctionality to control PhoP target gene expression dynamics. Mol Microbiol. 2016. doi: 10.1111/mmi.13471. PubMed PMID: 27447896.

20. Lippa AM, Goulian M. Perturbation of the oxidizing environment of the periplasm stimulates the PhoQ/PhoP system in Escherichia coli. J Bacteriol. 2012;194(6):1457–63. doi: 10.1128/JB.06055-11. PubMed PMID: 22267510; PubMed Central PMCID: PMCPMC3294871.

21. Yadavalli SS, Carey JN, Leibman RS, Chen AI, Stern AM, Roggiani M, et al. Antimicrobial peptides trigger a division block in Escherichia coli through stimulation of a signalling system. Nat Commun. 2016;7:12340. doi: 10.1038/ncomms12340. PubMed PMID: 27471053; PubMed Central PMCID: PMCPMC4974570.

22. Kidd TJ, Mills G, Sa-Pessoa J, Dumigan A, Frank CG, Insua JL, et al. A Klebsiella pneumoniae antibiotic resistance mechanism that subdues host defences and promotes virulence. EMBO Mol Med. 2017;9(4):430–47. doi: 10.15252/emmm.201607336. PubMed PMID: 28202493; PubMed Central PMCID: PMCPMC5376759.

23. Cannatelli A, Giani T, D’Andrea MM, Di Pilato V, Arena F, Conte V, et al. MgrB inactivation is a common mechanism of colistin resistance in KPC-producing Klebsiella pneumoniae of clinical origin. Antimicrob Agents Chemother. 2014;58(10):5696–703. doi: papers2://publication/doi/10.1128/AAC.03110-14.

24. Olaitan AO, Morand S, Rolain JM. Mechanisms of polymyxin resistance: acquired and intrinsic resistance in bacteria. Front Microbiol. 2014;5:643. Epub 2014/12/17. doi: 10.3389/fmicb.2014.00643. PubMed PMID: 25505462; PubMed Central PMCID: PMCPMC4244539.

25. Poirel L, Jayol A, Bontron S, Villegas MV, Ozdamar M, Turkoglu S, et al. The mgrB gene as a key target for acquired resistance to colistin in Klebsiella pneumoniae. J Antimicrob Chemother. 2015;70(1):75–80. doi: 10.1093/jac/dku323. PubMed PMID: 25190723.

26. Eguchi Y, Itou J, Yamane M, Demizu R, Yamato F, Okada A, et al. B1500, a small membrane protein, connects the two-component systems EvgS/EvgA and PhoQ/PhoP in Escherichia coli. Proc Natl Acad Sci USA. 2007;104(47):18712–7. doi: papers2://publication/doi/10.1073/pnas.0705768104.

27. Eguchi Y, Ishii E, Hata K, Utsumi R. Regulation of acid resistance by connectors of two-component signal transduction systems in Escherichia coli. J Bacteriol. 2011;193(5):1222–8. doi: 10.1128/JB.01124-10. PubMed PMID: 21193607; PubMed Central PMCID: PMCPMC3067605.

28. Roggiani M, Yadavalli SS, Goulian M. Natural variation of a sensor kinase controlling a conserved stress response pathway in Escherichia coli. PLoS Genet. 2017;13(11):e1007101. Epub 2017/11/16. doi: 10.1371/journal.pgen.1007101. PubMed PMID: 29140975; PubMed Central PMCID: PMCPMC5706723.

29. Burton NA, Johnson MD, Antczak P, Robinson A, Lund PA. Novel aspects of the acid response network of E. coli K-12 are revealed by a study of transcriptional dynamics. Journal of Molecular Biology. 2010. doi: papers2://publication/uuid/EE91AF7A-5760-40B4-B549-BE44E24F43E7.

30. Gerken H, Charlson ES, Cicirelli EM, Kenney LJ, Misra R. MzrA: a novel modulator of the EnvZ/OmpR two-component regulon. Mol Microbiol. 2009;72(6):1408–22. doi: papers2://publication/doi/10.1111/j.1365-2958.2009.06728.x.

31. Ishii E, Eguchi Y, Utsumi R. Mechanism of activation of PhoQ/PhoP two-component signal transduction by SafA, an auxiliary protein of PhoQ histidine kinase in Escherichia coli. Biosci Biotechnol Biochem. 2013;77(4):814–9. doi: papers2://publication/doi/10.1271/bbb.120970.

32. Gerken H, Misra R. MzrA-EnvZ interactions in the periplasm influence the EnvZ/OmpR two-component regulon. J Bacteriol. 2010. doi: papers2://publication/uuid/0796C281-CF57-426D-AB5B-52D4AAA89C30.

33. Leeds JA, Beckwith J. Lambda repressor N-terminal DNA-binding domain as an assay for protein transmembrane segment interactions in vivo. J Mol Biol. 1998;280(5):799–810. Epub 1998/07/22. doi: 10.1006/jmbi.1998.1893. PubMed PMID: 9671551.

34. Eguchi Y, Ishii E, Yamane M, Utsumi R. The connector SafA interacts with the multi-sensing domain of PhoQ in Escherichia coli. Mol Microbiol. 2012;85(2):299–313. doi: 10.1111/j.1365-2958.2012.08114.x. PubMed PMID: 22651704.

35. Sampson BA, Misra R, Benson SA. Identification and characterization of a new gene of Escherichia coli K-12 involved in outer membrane permeability. Genetics. 1989;122(3):491–501. PubMed PMID: 2547691; PubMed Central PMCID: PMCPMC1203724.

36. Battesti A, Bouveret E. The bacterial two-hybrid system based on adenylate cyclase reconstitution in Escherichia coli. Methods. 2012;58(4):325–34. doi: papers2://publication/doi/10.1016/j.ymeth.2012.07.018.

37. Karimova G, Pidoux J, Ullmann A, Ladant D. A bacterial two-hybrid system based on a reconstituted signal transduction pathway. Proc Natl Acad Sci U S A. 1998;95(10):5752–6. Epub 1998/05/20. PubMed PMID: 9576956; PubMed Central PMCID: PMCPMC20451.

38. Kentner D, Sourjik V. Dynamic map of protein interactions in the Escherichia coli chemotaxis pathway. Mol Syst Biol. 2009;5:238. doi: 10.1038/msb.2008.77. PubMed PMID: 19156130; PubMed Central PMCID: PMCPMC2644175.

39. Sourjik V, Vaknin A, Shimizu TS, Berg HC. In vivo measurement by FRET of pathway activity in bacterial chemotaxis. Methods Enzymol. 2007;423:365–91. doi: 10.1016/S0076-6879(07)23017-4. PubMed PMID: 17609141.

40. Meiresonne NY, van der Ploeg R, Hink MA, den Blaauwen T. Activity-Related Conformational Changes in d,d-Carboxypeptidases Revealed by In Vivo Periplasmic Forster Resonance Energy Transfer Assay in Escherichia coli. MBio. 2017;8(5). doi: 10.1128/mBio.01089-17. PubMed PMID: 28900026; PubMed Central PMCID: PMCPMC5596342.

41. Fink A, Sal-Man N, Gerber D, Shai Y. Transmembrane domains interactions within the membrane milieu: principles, advances and challenges. Biochim Biophys Acta. 2012;1818(4):974–83. Epub 2011/12/14. doi: 10.1016/j.bbamem.2011.11.029. PubMed PMID: 22155642.

42. von Heijne G. Membrane-protein topology. Nature Reviews Molecular Cell Biology. 2006;7:909. doi: 10.1038/nrm2063 https://www.nature.com/articles/nrm2063-supplementary-information.

43. Ulmschneider MB, Sansom MS. Amino acid distributions in integral membrane protein structures. Biochim Biophys Acta. 2001;1512(1):1–14. Epub 2001/05/04. PubMed PMID: 11334619.

44. Matamouros S, Hager KR, Miller SI. HAMP Domain Rotation and Tilting Movements Associated with Signal Transduction in the PhoQ Sensor Kinase. MBio. 2015;6(3):e00616–15. doi: 10.1128/mBio.00616-15. PubMed PMID: 26015499; PubMed Central PMCID: PMCPMC4447245.

45. Yau WM, Wimley WC, Gawrisch K, White SH. The preference of tryptophan for membrane interfaces. Biochemistry. 1998;37(42):14713–8. doi: 10.1021/bi980809c. PubMed PMID: 9778346.

46. de Jesus AJ, Allen TW. The role of tryptophan side chains in membrane protein anchoring and hydrophobic mismatch. Biochim Biophys Acta. 2013;1828(2):864–76. doi: 10.1016/j.bbamem.2012.09.009. PubMed PMID: 22989724.

47. Goldberg SD, Clinthorne GD, Goulian M, DeGrado WF. Transmembrane polar interactions are required for signaling in the Escherichia coli sensor kinase PhoQ. Proc Natl Acad Sci U S A. 2010;107(18):8141–6. doi: 10.1073/pnas.1003166107. PubMed PMID: 20404199; PubMed Central PMCID: PMCPMC2889538.

48. Bhate MP, Molnar KS, Goulian M, DeGrado WF. Signal transduction in histidine kinases: insights from new structures. Structure. 2015;23(6):981–94. Epub 2015/05/20. doi: 10.1016/j.str.2015.04.002. PubMed PMID: 25982528; PubMed Central PMCID: PMCPMC4456306.

49. Bi S, Jin F, Sourjik V. Inverted signaling by bacterial chemotaxis receptors. Nat Commun. 2018;9(1):2927. doi: 10.1038/s41467-018-05335-w. PubMed PMID: 30050034; PubMed Central PMCID: PMCPMC6062612.

50. Olaitan AO, Diene SM, Kempf M, Berrazeg M, Bakour S, Gupta SK, et al. Worldwide emergence of colistin resistance in Klebsiella pneumoniae from healthy humans and patients in Lao PDR, Thailand, Israel, Nigeria and France owing to inactivation of the PhoP/PhoQ regulator mgrB: an epidemiological and molecular study. Int J Antimicrob Agents. 2014;44(6):500–7. Epub 2014/09/30. doi: 10.1016/j.ijantimicag.2014.07.020. PubMed PMID: 25264127.

51. Cheng YH, Lin TL, Pan YJ, Wang YP, Lin YT, Wang JT. Colistin resistance mechanisms in Klebsiella pneumoniae strains from Taiwan. Antimicrob Agents Chemother. 2015;59(5):2909–13. Epub 2015/02/19. doi: 10.1128/AAC.04763-14. PubMed PMID: 25691646; PubMed Central PMCID: PMCPMC4394772.

52. Bakthavatchalam YD, Pragasam AK, Biswas I, Veeraraghavan B. Polymyxin susceptibility testing, interpretative breakpoints and resistance mechanisms: An update. J Glob Antimicrob Resist. 2018;12:124–36. Epub 2017/10/01. doi: 10.1016/j.jgar.2017.09.011. PubMed PMID: 28962863.

53. Nordmann P, Jayol A, Poirel L. A Universal Culture Medium for Screening Polymyxin-Resistant Gram-Negative Isolates. J Clin Microbiol. 2016;54(5):1395–9. Epub 2016/03/18. doi: 10.1128/JCM.00446-16. PubMed PMID: 26984971; PubMed Central PMCID: PMCPMC4844728.

54. Bonura C, Giuffre M, Aleo A, Fasciana T, Di Bernardo F, Stampone T, et al. An Update of the Evolving Epidemic of blaKPC Carrying Klebsiella pneumoniae in Sicily, Italy, 2014: Emergence of Multiple Non-ST258 Clones. PLoS One. 2015;10(7):e0132936. Epub 2015/07/16. doi: 10.1371/journal.pone.0132936. PubMed PMID: 26177547; PubMed Central PMCID: PMCPMC4503429.

55. Haeili M, Javani A, Moradi J, Jafari Z, Feizabadi MM, Babaei E. MgrB Alterations Mediate Colistin Resistance in Klebsiella pneumoniae Isolates from Iran. Front Microbiol. 2017;8:2470. Epub 2018/01/13. doi: 10.3389/fmicb.2017.02470. PubMed PMID: 29326662; PubMed Central PMCID: PMCPMC5741654.

56. Baba T, Ara T, Hasegawa M, Takai Y, Okumura Y, Baba M, et al. Construction of Escherichia coli K-12 in-frame, single-gene knockout mutants: the Keio collection. Mol Syst Biol. 2006;2:2006 0008. doi: 10.1038/msb4100050. PubMed PMID: 16738554; PubMed Central PMCID: PMC1681482.

57. Datsenko KA, Wanner BL. One-step inactivation of chromosomal genes in Escherichia coli K-12 using PCR products. Proc Natl Acad Sci USA. 2000;97(12):6640–5. doi: papers2://publication/doi/10.1073/pnas.120163297.

58. Ochman H, Ajioka JW, Garza D, Hartl DL. Inverse polymerase chain reaction. Biotechnology (N Y). 1990;8(8):759–60. Epub 1990/08/01. PubMed PMID: 1366903.

59. Gibson DG, Young L, Chuang RY, Venter JC, Hutchison CA, 3rd, Smith HO. Enzymatic assembly of DNA molecules up to several hundred kilobases. Nat Methods. 2009;6(5):343–5. Epub 2009/04/14. doi: 10.1038/nmeth.1318. PubMed PMID: 19363495.

60. Besharova O, Suchanek VM, Hartmann R, Drescher K, Sourjik V. Diversification of Gene Expression during Formation of Static Submerged Biofilms by Escherichia coli. Front Microbiol. 2016;7:1568. doi: 10.3389/fmicb.2016.01568. PubMed PMID: 27761132; PubMed Central PMCID: PMCPMC5050211.

61. Miller JH. A Short Course in Bacterial Genetics: CSHL Press; 1992.

62. Thibodeau SA, Fang R, Joung JK. High-throughput beta-galactosidase assay for bacterial cell-based reporter systems. Biotechniques. 2004;36(3):410–5. doi: 10.2144/04363BM07. PubMed PMID: 15038156.

63. Miyashiro T, Goulian M. Stimulus-dependent differential regulation in the Escherichia coli PhoQ PhoP system. Proc Natl Acad Sci USA. 2007;104(41):16305–10. doi: papers2://publication/doi/10.1073/pnas.0700025104.

64. Haldimann A, Wanner BL. Conditional-replication, integration, excision, and retrieval plasmid-host systems for gene structure-function studies of bacteria. J Bacteriol. 2001;183(21):6384–93. Epub 2001/10/10. doi: 10.1128/JB.183.21.6384-6393.2001. PubMed PMID: 11591683; PubMed Central PMCID: PMCPMC100134.

65. Bullock WO, Fernandez JM, Short JM. XL1-Blue: a high efficiency plasmid transforming recA *Escherichia coli* strain with beta-galactosidase selection. BioTechniques. 1987;5:376–9.

66. Cherepanov PP, Wackernagel W. Gene disruption in Escherichia coli: Tc R and Km R cassettes with the option of Flp-catalyzed excision of the antibiotic-resistance determinant. Gene. 1995. doi: papers2://publication/uuid/31A60367-CF87-4BF6-B690-93462755E5ED.

67. Hay AJ, Yang M, Xia X, Liu Z, Hammons J, Fenical W, et al. Calcium Enhances Bile Salt-Dependent Virulence Activation in Vibrio cholerae. Infect Immun. 2017;85(1). Epub 2016/11/17. doi: 10.1128/IAI.00707-16. PubMed PMID: 27849180; PubMed Central PMCID: PMCPMC5203667.

68. Amann E, Ochs B, Abel KJ. Tightly regulated tac promoter vectors useful for the expression of unfused and fused proteins in Escherichia coli. Gene. 1988;69(2):301–15. PubMed PMID: 3069586.

