## Supplementary material for "Functional determinants of a small protein controlling a broadly conserved bacterial sensor kinase": PhoQ RM screening

| | |
|---|---|
| **F** | sequencing failed |
| **NS** | nonsense mutation |
| **FS** | frameshift |
| **S** | silent mutation |
| **IS** | insertion |
| **D** | deletion |

| Label Numbe | Sample Name | Primer Name | sequencing results |
|---|---|---|---|
| 90073451 | RMphoQ001 | pBADfw | **F** |
| 90073452 | RMphoQ002 | pBADfw | **NS** |
| 90073453 | RMphoQ003 | pBADfw | **F** |
| 90073454 | RMphoQ004 | pBADfw | **NS** |
| 90073455 | RMphoQ005 | pBADfw | **F** |
| 90073456 | RMphoQ006 | pBADfw | **NS** |
| 90073457 | RMphoQ007 | pBADfw | **R219G** |
| 90073458 | RMphoQ008 | pBADfw | **F** |
| 90073459 | RMphoQ009 | pBADfw | **F** |
| 90073460 | RMphoQ010 | pBADfw | **R262H** |
| 90073461 | RMphoQ011 | pBADfw | **F** |
| 90073462 | RMphoQ012 | pBADfw | **NS** |
| 90073463 | RMphoQ013 | pBADfw | **F** |
| 90073464 | RMphoQ014 | pBADfw | **NS** |
| 90073465 | RMphoQ015 | pBADfw | **F** |
| 90073466 | RMphoQ016 | pBADfw | **F** |
| 90073467 | RMphoQ017 | pBADfw | **F** |
| 90073468 | RMphoQ018 | pBADfw | **F** |
| 90073469 | RMphoQ019 | pBADfw | **F** |
| 90073470 | RMphoQ020 | pBADfw | **F** |

| Label Numbe | Sample Name | Primer Name | sequencing results |
|---|---|---|---|
| 90073499 | RMphoQ2_001 | pBADfw | **NS** |
| 90073500 | RMphoQ2_002 | pBADfw | **I196F, S217N** |
| 90073501 | RMphoQ2_003 | pBADfw | **IS** |
| 90073502 | RMphoQ2_004 | pBADfw | **R262S** |
| 90073503 | RMphoQ2_005 | pBADfw | **L250M** |
| 90073504 | RMphoQ2_006 | pBADfw | **WT** |
| 90073505 | RMphoQ2_007 | pBADfw | **F** |
| 90073506 | RMphoQ2_008 | pBADfw | **WT** |
| 90073507 | RMphoQ2_009 | pBADfw | **WT** |
| 90073508 | RMphoQ2_010 | pBADfw | **F** |
| 90073509 | RMphoQ2_011 | pBADfw | **WT** |
| 90073510 | RMphoQ2_012 | pBADfw | **F** |
| 90073511 | RMphoQ2_013 | pBADfw | **L199H, E237V, S249R** |
| 90073512 | RMphoQ2_014 | pBADfw | **W211C, L247Q** |
| 90073513 | RMphoQ2_015 | pBADfw | **W215L** |
| 90073514 | RMphoQ2_016 | pBADfw | **D** |
| 90073515 | RMphoQ2_017 | pBADfw | **R262L** |
| 90073516 | RMphoQ2_018 | pBADfw | **F** |
| 90073517 | RMphoQ2_019 | pBADfw | **FS** |
| 90073518 | RMphoQ2_020 | pBADfw | **FS** |

| | | | | | | | |
|---|---|---|---|---|---|---|---|
| 90073471 | RMphoQ021 | pBADfw | **F** | 90073519 | RMphoQ2_021 | pBADfw | **F** |
| 90073472 | RMphoQ022 | pBADfw | **F** | 90073520 | RMphoQ2_022 | pBADfw | **F** |
| 90073473 | RMphoQ023 | pBADfw | **R219G** | 90073521 | RMphoQ2_023 | pBADfw | **A213T** |
| 90073474 | RMphoQ024 | pBADfw | **F** | 90073522 | RMphoQ2_024 | pBADfw | **D** |
| 90073475 | RMphoQ025 | pBADfw | **K259R** | 90073523 | RMphoQ2_025 | pBADfw | **WT** |
| 90073476 | RMphoQ026 | pBADfw | **N235Z** | 90073524 | RMphoQ2_026 | pBADfw | **F** |
| 90073477 | RMphoQ027 | pBADfw | **WT** | 90073525 | RMphoQ2_027 | pBADfw | **F** |
| 90073478 | RMphoQ028 | pBADfw | **WT** | 90073526 | RMphoQ2_028 | pBADfw | **FS** |
| 90073479 | RMphoQ029 | pBADfw | **WT** | 90073527 | RMphoQ2_029 | pBADfw | **F** |
| 90073480 | RMphoQ030 | pBADfw | **WT** | 90073528 | RMphoQ2_030 | pBADfw | **WT** |
| 90073481 | RMphoQ031 | pBADfw | **V198A, R219C** | 90073529 | RMphoQ2_031 | pBADfw | **FS** |
| 90073482 | RMphoQ032 | pBADfw | **WT** | 90073530 | RMphoQ2_032 | pBADfw | **NS** |
| 90073483 | RMphoQ033 | pBADfw | **WT** | 90073531 | RMphoQ2_033 | pBADfw | **WT** |
| 90073484 | RMphoQ034 | pBADfw | **S** | 90073532 | RMphoQ2_034 | pBADfw | **WT** |
| 90073485 | RMphoQ035 | pBADfw | **WT** | 90073533 | RMphoQ2_035 | pBADfw | **F** |
| 90073486 | RMphoQ036 | pBADfw | **WT** | 90073534 | RMphoQ2_036 | pBADfw | **L247Q mixed with WT** |
| 90073487 | RMphoQ037 | pBADfw | **WT** | 90073535 | RMphoQ2_037 | pBADfw | **WT** |
| 90073488 | RMphoQ038 | pBADfw | **V198E, E230L** | 90073536 | RMphoQ2_038 | pBADfw | **WT** |
| 90073489 | RMphoQ039 | pBADfw | **WT** | 90073537 | RMphoQ2_039 | pBADfw | **WT** |
| 90073490 | RMphoQ040 | pBADfw | **WT** | 90073538 | RMphoQ2_040 | pBADfw | **WT** |
| 90073491 | RMphoQ041 | pBADfw | **WT** | 90073539 | RMphoQ2_041 | pBADfw | **WT** |
| 90073492 | RMphoQ042 | pBADfw | **V198A, R219C** | 90073540 | RMphoQ2_042 | pBADfw | **FS** |

| Label Numbe | Sample Name | Primer Name | sequencing results |
|---|---|---|---|
| 90073546 | RMphoQ3_001 | pBADfw | **F** |
| 90073547 | RMphoQ3_002 | pBADfw | **I221T** |
| 90073548 | RMphoQ3_003 | pBADfw | **NS** |
| 90073549 | RMphoQ3_004 | pBADfw | **R262C** |
| 90073550 | RMphoQ3_005 | pBADfw | **WT** |
| 90073551 | RMphoQ3_006 | pBADfw | **V251I** |
| 90073552 | RMphoQ3_007 | pBADfw | **FS** |
| 90073553 | RMphoQ3_008 | pBADfw | **S217C** |
| 90073554 | RMphoQ3_009 | pBADfw | **WT** |
| 90073555 | RMphoQ3_010 | pBADfw | **A223T** |
| 90073556 | RMphoQ3_011 | pBADfw | **D274N** |
| 90073557 | RMphoQ3_012 | pBADfw | **FS** |
| 90073558 | RMphoQ3_013 | pBADfw | **FS** |
| 90073559 | RMphoQ3_014 | pBADfw | **WT** |
| 90073560 | RMphoQ3_015 | pBADfw | **FS** |
| 90073561 | RMphoQ3_016 | pBADfw | **W211R** |
| 90073562 | RMphoQ3_017 | pBADfw | **L210M** |
| 90073563 | RMphoQ3_018 | pBADfw | **WT** |
| 90073564 | RMphoQ3_019 | pBADfw | **FS** |
| 90073565 | RMphoQ3_020 | pBADfw | **D274N** |
| 90073566 | RMphoQ3_021 | pBADfw | **S217C** |
| 90073567 | RMphoQ3_022 | pBADfw | **V206N** |
| 90073568 | RMphoQ3_023 | pBADfw | **T270R** |
| 90073569 | RMphoQ3_024 | pBADfw | **L203Q** |
| 90073570 | RMphoQ3_025 | pBADfw | **FS** |
| 90073571 | RMphoQ3_026 | pBADfw | **F** |
| 90073572 | RMphoQ3_027 | pBADfw | **L209P** |

| Label Numbe | Sample Name | Primer Name | sequencing results |
|---|---|---|---|
| 90073573 | RMphoQ3_028 | pBADfw | **WT** |
| 90073574 | RMphoQ3_029 | pBADfw | **WT** |
| 90073575 | RMphoQ3_030 | pBADfw | **NS** |
| 90073576 | RMphoQ3_031 | pBADfw | **R256L** |
| 90073577 | RMphoQ3_032 | pBADfw | **W216R, E230D** |
| 90073578 | RMphoQ3_033 | pBADfw | **IS** |
| 90073579 | RMphoQ3_034 | pBADfw | **WT** |
| 90073580 | RMphoQ3_035 | pBADfw | **F** |
| 90073581 | RMphoQ3_036 | pBADfw | **F** |
| 90073582 | RMphoQ3_037 | pBADfw | **IS** |
| 90073583 | RMphoQ3_038 | pBADfw | **A201S** |

| Label Numbe | Sample Name | Primer Name | sequencing results |
|---|---|---|---|
| 90073594 | RMphoQ4_001 | pBADfw | **FS** |
| 90073595 | RMphoQ4_002 | pBADfw | **A214T** |
| 90073596 | RMphoQ4_003 | pBADfw | **FS** |
| 90073597 | RMphoQ4_004 | pBADfw | **FS** |
| 90073598 | RMphoQ4_005 | pBADfw | **L254R** |
| 90073599 | RMphoQ4_006 | pBADfw | **FS** |
| 90073600 | RMphoQ4_007 | pBADfw | **F195L** |
| 90073601 | RMphoQ4_008 | pBADfw | **FS** |
| 90073602 | RMphoQ4_009 | pBADfw | **IS** |
| 90073603 | RMphoQ4_010 | pBADfw | **WT** |
| 90073604 | RMphoQ4_011 | pBADfw | **FS** |
| 90073605 | RMphoQ4_012 | pBADfw | **IS** |
| 90073606 | RMphoQ4_013 | pBADfw | **F195L** |
