## Supplementary material for "Functional determinants of a small protein controlling a broadly conserved bacterial sensor kinase": MgrB alignment

CLUSTAL O(1.2.4) multiple sequence alignment

```
ppe:Proteus                 ----------------MNAKKIIISLIIALVITFGLYLVALDNFCDRGE-DFQQGLCRFT     43
prq:CYG50_03805             -----------MKELSVFGRKLI-IAVSAVLACIFFYLLAMDNLCDQGGENFIFGICRVT     48
psi:S70_01320               -----------------MKYKLI-IGVCIVITCIFLYLLAMDSLCDQGGENFIFGICQIT     42
psb:Providencia             -----------------MKYKLI-IGVCIVITCIFLYLLAMDSLCDQGGENFIFGICQIT     42
pvc:G3341_09230             -----------------MKYKLV-IGICIVITCIFLYLLAMDSLCDQGGENFIYGICGVT     42
sod:Sant_1773               -------MIRHFRNRVLKRGTTI-KLVAILLLCLALWLSALNSFCDQGG-DFFSGMCLVT     51
pes:SOPEG_1711              ----------------MKRGTTI-KLVAILLLCLALWLSALNSFCDQGG-DFFSGMCLVT     42
raa:Q7S_14675               -----------MKKPVFNVRRII-TFVLFLLICIGLYLLALNSYCDQGG-NFDLGVCAIT     47
rox:BV494_09480             ------------------------MFVLFLLICIGLYLLALNSYCDQGG-NFDLGVCSIT     35
plu:plu2786                 ----------------MDIKKTI-AILALLGICLFLYLLALDSYCDQGE-QFDSGICSVT     42
pay:PAU_01751               -------MLTKLKEIELDIKKTI-AILVLLGICLFLYLLALDRYCDQGE-KFDSGICFVT     51
yrb:UGYR_03160              MLGQNRVDENILKVPDVKIKKLV-ATVFLIAVCCLFYLLALDSYCDQGG-SFSNGICSIT     58
yma:DA391_09980             ----------------MNIKKLV-ATVGIIAVCCLFYLLALDSYCDQGG-TFSTGLCTIT     42
yey:Y11_06121               ----------------MNIKKLV-ATVGIIAVCCLFYLLALDSYCDQGG-TFSTGMCTIT     42
yca:F0T03_09435             ----------------MNIKKLV-AAVAIIAVCCLFYLLALDSYCDQGG-TFSAGICTIT     42
yhi:D5F51_06855             ----------------MNIKKLV-AAVVIIAVCCLFYLLALDSYCDQGG-TFSMGICTIT     42
yel:LC20_03176              ----------------MNIKKLV-AAVVIIAVCCLFYLLALDSYCDQGG-TFSMGICTIT     42
ypk6:Yersinia              -------------MLDLNITKLV-TTVVIIAACCLFYLLALDSYCDQGG-TFSTGICAIT     45
sfw:WN53_23485              ----------------MGRKKLL-VLAATLVACLLFYLLALDSYCDQGG-KFALGICSIT     42
sfg:AV650_19225             ----------------MGRKKLL-VIAATLFACLLFYLLALDSYCDQGE-KFALGICSIT     42
serm:CLM71_13705            ----------------MLRNKIL-LIVATVLACLFFNLLALDSYCDQGG-NFALGICSIT     42
smaf:D781_2649              -----------MRLLSLLRNKIL-LIAATVLACLFFNLLALDSYCDQGG-NFALGICSIT     47
smac:SMDB11_2102            -----------MRLLNLLRKKIL-VIMTAVAACLFFYLLALDSYCDDGG-NFALGICSVT     47
smar:SM39_2308              -----------MRLLNLLRKKIL-VIMTAVAACLFFYLLALDSYCDDGG-NFALGICSVT     47
sers:SERRSCBI_13410         --------------------------MTAVAACLFFYLLALDSYCDDGG-NFALGICSVT     33
serf:L085_14565             --------------------------MTAVAACLFFYLLALDSYCDDGG-NFALGICSVT     33
gqu:AWC35_22335             ------------------------MIIAILAACLFFYLLALDSYCDEDG-SFALGICSIT     35
squ:E4343_01905             ----------------MLRKKIL-VIIAFLAACLFFYLLALDSYCDAGG-TFALGVCSVT     42
spe:Spro_2829               ------------------------MIIAFLAACLFFYLLALDSYCDAGG-TFALGVCSVT     35
yre:HEC60_04460             ----------------MKKFRWVILVVVIILVCMVLWAQMINVMCDQGV-PFFNGICTIN     43
ahn:NCTC12129_02021         ----------------MKKLKWALLVVA-LLLCLLLWAQTLNAMCDQDV-PFFSGICIIN     42
kle:AO703_12345             ----------------MKKLRWIIVIVVLLIACLLLWTQTINVLCDQDV-QFFSGICTIN     43
esc:Entcl_1967              ----------------MKKLRWIIVIVVLLIACLLLWTQTINVLCDQDV-QFFSGICTIN     43
pge:LG71_14635              ----------------MKKLPRIILVVVLLIACLLLWTQMINVMCDQDV-QFFSGICTVN     43
ebf:D782_1816               ----------------MKKFRWVLLIVV-LAACLLLWTQTLNVLCDQDV-QFFNGICSIN     42
kgo:CEW81_10130             ----------------MKKFRWVLLIIV-IAACLLLWTQMINVMCDQDV-QFFSGVCSIN     42
rao:DSD31_09510             ----------------MKKVRWVLLIVI-IAGCLLLWTQMLNVMCDQDV-QFFSGICTIN     42
rpln:B1209_09510            ----------------MKKVRWVLLIVI-IAGCLLLWTQMLNVMCDQDV-QFFSGICTIN     42
ror:RORB6_03065             ----------------MKKVRWVLLIVI-IAGCLLLWTQMLNVMCDQDV-QFFSGICTIN     42
ebu:CUC76_24835             ----------------MKKLRWVLLIVI-IAGCLLLWTQMLNVMCDQDV-QFFSGICTIN     42
klw:DA718_11175             ----------------MKKLRWVLLIVI-IAGCLLLWTQMLNVMCDQDV-QFFSGICTIN     42
```

```
kll:BJF97_17875          ----------------MKKLRWVLLIVI-IAGCLLLWTQMLNVMCDQDV-QFFSGICTIN     42
kqv:B8P98_10630          ----------------MKKLRWVLLIVI-IAGCLLLWTQMLNVMCDQDV-QFFSGICTIN     42
koc:AB185_18145          ----------------MKKLRWVLLIVI-IAGCLLLWTQMLNVMCDQDV-QFFSGICTIN     42
kok:KONIH1_17980         ----------------MKKLRWVLLIVI-IAGCLLLWTQMLNVMCDQDV-QFFSGICTIN     42
kom:HR38_22005           ----------------MKKLRWVLLIVI-IAGCLLLWTQMLNVMCDQDV-QFFSGICTIN     42
koy:J415_13935           ----------------MKKLRWVLLIVI-IAGCLLLWTQMLNVMCDQDV-QFFSGICTIN     42
koe:A225_3631            ----------------MKKLRWVLLIVI-IAGCLLLWTQMLNVMCDQDV-QFFSGICTIN     42
kox:KOX_23700            ----------------MKKLRWVLLIVI-IAGCLLLWTQMLNVMCDQDV-QFFSGICTIN     42
kvq:SP68_05400           ----------------MKKLRWVLLIVI-IAGCLLLWTQMLNVMCDQDV-QFFSGICTIN     42
kvd:KR75_25030           ----------------MKKLRWVLLIVI-IAGCLLLWTQMLNVMCDQDV-QFFSGICTIN     42
kpk:A593_01320           ----------------MKKLRWVLLIVI-IAGCLLLWTQMLNVMCDQDV-QFFSGICTIN     42
kva:Kvar_1840            ----------------MKKLRWVLLIVI-IAGCLLLWTQMLNVMCDQDV-QFFSGICTIN     42
kpi:D364_11900           ----------------MKKLRWVLLIVI-IAGCLLLWTQMLNVMCDQDV-QFFSGICTIN     42
kpo:KPN2242_14680        ----------------MKKLRWVLLIVI-IAGCLLLWTQMLNVMCDQDV-QFFSGICTIN     42
kpe:KPK_1949             ----------------MKKLRWVLLIVI-IAGCLLLWTQMLNVMCDQDV-QFFSGICTIN     42
kpt:VK055_0108           ----------------MKKLRWVLLIVI-IAGCLLLWTQMLNVMCDQDV-QFFSGICTIN     42
kpp:A79E_1892            ----------------MKKLRWVLLIVI-IAGCLLLWTQMLNVMCDQDV-QFFSGICTIN     42
kpu:KP1_3468             ----------------MKKLRWVLLIVI-IAGCLLLWTQMLNVMCDQDV-QFFSGICTIN     42
ear:CCG29473             ----------------MKKLRWVLLIVV-IAGCLLLWTQMLNVMCDQDV-QFFSGICAIN     42
eae:EAE_22650            ----------------MKKLRWVLLIVV-IAGCLLLWTQMLNVMCDQDV-QFFSGICAIN     42
ron:TE10_18220           ----------------MKKLRWVLLIVV-IAGCLLLWTQMLNVMCDQDV-QFFSGICTIN     42
izh:FEM41_19715          ----------------MRKYWWIILIVV-LIACLLMWTQMLNAMCDQDE-PFFNGICVVN     42
clap:NCTC11466_01805     ----------------MRKYRWVIVFII-VVTCLLLWTQMINVMCDQDV-QFFSGVCTIN     42
lea:GNG26_13585          ----------------MKKIRWVILVLV-LVACVLLWTQTINVMCDQDV-QFFSGICSIN     42
ler:GNG29_13315          ----------------MKKIRWVILVLV-LVACVLLWTQTINVMCDQDV-QFFSGICSIN     42
lee:DVA44_09365          ----------------MKKIRWVILVLV-LVACVLLWTQTINVMCDQDV-QFFSGICSIN     42
lei:C2U54_18790          ----------------MKKIRWVILVLV-LVACVLLWTQTINVMCDQDV-QFFSGICSIN     42
leh:C3F35_20965          ----------------MKKIRWVILVLV-LLACVLLWTQTINVMCDQDV-QFFSGICSVN     42
lax:APT61_08755          ----------------MKKIRWVILVLV-LLACVLLWTQTINVMCDQDV-QFFSGICSVN     42
ent:Ent638_2395          ----------------MKKIRWVILIIV-LIACVILWTQTINVMCDQDV-QFFSGICAIN     42
lew:DAI21_13880          ----------------MKKFRWVILVVV-LIACVILWTQTINVMCDQDV-QFFSGICAIN     42
lef:LJPFL01_2460         ----------------MKKFRWVILIVV-LIACVILWTQTINVMCDQDV-QFFSGICAIN     42
laz:A8A57_12460          ----------------MKKIRWVILIIV-LVACAFLWTQTINVMCDQDV-QFFSGICAIN     42
eclc:ECR091_12750        ----------------MKKIRWVILIIV-LIACVVLWTQTINVMCDQDV-QFFSGICSIN     42
ecla:ECNIH3_12815        ----------------MKKIRWVILIIV-LIACVVLWTQTINVMCDQDV-QFFSGICSIN     42
exf:BFV63_13130          ----------------MKKIRWVILIIV-LIACVVLWTQTINVMCDQDV-QFFSGICSIN     42
ehm:AB284_12310          ----------------MKKIRWVILIIV-LIACVVLWTQTINVMCDQDV-QFFSGICSIN     42
eclz:LI64_12870          ----------------MKKIRWVILIIV-LIACVVLWTQTINVMCDQDV-QFFSGICSIN     42
ecly:LI62_14820          ----------------MKKIRWVILIIV-LIACVVLWTQTINVMCDQDV-QFFSGICSIN     42
eclx:LI66_13130          ----------------MKKIRWVILIIV-LIACVVLWTQTINVMCDQDV-QFFSGICSIN     42
ecli:ECNIH5_12770        ----------------MKKIRWVILIIV-LIACVVLWTQTINVMCDQDV-QFFSGICSIN     42
ecle:ECNIH2_13305        ----------------MKKIRWVILIIV-LIACVVLWTQTINVMCDQDV-QFFSGICSIN     42
elg:BH714_09510          ------------------------MIIV-LVACVVLWTQTINVMCDQDV-QFFSGVCAIN     34
eec:EcWSU1_02751         ----------------MKKIRWVILIIV-LVACVVLWTQTINVMCDQDV-QFFSGVCAIN     42
```

```
eas:Entas_2535           ----------------MKKIRWVILVIV-LIACVVLWTQTINVMCDQDV-QFFSGVCAIN     42
eclg:EC036_26650         ----------------MKKIRWVILVIV-LIACVVLWTQTINVMCDQDV-QFFSGVCAIN     42
enl:A3UG_13770           ----------------MKKIRWVILVIV-LIACVVLWTQTINVMCDQDV-QFFSGVCAIN     42
enc:ECL_01476            ----------------MKKIRWVILVIV-LIACVVLWTQTINVMCDQDV-QFFSGVCAIN     42
end:A4308_17820          ----------------MKKIRWVILIVV-LIACVVLWTQTINVMCDQDV-QFFSGVCAIN     42
ekb:BFV64_13565          ----------------MKKIRWVILIVV-LIACVVLWTQTINVMCDQDV-QFFSGVCAIN     42
ecln:ECNIH4_09440        ----------------MKKIRWVILIVV-LIACVVLWTQTINVMCDQDV-QFFSGVCAIN     42
eno:ECENHK_13510         ----------------MKKIRWVILIVV-LIACVVLWTQTINVMCDQDV-QFFSGVCAIN     42
lni:CWR52_21655          ----------------MKKIRWVILIIV-LIACVVLWTQTINVMCDQDV-QFFSGVCAIN     42
ebg:FAI37_15810          ----------------MKKIRWVILIIV-LIACVVLWTQTINVMCDQDV-QFFSGVCAIN     42
enx:NI40_013485          ----------------MKKIRWVILIIV-LIACVVLWTQTINVMCDQDV-QFFSGVCAIN     42
esh:C1N69_13625          ----------------MKKIRWVILIIV-LIACVVLWTQTINVMCDQDV-QFFSGVCAIN     42
ern:BFV67_13165          ----------------MKKIRWVILIIV-LIACVVLWTQTINVMCDQDV-QFFSGVCAIN     42
ecan:CWI88_09260         ----------------MKKIRWVILIIV-LIACVVLWTQTINVMCDQDV-QFFSGVCAIN     42
eau:DI57_05700           ----------------MKKIRWVILIIV-LIACVVLWTQTINVMCDQDV-QFFSGVCAIN     42
buf:D8682_04555          ----------------MKKYRWFILVAV-FFVCLLLWTQMVNVMCDQDV-QFFSGICTIN     42
krd:A3780_13585          ----------------MRKYKWLILIVV-LVGCLLLWTQMLNVMCDQDV-QFFSGICIVN     42
kor:AWR26_10555          ----------------MRKYKWLILIVV-LVGCLLLWTQMLNVMCDQDV-QFFSGICIVN     42
ccon:AFK62_11425         ----------------MKKLRWAILLAV-LVACLLLWAQMINVMCDQDV-QFFSGICTIN     42
csj:CSK29544_02729       ----------------MKKFRWAILLAV-LVACLLLWMQTLNVMCDQDV-QFFSGICTIN     42
csz:CSSP291_06955        ----------------MKKFRWAILLAV-LVACLLLWMQTLNVMCDQDV-QFFSGICTIN     42
esa:ESA_01425            ----------------MKKFRWAILLAV-LVACLLLWMQTLNVMCDQDV-QFFSGICTIN     42
ctu:CTU_25090            ----------------MKKFRWAILLAV-LVACLLLWTQTINVMCDQDV-QFFSGICTIN     42
cui:AFK65_11815          ----------------MKKFRWAILLAV-LVACLLLWTQTINVMCDQDV-QFFSGICTIN     42
cmw:AFK63_06815          ----------------MKKLRWAILLAV-LVACVLLWTQTINVMCDQDV-QFFSGICTIN     42
cmj:AFK66_007820         ----------------MKKLRWAILLAV-LVACLLLWTQTINVMCDQDV-QFFSGICTIN     42
csi:P262_02367           ----------------MKKLRWAILLAV-LVACLLLWTQTINVMCDQDV-QFFSGICTIN     42
cdm:AFK67_11770          ----------------MKKLRWAILLAV-LVACLLLWTQTINVMCDQDV-QFFSGICTIN     42
kot:EH164_09600          ----------------MRKFRWLLLIAV-VVVCLLLWTQMINVMCDQDV-QFFSGICVIN     42
kco:BWI95_18645          ----------------MRKFRWLLLIAV-VVVCLLLWTQMINVMCDQDV-QFFSGICVIN     42
enf:AKI40_3542           ----------------MRKFRWLILAAV-LVVCLLLWTQMINVMCDQDV-QFFSGICVIN     42
cwe:CO701_08325          ----------------MKKIRWVVLIV-VILVCILMWAQVFNIMCDQDV-QFFNGICAIN     42
cfq:C2U38_10190          ----------------MKKIRWVVLIV-VVLVCILMWAQVFNIMCDQDV-QFFSGICAFN     42
cpar:CUC49_09605         ----------------MKKIRWVVLIV-VVLVCILMWAQVFNIMCDQDV-QFFSGICAIN     42
cif:AL515_13855          ----------------MKKIRWVVLIV-VVLVCILMWAQVFNIMCDQDV-QFFSGICAIN     42
cpot:FOB25_21665         ----------------MKKIRWVVLIV-VVLVCILMWAQVFNIMCDQDV-QFFSGICAIN     42
cyo:CD187_10070          ----------------MKKIRWVVLIV-VVLVCILMWAQVFNIMCDQDV-QFFSGICAIN     42
cbra:A6J81_06490         ----------------MKKIRWVVLIV-VVLVCILMWAQVFNIMCDQDV-QFFSGICAIN     42
cie:AN232_16140          ----------------MKKIRWVVLIV-IVLVCILMWAQVFNIMCDQDV-QFFSGICAIN     42
cfd:CFNIH1_19945         ----------------MKKIRWVVLIV-IVLVCILMWAQVFNIMCDQDV-QFFSGICAIN     42
efe:EFER_1249            ------------MEHEVKKFRWLVLIV-VVLACLVLWAQVINIMCDQDV-QFFSGICAIN     46
cro:ROD_18651            ----------------MKKFRWIILVMVVVPVCLLLWAQVVNLMCDQDV-QFFSGICAIN     43
cko:CKO_01149            ----------MDAESGVKKFRWVILII-VALVCLLLWAQVFNIMCDQDV-QFFNGICAIN     48
cir:C2U53_24200          ----------------MKKFRWIILVV-VVLVCLLLWAQVFNIMCDQDV-QFFSGICAIN     42
```

```
cfar:CI104_15185          ----------------MKKFRWIILVV-VVLVCLLLWAQVFNIMCDQDV-QFFSGICAIN     42
caf:AL524_13460           ----------------MKKFRWIILVV-VVLVCLLLWAQVFNIMCDQDV-QFFSGICAIN     42
cama:F384_08800           ----------------MKKFRWIILVM-VVLVCLLLWAQVFNIMCDQDV-QFFSGICAIN     42
sed:SeD_A1475             ----------------MKTFRWVVLGI-VVVVCLLLWAQVFNIMCDQDV-QFFSGICAIN     42
salz:EOS98_10020          ----------------MKKFRWVVLGI-VVVVCLLLWAQVFNIMCDQDV-QFFSGICAIN     42
sbv:N643_08235            ----------------MKKFRWVVLGI-VVVVCLLLWAQVFNIMCDQDV-QFFSGICAIN     42
sbz:A464_1940             ----------------MKKFRWVVLGI-VVVVCLLLWAQVFNIMCDQDV-QFFSGICAIN     42
ses:SARI_01101            ----------------MKKFRWVVLGI-VVVVCLLLWAQVFNIMCDQDV-QFFSGICAIN     42
senc:SEET0819_15940       ----------------MKKFRWVVLGI-VVVVCLLLWAQVFNIMCDQDV-QFFSGICAIN     42
sene:IA1_09130            ----------------MKKFRWVVLGI-VVVVCLLLWAQVFNIMCDQDV-QFFSGICAIN     42
senb:BN855_18970          ----------------MKKFRWVVLGI-VVVVCLLLWAQVFNIMCDQDV-QFFSGICAIN     42
seeb:SEEB0189_010300      ----------------MKKFRWVVLGI-VVVVCLLLWAQVFNIMCDQDV-QFFSGICAIN     42
seec:CFSAN002050_15670    ----------------MKKFRWVVLGI-VVVVCLLLWAQVFNIMCDQDV-QFFSGICAIN     42
senj:CFSAN001992_02275    ----------------MKKFRWVVLGI-VVVVCLLLWAQVFNIMCDQDV-QFFSGICAIN     42
senl:IY59_06320           ----------------MKKFRWVVLGI-VVVVCLLLWAQVFNIMCDQDV-QFFSGICAIN     42
senq:AU40_06935           ----------------MKKFRWVVLGI-VVVVCLLLWAQVFNIMCDQDV-QFFSGICAIN     42
senv:AU39_06160           ----------------MKKFRWVVLGI-VVVVCLLLWAQVFNIMCDQDV-QFFSGICAIN     42
seno:AU37_06160           ----------------MKKFRWVVLGI-VVVVCLLLWAQVFNIMCDQDV-QFFSGICAIN     42
sena:AU38_06165           ----------------MKKFRWVVLGI-VVVVCLLLWAQVFNIMCDQDV-QFFSGICAIN     42
set:SEN1197               ----------------MKKFRWVVLGI-VVVVCLLLWAQVFNIMCDQDV-QFFSGICAIN     42
sens:Q786_05995           ----------------MKKFRWVVLGI-VVVVCLLLWAQVFNIMCDQDV-QFFSGICAIN     42
sea:SeAg_B1291            ----------------MKKFRWVVLGI-VVVVCLLLWAQVFNIMCDQDV-QFFSGICAIN     42
sew:SeSA_A1983            ----------------MKKFRWVVLGI-VVVVCLLLWAQVFNIMCDQDV-QFFSGICAIN     42
senn:SN31241_29250        ----------------MKKFRWVVLGI-VVVVCLLLWAQVFNIMCDQDV-QFFSGICAIN     42
see:SNSL254_A1979         ----------------MKKFRWVVLGI-VVVVCLLLWAQVFNIMCDQDV-QFFSGICAIN     42
seeh:SEEH1578_18455       ----------------MKKFRWVVLGI-VVVVCLLLWAQVFNIMCDQDV-QFFSGICAIN     42
senh:CFSAN002069_22645    ----------------MKKFRWVVLGI-VVVVCLLLWAQVFNIMCDQDV-QFFSGICAIN     42
shb:SU5_02441             ----------------MKKFRWVVLGI-VVVVCLLLWAQVFNIMCDQDV-QFFSGICAIN     42
seh:SeHA_C2041            ----------------MKKFRWVVLGI-VVVVCLLLWAQVFNIMCDQDV-QFFSGICAIN     42
sec:SCH_1836              ----------------MKKFRWVVLGI-VVVVCLLLWAQVFNIMCDQDV-QFFSGICAIN     42
sei:SPC_1889              ----------------MKKFRWVVLGI-VVVVCLLLWAQVFNIMCDQDV-QFFSGICAIN     42
spq:SPAB_01371            ----------------MKKFRWVVLGI-VVVVCLLLWAQVFNIMCDQDV-QFFSGICAIN     42
sek:SSPA0963              ----------------MKKFRWVVLGI-VVVVCLLLWAQVFNIMCDQDV-QFFSGICAIN     42
spt:SPA1033               ----------------MKKFRWVVLGI-VVVVCLLLWAQVFNIMCDQDV-QFFSGICAIN     42
seen:SE451236_15135       ----------------MKKFRWVVLGI-VVVVCLLLWAQVFNIMCDQDV-QFFSGICAIN     42
seni:CY43_09385           ----------------MKKFRWVVLGI-VVVVCLLLWAQVFNIMCDQDV-QFFSGICAIN     42
senr:STMDT2_17601         ----------------MKKFRWVVLGI-VVVVCLLLWAQVFNIMCDQDV-QFFSGICAIN     42
setc:CFSAN001921_07895    ----------------MKKFRWVVLGI-VVVVCLLLWAQVFNIMCDQDV-QFFSGICAIN     42
setu:STU288_05560         ----------------MKKFRWVVLGI-VVVVCLLLWAQVFNIMCDQDV-QFFSGICAIN     42
sef:UMN798_1936           ----------------MKKFRWVVLGI-VVVVCLLLWAQVFNIMCDQDV-QFFSGICAIN     42
seb:STM474_1863           ----------------MKKFRWVVLGI-VVVVCLLLWAQVFNIMCDQDV-QFFSGICAIN     42
sej:STMUK_1813            ----------------MKKFRWVVLGI-VVVVCLLLWAQVFNIMCDQDV-QFFSGICAIN     42
sem:STMDT12_C18600        ----------------MKKFRWVVLGI-VVVVCLLLWAQVFNIMCDQDV-QFFSGICAIN     42
sey:SL1344_1769           ----------------MKKFRWVVLGI-VVVVCLLLWAQVFNIMCDQDV-QFFSGICAIN     42
```

```
seo:STM14_2226             ----------------MKKFRWVVLGI-VVVVCLLLWAQVFNIMCDQDV-QFFSGICAIN     42
stm:STM1840                ----------------MKKFRWVVLGI-VVVVCLLLWAQVFNIMCDQDV-QFFSGICAIN     42
sent:TY21A_05295           ----------------MKKFRWVVLGI-VVVVCLLLWAQVFNIMCDQDV-QFFSGICAIN     42
sex:STBHUCCB_11130         ----------------MKKFRWVVLGI-VVVVCLLLWAQVFNIMCDQDV-QFFSGICAIN     42
stt:t1038                  ----------------MKKFRWVVLGI-VVVVCLLLWAQVFNIMCDQDV-QFFSGICAIN     42
sty:STY1970                ----------------MKKFRWVVLGI-VVVVCLLLWAQVFNIMCDQDV-QFFSGICAIN     42
seep:I137_07490            ----------------MKKFRWVVLGI-VVVVCLLLWAQVFNIMCNQDV-QFFSGICAIN     42
sega:SPUCDC_1657           ----------------MKKFRWVVLGI-VVVVCLLLWAQVFNIMCNQDV-QFFSGICAIN     42
sel:SPUL_1657              ----------------MKKFRWVVLGI-VVVVCLLLWAQVFNIMCNQDV-QFFSGICAIN     42
seg:SG1277                 ----------------MKKFRWVVLGI-VVVVCLLLWAQVFNIMCNQDV-QFFSGICAIN     42
esz:FEM44_24025            ----------------MKKFRWVVLVA-AMLAFLLLWMQVFNIMCDQDV-QFFSGICAIN     42
elp:P12B_c1257             ----------------------------MVLACLLLWAQVFNMMCDQDV-QFFSGICAIN     31
elu:UM146_08040            ----------------MKKFRWVALVV-VVLACLLLWAQVFNMMCDQDV-QFFSGICALN     42
ecz:ECS88_1878             ----------------MKKFRWVALVV-VVLACLLLWAQVFNMMCDQDV-QFFSGICALN     42
eih:ECOK1_1942             ----------------MKKFRWVALVV-VVLACLLLWAQVFNMMCDQDV-QFFSGICALN     42
eci:UTI89_C2026            ----------------MKKFRWVALVV-VVLACLLLWAQVFNMMCDQDV-QFFSGICALN     42
eoi:ECO111_2333            ----------------MKKFRWVVLVV-VVLACLLLWAQVFNMMCDQDV-QFFSGICAIN     42
elr:ECO55CA74_10945        ----------------MKKFRWVVLVV-VVLACLLLWAQVFNMMCDQDV-QFFSGICAIN     42
eok:G2583_2275             ----------------MKKFRWVVLVV-VVLACLLLWAQVFNMMCDQDV-QFFSGICAIN     42
ecoj:P423_09680            ----------------MKKFRWVALVV-VVLACLLLWAQVFNMMCDQDV-QFFSGICAIN     42
elf:LF82_1341              ----------------MKKFRWVALVV-VVLACLLLWAQVFNMMCDQDV-QFFSGICAIN     42
eld:i02_2052               ----------------MKKFRWVALVV-VVLACLLLWAQVFNMMCDQDV-QFFSGICAIN     42
elc:i14_2052               ----------------MKKFRWVALVV-VVLACLLLWAQVFNMMCDQDV-QFFSGICAIN     42
ese:ECSF_1682              ----------------MKKFRWVALVV-VVLACLLLWAQVFNMMCDQDV-QFFSGICAIN     42
eln:NRG857_09130           ----------------MKKFRWVALVV-VVLACLLLWAQVFNMMCDQDV-QFFSGICAIN     42
ecc:c2234                  ----------------MKKFRWVALVV-VVLACLLLWAQVFNMMCDQDV-QFFSGICAIN     42
ecq:ECED1_2029             ----------------MKKFRWVALVV-VVLACLLLWAQVFNMMCDQDV-QFFSGICAIN     42
ecos:EC958_2044            ----------------MKKFRWVALVV-VVLACLLLWAQVFNMMCDQDV-QFFSGICAIN     42
ena:ECNA114_1871           ----------------MKKFRWVALVV-VVLACLLLWAQVFNMMCDQDV-QFFSGICAIN     42
ecg:E2348C_1950            ----------------MKKFRWVALVV-VVLACLLLWAQVFNMMCDQDV-QFFSGICAIN     42
shq:A0259_13235            ----------------MKKFRWVVLVV-VVLACLLLWAQVFNMMCDQDV-QFFSGICAIN     42
sdz:Asd1617_02658          ----------------MKKFRWVVLVV-VVLACLLLWAQVFNMMCDQDV-QFFSGICAIN     42
sdy:SDY_1973               ----------------MKKFRWVVLVV-VVLACLLLWAQVFNMMCDQDV-QFFSGICAIN     42
sbc:SbBS512_E2093          ----------------MKKFRWVVLVV-VVLACLLLWAQVFNMMCDQDV-QFFSGICAIN     42
sbo:SBO_1239               ----------------MKKFRWVVLVV-VVLACLLLWAQVFNMMCDQDV-QFFSGICAIN     42
ssn:SSON_1335              ----------------MKKFRWVVLVV-VVLACLLLWAQVFNMMCDQDV-QFFSGICAIN     42
sfs:SFyv_2061              ----------------MKKFRWVVLVV-VVLACLLLWAQVFNMMCDQDV-QFFSGICAIN     42
sfn:SFy_2009               ----------------MKKFRWVVLVV-VVLACLLLWAQVFNMMCDQDV-QFFSGICAIN     42
sfe:SFxv_1587              ----------------MKKFRWVVLVV-VVLACLLLWAQVFNMMCDQDV-QFFSGICAIN     42
sfv:SFV_1403               ----------------MKKFRWVVLVV-VVLACLLLWAQVFNMMCDQDV-QFFSGICAIN     42
sfx:S1515                  ----------------MKKFRWVVLVV-VVLACLLLWAQVFNMMCDQDV-QFFSGICAIN     42
sfl:SF1400                 ----------------MKKFRWVVLVV-VVLACLLLWAQVFNMMCDQDV-QFFSGICAIN     42
ecol:LY180_09505           ----------------MKKFRWVVLVV-VVLACLLLWAQVFNMMCDQDV-QFFSGICAIN     42
ell:WFL_09805              ----------------MKKFRWVVLVV-VVLACLLLWAQVFNMMCDQDV-QFFSGICAIN     42
```

```
elw:ECW_m1996               ----------------MKKFRWVVLVV-VVLACLLLWAQVFNMMCDQDV-QFFSGICAIN      42
edj:ECDH1ME8569_1771        ----------------MKKFRWVVLVV-VVLACLLLWAQVFNMMCDQDV-QFFSGICAIN      42
edh:EcDH1_1818              ----------------MKKFRWVVLVV-VVLACLLLWAQVFNMMCDQDV-QFFSGICAIN      42
ekf:KO11_13585              ----------------MKKFRWVVLVV-VVLACLLLWAQVFNMMCDQDV-QFFSGICAIN      42
ecl:EcolC_1807              ----------------MKKFRWVVLVV-VVLACLLLWAQVFNMMCDQDV-QFFSGICAIN      42
ebd:ECBD_1815               ----------------MKKFRWVVLVV-VVLACLLLWAQVFNMMCDQDV-QFFSGICAIN      42
ebe:B21_01784               ----------------MKKFRWVVLVV-VVLACLLLWAQVFNMMCDQDV-QFFSGICAIN      42
ebl:ECD_01796               ----------------MKKFRWVVLVV-VVLACLLLWAQVFNMMCDQDV-QFFSGICAIN      42
ebr:ECB_01796               ----------------MKKFRWVVLVV-VVLACLLLWAQVFNMMCDQDV-QFFSGICAIN      42
eoc:CE10_2109               ----------------MKKFRWVVLVV-VVLACLLLWAQVFNMMCDQDV-QFFSGICAIN      42
eum:ECUMN_2119              ----------------MKKFRWVVLVV-VVLACLLLWAQVFNMMCDQDV-QFFSGICAIN      42
ecr:ECIAI1_1896             ----------------MKKFRWVVLVV-VVLACLLLWAQVFNMMCDQDV-QFFSGICAIN      42
ecy:ECSE_2000               ----------------MKKFRWVVLVV-VVLACLLLWAQVFNMMCDQDV-QFFSGICAIN      42
ecm:EcSMS35_1362            ----------------MKKFRWVVLVV-VVLACLLLWAQVFNMMCDQDV-QFFSGICAIN      42
ecx:EcHS_A1916              ----------------MKKFRWVVLVV-VVLACLLLWAQVFNMMCDQDV-QFFSGICAIN      42
ecoa:APECO78_12930          ----------------MKKFRWVVLVV-VVLACLLLWAQVFNMMCDQDV-QFFSGICAIN      42
eun:UMNK88_2297             ----------------MKKFRWVVLVV-VVLACLLLWAQVFNMMCDQDV-QFFSGICAIN      42
ecw:EcE24377A_2054          ----------------MKKFRWVVLVV-VVLACLLLWAQVFNMMCDQDV-QFFSGICAIN      42
elh:ETEC_1858               ----------------MKKFRWVVLVV-VVLACLLLWAQVFNMMCDQDV-QFFSGICAIN      42
eck:EC55989_2000            ----------------MKKFRWVVLVV-VVLACLLLWAQVFNMMCDQDV-QFFSGICAIN      42
esm:O3M_10790               ----------------MKKFRWVVLVV-VVLACLLLWAQVFNMMCDQDV-QFFSGICAIN      42
eso:O3O_14805               ----------------MKKFRWVVLVV-VVLACLLLWAQVFNMMCDQDV-QFFSGICAIN      42
esl:O3K_10820               ----------------MKKFRWVVLVV-VVLACLLLWAQVFNMMCDQDV-QFFSGICAIN      42
ecoh:ECRM13516_2234         ----------------MKKFRWVVLVV-VVLACLLLWAQVFNMMCDQDV-QFFSGICAIN      42
ecoo:ECRM13514_2331         ----------------MKKFRWVVLVV-VVLACLLLWAQVFNMMCDQDV-QFFSGICAIN      42
eoh:ECO103_2016             ----------------MKKFRWVVLVV-VVLACLLLWAQVFNMMCDQDV-QFFSGICAIN      42
eoj:ECO26_2596              ----------------MKKFRWVVLVV-VVLACLLLWAQVFNMMCDQDV-QFFSGICAIN      42
elx:CDCO157_2370            ----------------MKKFRWVVLVV-VVLACLLLWAQVFNMMCDQDV-QFFSGICAIN      42
etw:ECSP_2399               ----------------MKKFRWVVLVV-VVLACLLLWAQVFNMMCDQDV-QFFSGICAIN      42
ecf:ECH74115_2557           ----------------MKKFRWVVLVV-VVLACLLLWAQVFNMMCDQDV-QFFSGICAIN      42
ecs:ECs2536                 ----------------MKKFRWVVLVV-VVLACLLLWAQVFNMMCDQDV-QFFSGICAIN      42
ece:Z2872                   ----------------MKKFRWVVLVV-VVLACLLLWAQVFNMMCDQDV-QFFSGICAIN      42
ecok:ECMDS42_1500           ----------------MKKFRWVVLVV-VVLACLLLWAQVFNMMCDQDV-QFFSGICAIN      42
ebw:BWG_1639                ----------------MKKFRWVVLVV-VVLACLLLWAQVFNMMCDQDV-QFFSGICAIN      42
ecd:ECDH10B_1964            ----------------MKKFRWVVLVV-VVLACLLLWAQVFNMMCDQDV-QFFSGICAIN      42
ecj:JW1815                  ----------------MKKFRWVVLVV-VVLACLLLWAQVFNMMCDQDV-QFFSGICAIN      42
eco:b1826                   ----------------MKKFRWVVLVV-VVLACLLLWAQVFNMMCDQDV-QFFSGICAIN      42
ema:C1192_03175             ----------------MKKFRWVVLVV-AVLACLLLWAQVFNMMCDQDV-QFFSGICAIN      42
                                                               :    .:  .: .   *  *:* ..

ppe:Proteus                 TLFPSKHH      51
prq:CYG50_03805             DLLPF---      53
psi:S70_01320               DLLPF---      47
psb:Providencia             DLLPF---      47
```

| | | |
|---|---|---|
| pvc:G3341_09230 | DWLPF--- | 47 |
| sod:Sant_1773 | KWMPW--- | 56 |
| pes:SOPEG_1711 | KWMPW--- | 47 |
| raa:Q7S_14675 | SFIPF--- | 52 |
| rox:BV494_09480 | SFIPF--- | 40 |
| plu:plu2786 | RYLPF--- | 47 |
| pay:PAU_01751 | RYLPF--- | 56 |
| yrb:UGYR_03160 | SIVPW--- | 63 |
| yma:DA391_09980 | SIIPW--- | 47 |
| yey:Y11_06121 | SIIPW--- | 47 |
| yca:F0T03_09435 | TIIPW--- | 47 |
| yhi:D5F51_06855 | SIIPW--- | 47 |
| yel:LC20_03176 | SIIPW--- | 47 |
| ypk6:Yersinia | TIVPW--- | 50 |
| sfw:WN53_23485 | RIVPW--- | 47 |
| sfg:AV650_19225 | RIVPW--- | 47 |
| serm:CLM71_13705 | RIIPW--- | 47 |
| smaf:D781_2649 | RIIPW--- | 52 |
| smac:SMDB11_2102 | RFVPW--- | 52 |
| smar:SM39_2308 | RFVPW--- | 52 |
| sers:SERRSCBI_13410 | RFVPW--- | 38 |
| serf:L085_14565 | RFVPW--- | 38 |
| gqu:AWC35_22335 | RFIPW--- | 40 |
| squ:E4343_01905 | RFVPW--- | 47 |
| spe:Spro_2829 | RFVPW--- | 40 |
| yre:HEC60_04460 | KFIPW--- | 48 |
| ahn:NCTC12129_02021 | KYIPW--- | 47 |
| kle:AO703_12345 | KFIPW--- | 48 |
| esc:Entcl_1967 | KFIPW--- | 48 |
| pge:LG71_14635 | KFIPW--- | 48 |
| ebf:D782_1816 | KFIPW--- | 47 |
| kgo:CEW81_10130 | RFIPW--- | 47 |
| rao:DSD31_09510 | KFIPW--- | 47 |
| rpln:B1209_09510 | KFIPW--- | 47 |
| ror:RORB6_03065 | KFIPW--- | 47 |
| ebu:CUC76_24835 | KFIPW--- | 47 |
| klw:DA718_11175 | KFIPW--- | 47 |
| kll:BJF97_17875 | KFIPW--- | 47 |
| kqv:B8P98_10630 | KFIPW--- | 47 |
| koc:AB185_18145 | KFIPW--- | 47 |
| kok:KONIH1_17980 | KFIPW--- | 47 |
| kom:HR38_22005 | KFIPW--- | 47 |
| koy:J415_13935 | KFIPW--- | 47 |
| koe:A225_3631 | KFIPW--- | 47 |

| | | |
|---|---|---|
| kox:KOX_23700 | KFIPW--- | 47 |
| kvq:SP68_05400 | KFIPW--- | 47 |
| kvd:KR75_25030 | KFIPW--- | 47 |
| kpk:A593_01320 | KFIPW--- | 47 |
| kva:Kvar_1840 | KFIPW--- | 47 |
| kpi:D364_11900 | KFIPW--- | 47 |
| kpo:KPN2242_14680 | KFIPW--- | 47 |
| kpe:KPK_1949 | KFIPW--- | 47 |
| kpt:VK055_0108 | KFIPW--- | 47 |
| kpp:A79E_1892 | KFIPW--- | 47 |
| kpu:KP1_3468 | KFIPW--- | 47 |
| ear:CCG29473 | KFIPW--- | 47 |
| eae:EAE_22650 | KFIPW--- | 47 |
| ron:TE10_18220 | KFIPW--- | 47 |
| izh:FEM41_19715 | KFIPW--- | 47 |
| clap:NCTC11466_01805 | KFIPW--- | 47 |
| lea:GNG26_13585 | KFIPW--- | 47 |
| ler:GNG29_13315 | KFIPW--- | 47 |
| lee:DVA44_09365 | KFIPW--- | 47 |
| lei:C2U54_18790 | KFIPW--- | 47 |
| leh:C3F35_20965 | KFIPW--- | 47 |
| lax:APT61_08755 | KFIPW--- | 47 |
| ent:Ent638_2395 | QFIPW--- | 47 |
| lew:DAI21_13880 | QFIPW--- | 47 |
| lef:LJPFL01_2460 | QFIPW--- | 47 |
| laz:A8A57_12460 | KFIPW--- | 47 |
| eclc:ECR091_12750 | KFIPW--- | 47 |
| ecla:ECNIH3_12815 | KFIPW--- | 47 |
| exf:BFV63_13130 | KFIPW--- | 47 |
| ehm:AB284_12310 | KFIPW--- | 47 |
| eclz:LI64_12870 | KFIPW--- | 47 |
| ecly:LI62_14820 | KFIPW--- | 47 |
| eclx:LI66_13130 | KFIPW--- | 47 |
| ecli:ECNIH5_12770 | KFIPW--- | 47 |
| ecle:ECNIH2_13305 | KFIPW--- | 47 |
| elg:BH714_09510 | KFIPW--- | 39 |
| eec:EcWSU1_02751 | KFIPW--- | 47 |
| eas:Entas_2535 | KFIPW--- | 47 |
| eclg:EC036_26650 | KFIPW--- | 47 |
| enl:A3UG_13770 | KFIPW--- | 47 |
| enc:ECL_01476 | KFIPW--- | 47 |
| end:A4308_17820 | KFIPW--- | 47 |
| ekb:BFV64_13565 | KFIPW--- | 47 |
| ecln:ECNIH4_09440 | KFIPW--- | 47 |

| | | |
|---|---|---|
| eno:ECENHK_13510 | KFIPW--- | 47 |
| lni:CWR52_21655 | KFIPW--- | 47 |
| ebg:FAI37_15810 | KFIPW--- | 47 |
| enx:NI40_013485 | KFIPW--- | 47 |
| esh:C1N69_13625 | KFIPW--- | 47 |
| ern:BFV67_13165 | KFIPW--- | 47 |
| ecan:CWI88_09260 | KFIPW--- | 47 |
| eau:DI57_05700 | KFIPW--- | 47 |
| buf:D8682_04555 | KFIPW--- | 47 |
| krd:A3780_13585 | KFIPW--- | 47 |
| kor:AWR26_10555 | KFIPW--- | 47 |
| ccon:AFK62_11425 | KFIPW--- | 47 |
| csj:CSK29544_02729 | KFIPW--- | 47 |
| csz:CSSP291_06955 | KFIPW--- | 47 |
| esa:ESA_01425 | KFIPW--- | 47 |
| ctu:CTU_25090 | KFIPW--- | 47 |
| cui:AFK65_11815 | KFIPW--- | 47 |
| cmw:AFK63_06815 | KFIPW--- | 47 |
| cmj:AFK66_007820 | KFIPW--- | 47 |
| csi:P262_02367 | KFIPW--- | 47 |
| cdm:AFK67_11770 | KFIPW--- | 47 |
| kot:EH164_09600 | KFIPW--- | 47 |
| kco:BWI95_18645 | KFIPW--- | 47 |
| enf:AKI40_3542 | KFIPW--- | 47 |
| cwe:CO701_08325 | KFIPW--- | 47 |
| cfq:C2U38_10190 | KFIPW--- | 47 |
| cpar:CUC49_09605 | KFIPW--- | 47 |
| cif:AL515_13855 | KFIPW--- | 47 |
| cpot:FOB25_21665 | KFIPW--- | 47 |
| cyo:CD187_10070 | KFIPW--- | 47 |
| cbra:A6J81_06490 | KFIPW--- | 47 |
| cie:AN232_16140 | KFIPW--- | 47 |
| cfd:CFNIH1_19945 | KFIPW--- | 47 |
| efe:EFER_1249 | KFIPW--- | 51 |
| cro:ROD_18651 | KFIPW--- | 48 |
| cko:CKO_01149 | KFIPW--- | 53 |
| cir:C2U53_24200 | KFIPW--- | 47 |
| cfar:CI104_15185 | KFIPW--- | 47 |
| caf:AL524_13460 | KFIPW--- | 47 |
| cama:F384_08800 | KFIPW--- | 47 |
| sed:SeD_A1475 | KFIPW--- | 47 |
| salz:EOS98_10020 | KFIPW--- | 47 |
| sbv:N643_08235 | KFIPW--- | 47 |
| sbz:A464_1940 | KFIPW--- | 47 |

| | | |
|---|---|---|
| ses:SARI_01101 | KFIPW--- | 47 |
| senc:SEET0819_15940 | KFIPW--- | 47 |
| sene:IA1_09130 | KFIPW--- | 47 |
| senb:BN855_18970 | KFIPW--- | 47 |
| seeb:SEEB0189_010300 | KFIPW--- | 47 |
| seec:CFSAN002050_15670 | KFIPW--- | 47 |
| senj:CFSAN001992_02275 | KFIPW--- | 47 |
| senl:IY59_06320 | KFIPW--- | 47 |
| senq:AU40_06935 | KFIPW--- | 47 |
| senv:AU39_06160 | KFIPW--- | 47 |
| seno:AU37_06160 | KFIPW--- | 47 |
| sena:AU38_06165 | KFIPW--- | 47 |
| set:SEN1197 | KFIPW--- | 47 |
| sens:Q786_05995 | KFIPW--- | 47 |
| sea:SeAg_B1291 | KFIPW--- | 47 |
| sew:SeSA_A1983 | KFIPW--- | 47 |
| senn:SN31241_29250 | KFIPW--- | 47 |
| see:SNSL254_A1979 | KFIPW--- | 47 |
| seeh:SEEH1578_18455 | KFIPW--- | 47 |
| senh:CFSAN002069_22645 | KFIPW--- | 47 |
| shb:SU5_02441 | KFIPW--- | 47 |
| seh:SeHA_C2041 | KFIPW--- | 47 |
| sec:SCH_1836 | KFIPW--- | 47 |
| sei:SPC_1889 | KFIPW--- | 47 |
| spq:SPAB_01371 | KFIPW--- | 47 |
| sek:SSPA0963 | KFIPW--- | 47 |
| spt:SPA1033 | KFIPW--- | 47 |
| seen:SE451236_15135 | KFIPW--- | 47 |
| seni:CY43_09385 | KFIPW--- | 47 |
| senr:STMDT2_17601 | KFIPW--- | 47 |
| setc:CFSAN001921_07895 | KFIPW--- | 47 |
| setu:STU288_05560 | KFIPW--- | 47 |
| sef:UMN798_1936 | KFIPW--- | 47 |
| seb:STM474_1863 | KFIPW--- | 47 |
| sej:STMUK_1813 | KFIPW--- | 47 |
| sem:STMDT12_C18600 | KFIPW--- | 47 |
| sey:SL1344_1769 | KFIPW--- | 47 |
| seo:STM14_2226 | KFIPW--- | 47 |
| stm:STM1840 | KFIPW--- | 47 |
| sent:TY21A_05295 | KFIPW--- | 47 |
| sex:STBHUCCB_11130 | KFIPW--- | 47 |
| stt:t1038 | KFIPW--- | 47 |
| sty:STY1970 | KFIPW--- | 47 |
| seep:I137_07490 | KFIPW--- | 47 |

| | | |
|---|---|---|
| sega:SPUCDC_1657 | KFIPW--- | 47 |
| sel:SPUL_1657 | KFIPW--- | 47 |
| seg:SG1277 | KFIPW--- | 47 |
| esz:FEM44_24025 | QFIPW--- | 47 |
| elp:P12B_c1257 | QFIPW--- | 36 |
| elu:UM146_08040 | QFIPW--- | 47 |
| ecz:ECS88_1878 | QFIPW--- | 47 |
| eih:ECOK1_1942 | QFIPW--- | 47 |
| eci:UTI89_C2026 | QFIPW--- | 47 |
| eoi:ECO111_2333 | QFNPW--- | 47 |
| elr:ECO55CA74_10945 | QFISW--- | 47 |
| eok:G2583_2275 | QFISW--- | 47 |
| ecoj:P423_09680 | QFIPW--- | 47 |
| elf:LF82_1341 | QFIPW--- | 47 |
| eld:i02_2052 | QFIPW--- | 47 |
| elc:i14_2052 | QFIPW--- | 47 |
| ese:ECSF_1682 | QFIPW--- | 47 |
| eln:NRG857_09130 | QFIPW--- | 47 |
| ecc:c2234 | QFIPW--- | 47 |
| ecq:ECED1_2029 | QFIPW--- | 47 |
| ecos:EC958_2044 | QFIPW--- | 47 |
| ena:ECNA114_1871 | QFIPW--- | 47 |
| ecg:E2348C_1950 | QFIPW--- | 47 |
| shq:A0259_13235 | QFIPW--- | 47 |
| sdz:Asd1617_02658 | QFIPW--- | 47 |
| sdy:SDY_1973 | QFIPW--- | 47 |
| sbc:SbBS512_E2093 | QFIPW--- | 47 |
| sbo:SBO_1239 | QFIPW--- | 47 |
| ssn:SSON_1335 | QFIPW--- | 47 |
| sfs:SFyv_2061 | QFIPW--- | 47 |
| sfn:SFy_2009 | QFIPW--- | 47 |
| sfe:SFxv_1587 | QFIPW--- | 47 |
| sfv:SFV_1403 | QFIPW--- | 47 |
| sfx:S1515 | QFIPW--- | 47 |
| sfl:SF1400 | QFIPW--- | 47 |
| ecol:LY180_09505 | QFIPW--- | 47 |
| ell:WFL_09805 | QFIPW--- | 47 |
| elw:ECW_m1996 | QFIPW--- | 47 |
| edj:ECDH1ME8569_1771 | QFIPW--- | 47 |
| edh:EcDH1_1818 | QFIPW--- | 47 |
| ekf:KO11_13585 | QFIPW--- | 47 |
| ecl:EcolC_1807 | QFIPW--- | 47 |
| ebd:ECBD_1815 | QFIPW--- | 47 |
| ebe:B21_01784 | QFIPW--- | 47 |

| | | |
|---|---|---|
| ebl:ECD_01796 | QFIPW--- | 47 |
| ebr:ECB_01796 | QFIPW--- | 47 |
| eoc:CE10_2109 | QFIPW--- | 47 |
| eum:ECUMN_2119 | QFIPW--- | 47 |
| ecr:ECIAI1_1896 | QFIPW--- | 47 |
| ecy:ECSE_2000 | QFIPW--- | 47 |
| ecm:EcSMS35_1362 | QFIPW--- | 47 |
| ecx:EcHS_A1916 | QFIPW--- | 47 |
| ecoa:APECO78_12930 | QFIPW--- | 47 |
| eun:UMNK88_2297 | QFIPW--- | 47 |
| ecw:EcE24377A_2054 | QFIPW--- | 47 |
| elh:ETEC_1858 | QFIPW--- | 47 |
| eck:EC55989_2000 | QFIPW--- | 47 |
| esm:O3M_10790 | QFIPW--- | 47 |
| eso:O3O_14805 | QFIPW--- | 47 |
| esl:O3K_10820 | QFIPW--- | 47 |
| ecoh:ECRM13516_2234 | QFIPW--- | 47 |
| ecoo:ECRM13514_2331 | QFIPW--- | 47 |
| eoh:ECO103_2016 | QFIPW--- | 47 |
| eoj:ECO26_2596 | QFIPW--- | 47 |
| elx:CDCO157_2370 | QFIPW--- | 47 |
| etw:ECSP_2399 | QFIPW--- | 47 |
| ecf:ECH74115_2557 | QFIPW--- | 47 |
| ecs:ECs2536 | QFIPW--- | 47 |
| ece:Z2872 | QFIPW--- | 47 |
| ecok:ECMDS42_1500 | QFIPW--- | 47 |
| ebw:BWG_1639 | QFIPW--- | 47 |
| ecd:ECDH10B_1964 | QFIPW--- | 47 |
| ecj:JW1815 | QFIPW--- | 47 |
| eco:b1826 | QFIPW--- | 47 |
| ema:C1192_03175 | QFIPW--- | 47 |

- Gammaproteobacteria - Enterobacteria
  - Escherichia (61)
    - eco  Escherichia coli K-12 MG1655
    - ecj  Escherichia coli K-12 W3110
    - ecd  Escherichia coli K-12 DH10B
    - ebw  Escherichia coli K-12 BW2952
    - ecok  Escherichia coli K-12 MDS42
    - ece  Escherichia coli O157:H7 EDL933 (EHEC)
    - ecs  Escherichia coli O157:H7 Sakai (EHEC)
    - ecf  Escherichia coli O157:H7 EC4115 (EHEC)
    - etw  Escherichia coli O157:H7 TW14359 (EHEC)
    - elx  Escherichia coli O157:H7 Xuzhou21 (EHEC)
    - eoi  Escherichia coli O111:H- 11128 (EHEC)
    - eoj  Escherichia coli O26:H11 11368 (EHEC)
    - eoh  Escherichia coli O103:H2 12009 (EHEC)
    - ecoo  Escherichia coli O145:H28 RM13514 (EHEC)
    - ecoh  Escherichia coli O145:H28 RM13516 (EHEC)
    - esl  Escherichia coli O104:H4 2011C-3493 (EAEC)
    - eso  Escherichia coli O104:H4 2009EL-2071 (EAEC)
    - esm  Escherichia coli O104:H4 2009EL-2050 (EAEC)
    - eck  Escherichia coli 55989 (EAEC)
    - ecg  Escherichia coli O127:H6 E2348/69 (EPEC)
    - eok  Escherichia coli O55:H7 CB9615 (EPEC)
    - elr  Escherichia coli O55:H7 RM12579 (EPEC)
    - elh  Escherichia coli O78:H11:K80 H10407 (ETEC)
    - ecw  Escherichia coli O139:H28 E24377A (ETEC)
    - eun  Escherichia coli UMNK88 (ETEC, porcine)
    - ena  Escherichia coli NA114 (UPEC)
    - ecos  Escherichia coli O25b:K100:H4-ST131 EC958 (UPEC)
    - ecoa  Escherichia coli APEC O78 (APEC)
    - ecx  Escherichia coli O9 HS (commensal)
    - ecm  Escherichia coli SMS-3-5 (environmental)
    - ecy  Escherichia coli O152:H28 SE11 (commensal)
    - ecr  Escherichia coli O8 IAI1 (commensal)
    - ecq  Escherichia coli O81 ED1a (commensal)
    - eum  Escherichia coli O17:K52:H18 UMN026 (ExPEC)
    - eoc  Escherichia coli O7:K1 CE10
    - ebr  Escherichia coli B REL606
    - ebl  Escherichia coli BL21(DE3)
    - ebe  Escherichia coli BL21(DE3)
    - ebd  Escherichia coli BL21-Gold(DE3)pLysS AG
    - eci  Escherichia coli O18:K1:H7 UTI89 (UPEC)
    - eih  Escherichia coli O18:K1:H7 IHE3034 (ExPEC)

ecz  Escherichia coli O45:K1:H7 S88 (ExPEC)
ecc  Escherichia coli O6:K2:H1 CFT073 (UPEC)
eln  Escherichia coli O83:H1 NRG 857C (AIEC)
ese  Escherichia coli O150:H5 SE15 (commensal)
ecl  Escherichia coli ATCC 8739
ekf  Escherichia coli KO11FL
edh  Escherichia coli DH1
edj  Escherichia coli DH1
elu  Escherichia coli UM146
elw  Escherichia coli W
ell  Escherichia coli W
elc  Escherichia coli clone D i14
eld  Escherichia coli clone D i2
elp  Escherichia coli P12b
elf  Escherichia coli LF82
ecol  Escherichia coli LY180
ecoj  Escherichia coli JJ1886
efe  Escherichia fergusonii
ema  Escherichia marmotae
esz  Escherichia sp. E4742

▼ Salmonella (51)

sty  Salmonella enterica subsp. enterica serovar Typhi CT18
stt  Salmonella enterica subsp. enterica serovar Typhi Ty2
sex  Salmonella enterica subsp. enterica serovar Typhi P-stx-12
sent  Salmonella enterica subsp. enterica serovar Typhi Ty21a
stm  Salmonella enterica subsp. enterica serovar Typhimurium LT2
seo  Salmonella enterica subsp. enterica serovar Typhimurium 14028S
sey  Salmonella enterica subsp. enterica serovar Typhimurium SL1344
sem  Salmonella enterica subsp. enterica serovar Typhimurium T000240
sej  Salmonella enterica subsp. enterica serovar Typhimurium UK-1
seb  Salmonella enterica subsp. enterica serovar Typhimurium ST4/74
sef  Salmonella enterica subsp. enterica serovar Typhimurium 798
setu  Salmonella enterica subsp. enterica serovar Typhimurium U288
setc  Salmonella enterica subsp. enterica serovar Typhimurium var. 5- CFSAN001921
senr  Salmonella enterica subsp. enterica serovar Typhimurium DT2
seni  Salmonella enterica subsp. enterica serovar Typhimurium 138736
seen  Salmonella enterica subsp. enterica serovar 4,[5],12:i:- str. 08-1736
spt  Salmonella enterica subsp. enterica serovar Paratyphi A ATCC9150
sek  Salmonella enterica subsp. enterica serovar Paratyphi A AKU12601
spq  Salmonella enterica subsp. enterica serovar Paratyphi B
sei  Salmonella enterica subsp. enterica serovar Paratyphi C
sec  Salmonella enterica subsp. enterica serovar Choleraesuis
seh  Salmonella enterica subsp. enterica serovar Heidelberg SL476

shb  Salmonella enterica subsp. enterica serovar Heidelberg B182
senh  Salmonella enterica subsp. enterica serovar Heidelberg CFSAN002069
seeh  Salmonella enterica subsp. enterica serovar Heidelberg 41578
see  Salmonella enterica subsp. enterica serovar Newport SL254
senn  Salmonella enterica subsp. enterica serovar Newport USMARC-S3124.1
sew  Salmonella enterica subsp. enterica serovar Schwarzengrund
sea  Salmonella enterica subsp. enterica serovar Agona SL483
sens  Salmonella enterica subsp. enterica serovar Agona 24249
sed  Salmonella enterica subsp. enterica serovar Dublin
seg  Salmonella enterica subsp. enterica serovar Gallinarum 287/91
sel  Salmonella enterica subsp. enterica serovar Gallinarum/pullorum RKS5078
sega  Salmonella enterica subsp. enterica serovar Gallinarum/pullorum CDC1983-67
set  Salmonella enterica subsp. enterica serovar Enteritidis P125109
sena  Salmonella enterica subsp. enterica serovar Enteritidis EC20090135
seno  Salmonella enterica subsp. enterica serovar Enteritidis EC20090193
senv  Salmonella enterica subsp. enterica serovar Enteritidis EC20090332
senq  Salmonella enterica subsp. enterica serovar Enteritidis EC20090531
senl  Salmonella enterica subsp. enterica serovar Enteritidis OLF-SE1-1019-1
senj  Salmonella enterica subsp. enterica serovar Javiana
seec  Salmonella enterica subsp. enterica serovar Cubana
seeb  Salmonella enterica subsp. enterica serovar Bareilly
seep  Salmonella enterica subsp. enterica serovar Pullorum
senb  Salmonella enterica subsp. enterica serovar Bovismorbificans
sene  Salmonella enterica subsp. enterica serovar Thompson
senc  Salmonella enterica subsp. enterica serovar Tennessee
ses  Salmonella enterica subsp. arizonae
sbz  Salmonella bongori N268-08
sbv  Salmonella bongori serovar 48:z41:--
salz  Salmonella sp. SSDFZ69

▼ Shigella (12)

sfl  Shigella flexneri 301 (serotype 2a)
sfx  Shigella flexneri 2457T (serotype 2a)
sfv  Shigella flexneri 8401 (serotype 5b)
sfe  Shigella flexneri 2002017 (serotype Fxv)
sfn  Shigella flexneri 2003036
sfs  Shigella flexneri Shi06HN006 (serotype Yv)
ssn  Shigella sonnei Ss046
sbo  Shigella boydii Sb227 (serotype 4)
sbc  Shigella boydii CDC 3083-94 (serotype 18)
sdy  Shigella dysenteriae Sd197
sdz  Shigella dysenteriae 1617 (serotype 1)
shq  Shigella sp. PAMC 28760

▼ Enterobacter (27)

enc Enterobacter cloacae subsp. cloacae ATCC 13047
eno Enterobacter cloacae subsp. cloacae ENHKU01
enl Enterobacter cloacae subsp. dissolvens SDM
eclg Enterobacter cloacae GGT036
ecle Enterobacter cloacae ECNIH2
ecln Enterobacter cloacae ECNIH4
ecli Enterobacter cloacae ECNIH5
eclx Enterobacter hormaechei subsp. xiangfangensis
ecly Enterobacter hormaechei subsp. steigerwaltii
eclz Enterobacter hormaechei subsp. hormaechei
ehm Enterobacter hormaechei CAV1176
exf Enterobacter hormaechei subsp. xiangfangensis
ecla Enterobacter hormaechei subsp. hoffmannii ECNIH3
eclc Enterobacter hormaechei subsp. hoffmannii ECR091
eau Enterobacter asburiae L1
ekb Enterobacter kobei
eec Enterobacter ludwigii EcWSU1
elg Enterobacter ludwigii EN-119
ecan Enterobacter cancerogenus
ern Enterobacter roggenkampii
esh Enterobacter sichuanensis
ent Enterobacter sp. 638
eas Enterobacter soli
enx Enterobacter sp. E20
enf Enterobacter sp. FY-07
ebg Enterobacter bugandensis
end Enterobacter sp. ODB01

▼ Cronobacter (10)

esa Cronobacter sakazakii ATCC BAA-894
csz Cronobacter sakazakii Sp291
csj Cronobacter sakazakii ATCC 29544
ccon Cronobacter condimenti
cdm Cronobacter dublinensis
csi Cronobacter malonaticus CMCC45402
cmj Cronobacter malonaticus LMG 23826
cui Cronobacter universalis
cmw Cronobacter muytjensii
ctu Cronobacter turicensis

▼ Klebsiella (21)

kpu Klebsiella pneumoniae subsp. pneumoniae NTUH-K2044 (serotype K1)
kpp Klebsiella pneumoniae subsp. pneumoniae 1084 (serotype K1)
kpt Klebsiella pneumoniae subsp. pneumoniae ATCC 43816 KPPR1
kpe Klebsiella pneumoniae 342

kpo Klebsiella pneumoniae KCTC 2242
kpi Klebsiella pneumoniae CG43
kva Klebsiella variicola At-22
kpk Klebsiella variicola KP5-1
kvd Klebsiella variicola DX120E
kvq Klebsiella variicola DSM 15968
kox Klebsiella michiganensis KCTC 1686
koe Klebsiella michiganensis E718
koy Klebsiella michiganensis HKOPL1
kom Klebsiella michiganensis M1
kok Klebsiella oxytoca KONIH1
koc Klebsiella oxytoca CAV1374
eae Klebsiella aerogenes KCTC 2190
ear Klebsiella aerogenes EA1509E
kqv Klebsiella quasivariicola
kll Klebsiella sp. LTGPAF-6F
klw Klebsiella huaxiensis

▼ Citrobacter (15)
cro Citrobacter rodentium
cko Citrobacter koseri
cfd Citrobacter freundii
cbra Citrobacter braakii
cwe Citrobacter werkmanii
cyo Citrobacter youngae
cpot Citrobacter portucalensis
cfq Citrobacter freundii complex sp. CFNIH3
cama Citrobacter amalonaticus Y19
caf Citrobacter amalonaticus FDAARGOS_165
cif Citrobacter sp. FDAARGOS_156
cfar Citrobacter farmeri
cir Citrobacter sp. CFNIH10
cie Citrobacter sp. CRE-46
cpar Citrobacter pasteurii

▼ Gibbsiella (1)
gqu Gibbsiella quercinecans

▼ Raoultella (4)
ror Raoultella ornithinolytica B6
ron Raoultella ornithinolytica S12
rpln Raoultella planticola
rao Raoultella sp. X13

▼ Cedecea (1)
clap Cedecea lapagei

▼ Pluralibacter (3)

- pge Pluralibacter gergoviae
- esc Enterobacter lignolyticus SCF1
- kle Enterobacter lignolyticus G5

▼ Kosakonia (4)
- kor Kosakonia oryzae
- krd Kosakonia radicincitans
- kco Kosakonia cowanii
- kot Kosakonia sp. CCTCC M2018092

▼ Kluyvera (1)
- kgo Kluyvera georgiana

▼ Leclercia (6)
- lax Leclercia adecarboxylata
- lei Leclercia sp. LSNIH1
- leh Leclercia sp. LSNIH3
- lee Leclercia sp. W17
- ler Leclercia sp. 1106151
- lea Leclercia sp. J807

▼ Lelliottia (4)
- laz Lelliottia amnigena
- lef Lelliottia jeotgali
- lni Lelliottia nimipressuralis
- lew Lelliottia sp. WB101

▼ Buttiauxella (1)
- buf Buttiauxella sp. 3AFRM03

▼ Atlantibacter (1)
- ahn Atlantibacter hermannii

▼ Izhakiella (1)
- izh Izhakiella sp. KSNA2

▼ Yokenella (1)
- yre Yokenella regensburgei

▼ Unclassified Enterobacteria (2)
- ebf Enterobacteriaceae bacterium FGI 57
- ebu Enterobacteriaceae bacterium S05

▼ Yersinia (7)
- ypk6 Yersinia pestis KIM6
- yey Yersinia enterocolitica subsp. palearctica Y11 (serotype:0:3)
- yel Yersinia enterocolitica LC20
- yrb Yersinia ruckeri Big Creek 74 (serotype O2)
- yma Yersinia massiliensis
- yhi Yersinia hibernica
- yca Yersinia canariae

▼ Serratia (10)

smar Serratia marcescens SM39
smac Serratia marcescens subsp. marcescens Db11
spe Serratia proteamaculans
smaf Serratia sp. FGI94
serf Serratia sp. FS14
sers Serratia sp. SCBI
sfw Serratia fonticola DSM 4576
sfg Serratia fonticola GS2
serm Serratia sp. MYb239
squ Serratia quinivorans

▼ Rahnella (2)

raa Rahnella aquatilis HX2
rox Rahnella sp. ERMR1:05

▼ Sodalis (2)

sod Sodalis praecaptivus
pes Candidatus Sodalis pierantonius

▼ Photorhabdus (2)

plu Photorhabdus laumondii subsp. laumondii TTO1
pay Photorhabdus asymbiotica

▼ Providencia (4)

psi Providencia stuartii MRSN 2154
psb Providencia stuartii ATCC 25827
prq Providencia sp. WCHPHu000369
pvc Providencia vermicola

▼ Proteus (1)

ppe Proteus penneri ATCC 35198
